# Topological mixing and irreversibility in animal chromosome evolution

**DOI:** 10.1101/2024.07.29.605683

**Authors:** Darrin T. Schultz, Arno Blümel, Dalila Destanović, Fatih Sarigol, Oleg Simakov

## Abstract

Animal chromosomes can persist with recognizable homology over hundreds of millions of years, in spite of homology-obfuscating processes such as chromosomal fusion and translocation. The frequency and pace of these major genome structural changes are unknown, and it remains unclear whether or how they impact long-term genome evolution. Here, we compare whole chromosomal sequences of 3,631 genomes from 2,291 species spanning all major animal clades and show that animal karyotypes evolve primarily via karyotype contraction, associated with increased rates of chromosomal fusion-with-mixing and dispersion that largely obey chromosomal algebra^1^, or karyotype expansion, which breaks up ancestral linkage groups and forms new chromosomal elements via non-algebraic changes. We show that chromosomal changes can be associated with major extinction events. Using a multi-scale encoding of pan-animal genome homology and a manifold representation of genomic changes, we find that genome evolution is not only driven by changes at the chromosomal level, but that subchromosomal mixing and irreversibility define clade-specific evolution. Using this ‘evolutionary genome topology’ approach, we calculate extrema of irreversible genomic configurations and identify species that occupy intermediate manifold positions, providing evidence for distinct macro-evolutionary trajectories. We propose that investigation of mixed state accumulation around important gene loci (such as Hox) will be crucial in capturing and further study of clade-specific regulatory innovations.

## Introduction

Animal chromosomes have remained remarkably conserved since the last common animal ancestor^1–3^ that existed over 700 million years ago^4^. Some of the ancient animal chromosomal homologies date back even further to the ancestor of the Filozoa 804 million to 1.18 billion years ago^4^, as animal chromosomes share synteny with pieces of chromosomes^1,3^, or whole chromosome arms^3^ in their unicellular relatives (e.g., the filasterean amoeba *Capsaspora*). These portions of chromosomes that are conserved over time across many species are called ancestral linkage groups^5^ (ALGs), or chromosomal elements^6^. Many animal chromosomes can be explained by four algebraic^1^ combinations of ancestral chromosomal elements. A key property of chromosomal algebra is fusion-with-mixing (FWM), a statistically irreversible process where genes on a fused chromosome mix, and can be used as strong phylogenetic characters^1,3^. However, the prevalence of chromosomal algebra and long-term (macro-)evolutionary effects caused by these chromosomal changes^1^, and the changes in the selective landscape they create remain poorly understood.

Inferring algebraic changes in chromosome evolution relies on detectable chromosomal homology, which is eroded over time through chromosomal rearrangements like translocations. Inter-chromosomal translocations are considered to be rare events as a portion of gametes produced after translocation are unbalanced and selected against^7^. Therefore, organisms that reproduce sexually or have not passed through substantial population bottlenecks are expected to retain their ancestral chromosomal complement, while asexually reproducing organisms may avoid this selection and more quickly fix translocations. Quantification of chromosomal translocations (**Supplementary Information 1**, *dispersal*) across different animal clades, however, has been hampered by the lack of chromosome-scale assemblies. The only metazoan baseline estimate for gene dispersal out of chromosomes is 1% per 40 million years^1^. However, whether and how this rate is different among clades or chromosomal element sizes has not been studied. Only now, with the increased availability of chromosome-scale genomes, are we in a position to study and quantify the evolutionary history of chromosomal homology by measuring dispersion in extant organisms’ genomes.

On top of these algebraic descriptions, several transitions in animal karyotypes have been reported that do not fit clearly within the framework of the four algebraic chromosomal changes. In this manuscript, we refer to these as non-algebraic changes, where fissions and/or large inter-chromosomal translocations break the chromosomal element homology. In particular, there are animal clades in which the ancestral animal chromosomal elements have partly or wholly dispersed or rearranged beyond recognizability. Clades with such rearrangements include the genome reshuffling in cephalopods^8^, the derived karyotype of ctenophores^3^, the ‘Merian elements’ of lepidopterans^9^, the ‘Nigon elements’ of nematodes^10^. Clades with rearrangements beyond recognition include the ‘Muller elements’ of drosophilids^6,11^, and clitellate annelids^12^. Whether and how these transitions can still be explained by the core algebraic processes involving complete ancestral chromosomal elements, or whether a non-algebraic breakage of chromosomes is required, remains to be investigated. In either case, these departures from ancestral animal chromosomal elements, some of which occur rapidly on short evolutionary branches^12^, can result in what we would recognize as new sets of clade-specific chromosomal elements. Whether and how these saltatory, rapid transitions are followed by stasis or further algebraic or non-algebraic changes remains unclear.

The presence of algebraic and non-algebraic processes as two major possibilities to change genomes poses the question of whether and how such chromosomal rearrangements have any long-term impact on the evolution of functions such as gene regulation. No global functional enrichment in any chromosomal element has been reported^1,3^. While this suggests that chromosome-level changes could be explained as a random process, chromosomal fusion-with-mixing may have a fundamental and barely studied outcome: by both expanding the chromosomal coordinate system and enabling new combinations of genes to enter regulatory interactions, it may drive phenotypic innovations^13^ over the long-term.

There are known regulatory constraints on local genome structure, including co-expression^14^ and co-regulation^15^and it has been suggested that genome-wide changes in chromosomal composition may help drive^16^ or strengthen^17^ novel genomic interactions. The functional importance of the conservation and change in the animal genome architecture among different animal clades remains a nascent research topic due to the limited gene regulatory information for many of these clades. However, it is clear that ancient local gene linkages can be conserved as functional regulatory units^15,18^ and that chromosomal changes can^13^ and do^19^ facilitate the evolution of novel regulatory linkages and phenotypic novelties^19–21^.

Lastly, while theoretical models have suggested that large-scale genomic changes could be correlated with extinction and speciation^22^, there is little evidence connecting these patterns over macro-evolutionary time^23^. Furthermore, there is no explanation of how changes in species diversity may be reflected in the genomic record, including the alleged^24^ 62 million year^25,26^ periodicity of species extinction and origination. Understanding what type, if any, of chromosomal rearrangements are associated with speciation or extinction, and correlating these changes with patterns seen in the paleontological record, are key steps toward understanding the forces inherent in chromosome evolution. New methods are needed to resolve the outstanding question of the degree to which chromosomal rearrangements cause, or are consequences of, speciation and extinction.

Using a broad sampling of 3,631 metazoan chromosome-scale genomes, we address these questions by identifying general trends of algebraic and non-algebraic changes. Determining the time scale and relationships among large chromosomal changes and the evolution of new regulatory linkages is key to understanding the evolutionary potential of animal chromosomes and the emergence of novelties (**Fig. 1**). In this manuscript, we quantify this time scale, and characterize the macro-evolutionary impact of large chromosomal changes. We find that these large chromosomal changes are correlated with species origination and extinction in different clades, and find that cycles in chromosomal rearrangement corroborate cycles in species diversity from the fossil record. We generalize the existing algebraic approach to involve both chromosomal and sub-chromosomal scales, governed by meiotic and regulatory constraints, by applying manifold (topological) approaches to study genome evolution. Our results show that each clade can be characterized by its own set of ‘mixed’ chromosomal and sub-chromosomal characters. This approach augments fusion-with-mixing, which is a process at the chromosomal level, and reveals a similar process of mixing and entanglement at the sub-chromosomal level. Our findings suggest that animal genome evolution to be a stepwise process of accumulation of irreversibly mixed states at different levels of genomic organization. We term this approach ‘evolutionary genome topology’, and propose that irreversible mixing is the driving force behind the observed macro-evolutionary changes, or modes, of genome evolution.

**Fig. 1.**
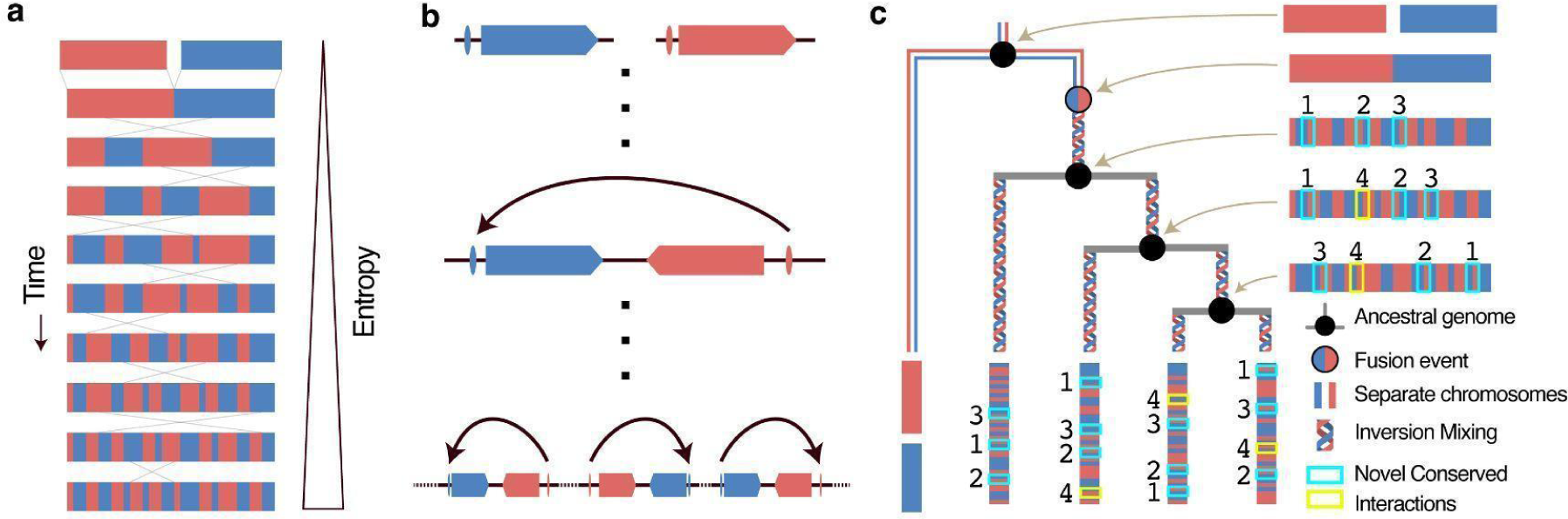
Fusion with mixing (FWM) at chromosomal and sub-chromosomal levels. **a**, Chromosome fusion-with-mixing (FWM) is a process that happens over long evolutionary time scales and leads to mixing of chromosomal regions previously on two separate chromosomes. The mixing over time can be measured as a form of entropy. **b**, Through FWM previously separate chromosome regions (top) can form new regulatory interactions not possible in the original separate chromosomes (middle). Over time many novel regulatory interactions can arise (bottom) and become constrained (which we refer to as entangled) through complex regulatory interactions. **c**, The FWM process is a sum of chromosomal-level mixing of gene sets and emergence of novel regulatory constrained linkages, novel regulatory ‘mixings’ can arise long after the original chromosomal fusions and therefore are synapomorphic local linkages.

## Results and Discussion

### Algebraic representation and completeness of metazoan genomes

We first sought to construct an overview of animal genome representation in the context of the Bilaterian-Cnidarian-Sponge ancestral linkage groups (BCnS ALGs^1^, **Supplementary Information 1**). Using available chromosome-scale genomes for 2291 metazoan species (**Methods** - *Constructing a genome database*), we inferred orthology with BCnS chromosomal elements (**Methods** - *Identifying BCnS ALGs*), as well as the BCnS chromosomal element persistence, fusions, and fissions (**Methods** - *Inferring ALG evolutionary history*). Among 406 possible combinations of BCnS ALG fusion pairs, we were able to detect 397 as either fused (●) or fused-and-mixed (⊗) in at least one species.

First considering subclades of the Metazoa, the Cnidaria are known for their well-preserved BCnS ALG chromosomal complement^1^. For a set of 17 chromosome-scale cnidarian genomes (n=17 species), all 29 BCnS ALGs were detected in at least one species, totaling 107 out of 406 possible ALG fusion combinations, meaning that 299 ALG fusion combinations did not appear in any of the 17 cnidarian species. Of the 107 possible ALG fusion combinations, 46 (33% of 107) were species-specific. These estimates of fusion represent a lower limit, as these counts can be impacted by species with poorly assembled chromosomes, or chromosomes that have rearranged to the point where certain ALGs are dispersed across chromosomes. In nematodes, a clade known for its genome contraction and proposed rearrangement-rich history^10^, 28 ALGs were detected in at least one of 37 species. However, the ALG composition varied strongly between nematode species. The highest number of retained ALGs was 27 in a single species, the median was 19 ALGs per species, compared to 28 in cnidarians. Similarly, in strong contrast to cnidarians, 5 nematode species completely lacked recognizable ALGs. In nematodes, 101 out of total 378 possible fusions were present, 34 of which were species-specific. Of these 101 fusions in nematodes, 76 were fusions also seen in at least one cnidarian genome, predictably so given the convergence expected from the low karyotype number in nematodes. These data point to different levels of ALG retention across metazoans, but interestingly also highlight a similar level of algebraic combination space exploration, and thus algebraic representation, of nematodes and cnidarians.

Across our dataset we found only nine ALG combinations that have not been fused in any of the species. These unobserved fusions include the combinations A1a⊗/●Qb, B1⊗/●Qb, B1⊗/●Qc, C2⊗/●G, J1⊗/●Qb, O1⊗/●Qb, O1⊗/●Qc, O2⊗/●R. We note that the BCnS ALGs R, Qb, and Qc are among the smallest BCnS ALGs with 24, 12, and 14 constitutive orthologs. Our ability to detect such fusions is affected by dispersion of genes to non-homologous chromosomes.

### The time scale of chromosomal element homology dispersal

We next characterized dispersal, a process in which inter-chromosomal translocations cause genes to move from one ALG to other chromosomes. We investigated these properties by computing the dispersion state in the animal genomes (n=3,631 genomes) in relation to the BCnS ALGs (**Methods** - *Dispersal characterization*). The average dispersion of genes relative to the BCnS ALG complement is as low as 0.2% for chromosomally highly preserved genomes like the scallop *Mimachlamys varia* (**Fig. 2b**, **Extended Data Fig. 1**), to between 53%-73% for genomes in the nematode genus *Caenorhabditis*, reflecting partial BCnS ALG loss, and 100% for species like *Drosophila* or *Anopheles* whose genomes are highly rearranged and the BCnS ALGs are no longer detectable (**Fig. 2**).

**Fig. 2.**
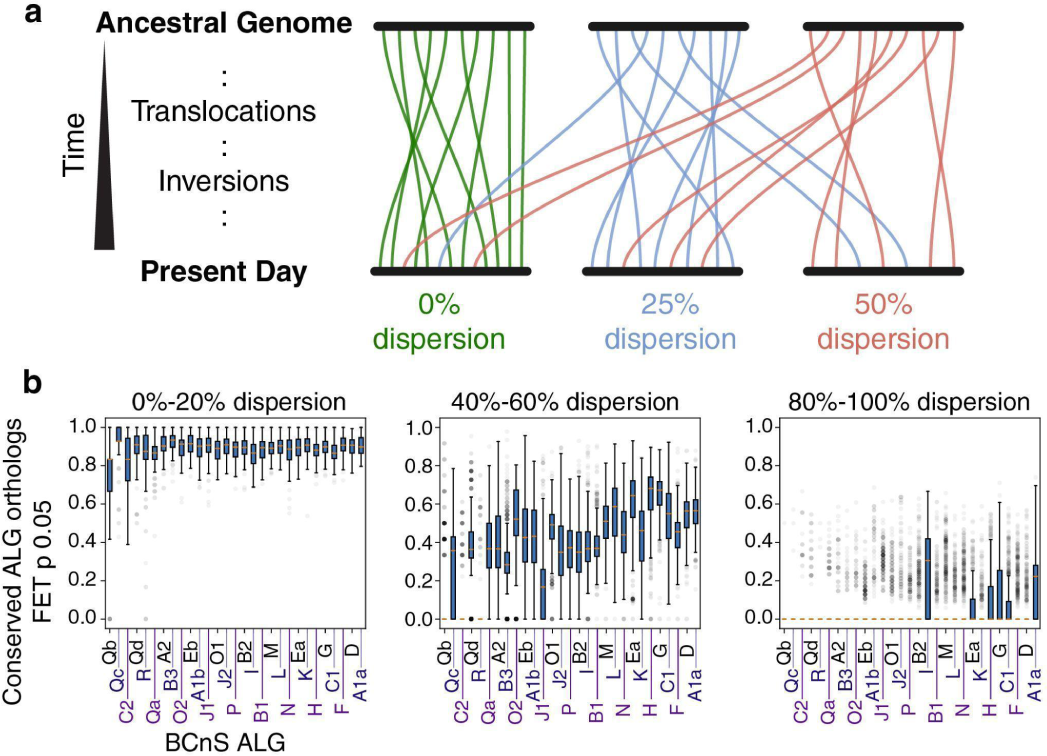
Dispersion of chromosomal elements is independent of their size. **a**, Dispersion is the process of loci leaving their chromosome of origin through translocations. Inversions over time cause loci that were previously on separate chromosomes to mix. **b**, Chromosome dispersion in animal genomes (n=3,631 genomes, 2,291 species) varies greatly depending on the evolutionary history of the species since the BCnS ancestor. Some species retain nearly 100% of BCnS ALGs as recognizable on one or more chromosomes, while some species retain no recognizable BCnS ALGs (*p* ≤ 0.05, Bonferroni-corrected one-sided Fisher’s exact test). Genomes are binned by the total percent of BCnS ALG dispersion across the entire genome. Individual BCnS ALG’s dispersion values for each genome are plotted for the boxplots. Box centerlines are the median values, box limits span from the first quartile to the third quartile, fliers span to 1.5 times the interquartile range (Q3-Q1), and outliers are plotted individually. See also **Extended Data Fig. 1** for *Pecten*-centered dispersion calculations.

The total dispersal scales with evolutionary time and the lineage in question. For example, a comparison of the genomes from two closely related bivalve species exhibit very little dispersion. Comparing genomes of animal or unicellular species with a common ancestor at the Choanozoa, Filozoa, or Holozoa nodes (**Extended Data Fig. 1**) reveals that very few chromosomal elements are conserved.

We find that the number of dispersed orthologs is independent of the chromosomal element size (**Fig. 2**). This has several important implications. First it enables predicting the life span of a given chromosomal homology. For example, because larger chromosomes have more positions available for homology detection, their homology can remain detectable longer than that of the smaller chromosomes. The smallest chromosome of the scallop *Pecten maximus* (18 BCnS orthogroups in 32.5 Mbp, BCnS ALG C2), for example, is unrecognizable in species that diverged over 550 million years ago, while the homology with larger *Pecten* chromosomes remain detectable (**Fig. 2**). For these reasons, while even the smaller chromosomes are still likely to persist within animals, without special analyses^3^ the same chromosomal homologies are completely undetectable in animal-unicell comparisons. The second implication is that for smaller chromosomes, homology detection will rely either on a shorter evolutionary distance or, with larger evolutionary distance, will rely more on them having fused with other chromosomes. When a small chromosome fuses with another, the probability that its genes are dispersed is reduced by the ratio of summed lengths of the two original chromosomes versus the total genome size. One example is the bilaterian linkage group (BLG) Q element^1,27^, which became colocalized on the same chromosome in the ancestor of Chordates from four very small elements (Qa, Qb, Qc, Qd) that are also found colocalized in various combinations in other animal clades. Under this model, smaller ancestral elements are predictably undetectable in certain clades, a prominent example being BCnS ALG R, which is not present in chordate genomes^1^. Together, these data clarify the time decay of BCnS chromosomal element homology, and highlight the process of dispersal that happens along algebraic chromosome evolution.

### Uneven rates of chromosomal element evolution

Slow chromosomal evolution is a marked feature in animals^1,27^, and is characterized by the presence and preservation of the BCnS ALGs, each on a single or a small number of chromosomes. However, it is unclear how much the fusion and dispersal rates can vary. To assay the evolutionary history across the whole animal tree we developed a phylogeny-aware method of ALG fusion rate estimation (**Methods** - *ALG evolution simulations*, **Extended Data Fig. 2**) and applied it to our dataset (**Fig. 3a,b**). These results show similar fusion rates within each clade for any given BCNS ALG combination. We did not find a clear preference of smaller BCnS ALGs to undergo fusions. However, using a phylogenetically-aware method (**Methods**), we observed general discrepancies between fusion rates in various clades. For example, in sponges the fusion rate is 1.07 fusions/million years whereas in spiralians it is lower at 0.8825 fusions/million years (**Extended Data Fig. 2**).

**Fig. 3.**
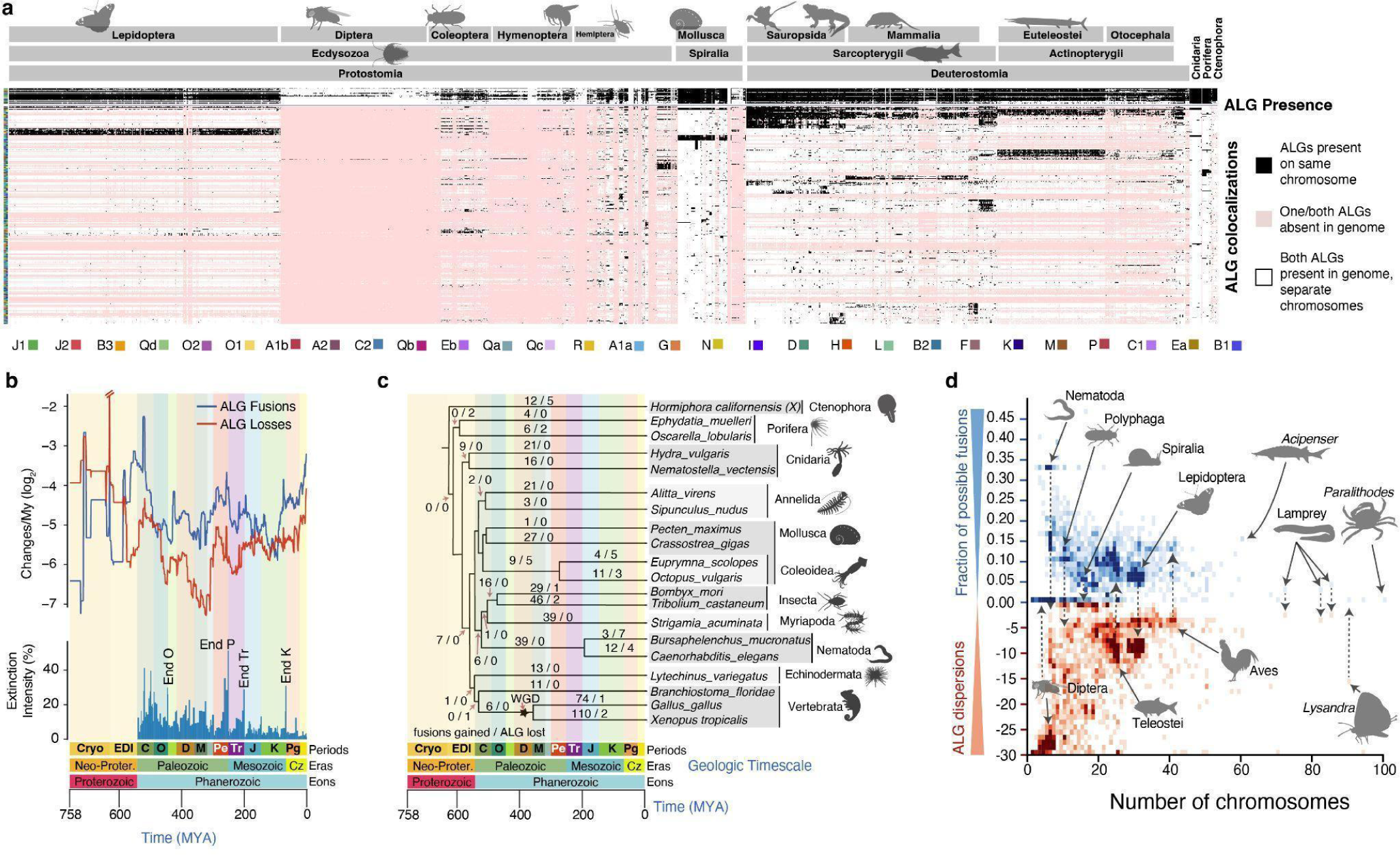
Global patterns of chromosomal fusions and fissions across metazoans. **a**, The grid of pink, black, and white pixels shows the BCnS ALG presence and colocalization in 2262 animal species. One column encodes one genome from each species. Black pixels in the top row, *ALG Presence*, shows BCnS ALG persistence in the extant genomes. The lower rows, labeled *ALG colocalizations*, depict pairs of ALG colocalizations on the same chromosome (black), or on separate chromosomes (white) are depicted. Pink indicates a lack of detectability due to a lost BCnS ALG homology. See **Extended Data Fig. 3** for a phylogenetic reconstruction of the extant genomes. **b,** Inferred rates of chromosomal changes on the animal tree in million-year bins. The period of most intense chromosomal changes occurred early during animal diversification. **c**, Counts of shared fusions and dispersions in major metazoan evolutionary transitions. **d**, Distribution of fusion and dispersion of BCnS linkage groups as a function of chromosome number (n=3,631 genomes).

Our results also show that chromosome evolution in animals did not occur at a steady rate, and that the processes of fusion and homology loss through dispersal vary in timing and intensity (**Fig. 3b**). For example, we find that the rate of BCnS ALG changes through both dispersal and fusion are highest in the branches leading to the last common ancestors of major extant clades (**Fig. 3b,c**). Notably, across all animals the averaged rate of BCnS ALG fusions increased during the end-Ediacaran 550 million years ago and peaked at the beginning of the Cambrian (approximately 1 fusion per 8 million years). The rate of chromosomal element fusions lowered by the end of the Cambrian period, and the rate varied (1 fusion per 32±16 million years) until the end of the Mesozoic, 66 million years ago. During this period, the rate of homology loss through dispersal was lower than the rate of fusions by a factor of two, with peaks of homology loss in the Cambrian (1 loss per 22 million years), Devonian (1 loss per 64 million years), and Permian (1 loss per 32 million years) periods.

Such cases of increased rates of chromosome rearrangements can result in observed saltatory changes along an evolutionary branch, in which the genomes of the extant species are highly rearranged relative to typical animal chromosomes with intact BCnS ALGs, and these changes are restricted to particular epochs. In some clades, for example the Panarthropoda, after an initial burst of fusions and losses at the beginning of the clade diversification during the Cambrian approximately 580 million years ago^28^ (1 fusion per 4 million years), these changes continued to happen at a rate twice as high as found averaged across animals (1 fusion and 1 loss per 16 My - **Extended Data Fig. 3d**). In other clades, like mosquitos or clitellate annelids, the changes appear to happen so rapidly that we did not observe remnants of transitionary genomic states in closely related species, and therefore can only estimate lower bounds of rates of change along these branches. For example, clitellate annelid genomes are dramatically rearranged relative to the genomes of their close polychaete relatives^12^, and none of the 29 ancestral BCnS ALGs are recognizable.

With the chromosomal evolutionary rates across the history of animal evolution in hand, we tested for a correlation between the observed rate of chromosomal changes and species origination or extinction^26^ (**Methods** - *Correlation inference with species diversity*). In one of the most species-rich groups in our dataset, the protostomes (n=1828 genomes), we found that chromosomal fusion rates are correlated with species extinction (one-tailed Spearman’s rank correlation coefficient, *r_s_* = 0. 53, *p* < 1 × 10^−40^, *n* = 542 million-year bins) and species origination rates (one-tailed Spearman’s rank correlation coefficient, *r_s_* = 0. 29, *p* < 1 × 10^−10^, *n* = 542 million-year bins). However, in vertebrates (n=1,598 genomes), both chromosomal fusion and loss rates are anti-correlated with species extinction (one-tailed Spearman’s rank correlation coefficient, BCnS ALG fusions *r_s_* =− 0. 29, *p* < 1 × 10^−11^, BCnS ALG losses *r_s_* =− 0. 51, *p* < 1 × 10^−37^, *n* = 542 million-year bins) and origination rates (one-tailed Spearman’s rank correlation coefficient, BCnS ALG fusions *r_s_* =− 0. 46, *p* < 1 × 10^−29^, BCnS ALG losses *r_s_* =− 0. 35, *p* < 1 × 10^−16^, *n* = 542 million-year bins) (**Extended Data Fig. 3e-n**). We find that several extinction events, such as the Permian–Triassic, Triassic–Jurassic, and the Cretaceous–Paleogene extinction events co-occur with spikes in both BCnS ALG fusions and losses across all animals (**Fig. 3b,c** and **Extended Data Fig. 3**).

Together these results suggest that protostome and vertebrate speciations, and the clades within them, had distinct underlying chromosomal evolution mechanisms that vary in intensity over time. In the case of protostomes, these results suggest that large-scale chromosomal changes that become fixed along an evolutionary branch may be the consequence of massive changes in selective constraints, like population bottlenecks caused by near-extinction events. In the case of vertebrates, on the contrary, our results suggest that higher rates of chromosomal change occurred in times of population stability. While counter-intuitive, this pattern could be expected for the rediploidization process that follows whole genome duplication, and therefore can happen in absence of any major extinction events^29^. Interestingly, we found evidence for periodicity in both BCnS ALG fusions and losses across animals (**Extended Data Fig. 4**, **Supplementary Information 2**), including a cycle of 62-64 million years in BCnS ALG fusions in Metazoa (Monte Carlo simulations, 93% support) similar to the proposed 62 million year cycle found in species origination and extinction events^25,26^ (**Methods** - *Chromosome evolution periodicity analysis*). Regardless of the observed extinction event co-occurrence, our data clearly indicate that fusion and dispersal rates vary with time and clade, but happen to the same degree across all BCnS ALGs (**Extended Data Fig. 3**). With this in mind, the divergent, clade-specific trends of changing rates of chromosome evolution are a new orthogonal dataset to better interpret patterns of speciation seen in the paleontological record.

### Algebraic and non-algebraic processes of chromosome evolution

The balancing act between dynamic rates of chromosomal fusions and translocations has had a substantial role in generating distinct genomic organizations in animals^1,30,31^. Using our pan-metazoan dataset, we next asked whether the observed chromosomal evolutionary patterns could be categorized according to their evolutionary mechanisms (**Methods** - *Inferring modes of animal evolution*). Because the ancestral myriazoan^3^ genome comprised between 20-30 chromosomes^1^, we reasoned that the chromosome evolution patterns of extant animals with fewer than 20 chromosomes would differ from the pattern found in extant animals with more than 30 chromosomes. In our dataset we find two mechanistically distinct modes of karyotype evolution (which we term consolidation and dissociation) that cause deviations from the ancestral myriazoan karyotype, and a ‘default’ mode of karyotypic stasis (**Fig. 3d**). In species with fewer than 10 chromosomes, we found that the predominant mechanism of karyotype reduction could be explained by extensive chromosomal fusions. Example clades with reduced karyotypes include: *Caenorhabditis* (n=28 genomes, 14 species), in which the BCnS ALGs have been fused and mixed, but are still recognizable on single chromosomes and can be expressed as algebraic BCnS ALG combinations; or *Drosophila* (n=216 genomes, 44 species) and the Culicidae (mosquitos, n=79 genomes, 36 species), in which the ancestral BCnS ALGs are completely dispersed across all chromosomes.

We found that the predominant pattern in animal genomes with few chromosomes is to have recognizable ALGs after fusions. This first mode of chromosome evolution, consolidation, is defined by the rate of chromosomal fusions being greater than the rate of chromosome fissions or translocations. We find that nematodes are archetypal of this evolutionary pattern. Despite nematodes having rapid sequence evolution rates^32,33^, and being suggested as having a new set of chromosomes^34^, nematode genomes (n=57 genomes, 37 species) can still be described by algebraic operations. In nematodes, BCnS ALGs are often retained as complete units, and only one BCnS homology loss (ALG C2) is shared among species in the clade (**Fig. 3d**). Similarly, we found that oyster (Ostreida, n=18 genomes, 9 species) chromosomes are the result of many fusions (43), but all nine species in our dataset still have all 29 BCnS ALGs recognizable and intact on single oyster chromosomes. As a consequence of the chromosomal fusions, the karyotype reduction during chromosome evolution causes genomes to be more dramatically affected by subsequent dispersal, resulting in the observed mode of evolution often associated with ‘faster evolving’ genomes.

The genomes that have substantially increased their chromosome counts since the ancestral myriazoan karyotype, on the other hand, could have done so through two processes. We call this second mode of evolution dissociation. In the first pattern of the dissociation mode, which is the non-algebraic type mentioned earlier, chromosomal fissions or translocations result in extant genomes in which BCnS ALGs are present on a large number of chromosomes. Furthermore, the BCnS ALGs are weakly or not dispersed, in contrast to the consolidation mode. The second pattern of the dissociation mode that could explain an expanded karyotype is whole genome duplications, which often result in secondary gene loss on the duplicated homeologous chromosomes, as seen in vertebrates^27,35,36^. While obeying different mechanisms, both dissociation scenarios result in similar syntenic patterns, stemming from the increase in karyotype number that can undergo additional fusions^8^. However, the nature of the non-algebraic fission/translocation process makes it impossible to represent these genomes in algebraic form, therefore a new ALG notation is required. Examples for these clades include coleoid cephalopods^13^ (**Fig. 3d**) and clitellate annelids^12^. Vertebrates, due to duplication of complete ancestral chromosomes, can fully be represented by whole-genome duplications and algebraic combinations^1,27^.

The causes of the fissions could be due to their holocentric nature^37^, centric fissions, or possibly due to chromothripsis^38^. Given that chromosomal fissions separate ALGs into unique combinations of ancestral ALGs, and the improbability of those loci coming together again^3^, non-algebraic chromosome evolution is therefore a major contributor to saltatory changes over short evolutionary time and may thus may be more strongly associated with periods of species origination and extinction than algebraic evolution.

Finally, our analysis also confirms the ‘default’ mode of animal chromosome evolution is a general karyotypic stasis with few intrachromosomal rearrangements, as previously described for several animal clades^1^. In the karyotypes of these species, centered around 20 chromosomes, there have been very few chromosomal fission or fusion events in the approximately 700 million years^4^ of evolution from the myriazoan ancestor.

Together, our data point to three predominant modes of chromosome evolution in animal genomes that have shaped animal genome evolution since divergence from the ancestral myriazoan genome. Due to the different timings and degrees of fusion/fission and dispersal, the processes of karyotype consolidation via extensive fusion and dissociation are inherently distinct, reflecting different meiotic constraints and population dynamics. Any species that has a consolidation or dissociation of the ancestral chromosome count is therefore the result of a lineage in which the balance of stasis shifted, irrevocably changing the substrate of future chromosomal evolutionary events. Importantly, our study provides evidence that there is no intermediate state or trajectory between the dissociation and consolidation modes. While over longer evolutionary times one mode could follow after another, such cases are extremely rare.

### Macro-evolutionary impact of chromosomal fusion processes

Both algebraic and non-algebraic processes bring together genes that previously existed on different chromosomal elements. As the functional advantage of distal relationships on single chromosomes is unknown^39^, we instead investigated the impact of fusion-with-mixing, and more generally chromosomal inversions, on the emergence of local subchromosomal linkages, which are far better understood and can confer coregulation, for example, the provenance of the closely linked gene clusters in bilaterians, such as Hox^17,40^ and many other local, or micro-syntenic, linkages^15,41^.

First, we sought to characterize the mixing time scales via a simple simulation model of inversions (**Methods** - *Simulation of fusion-with-mixing*) using beads-on-a-string genomes. We define τ_s_ as the number of inversion cycles required to reach a fully mixed, high-entropy state (**Extended Data Fig. 5**). We compared the τ_s_ of chromosomes of different sizes. We find that smaller chromosomes (fusion products of 100 genes) approach τ_s_ very quickly already after ∼50 inversion cycles. Expectedly, larger chromosomes (fusion products of 1000 genes) approach τ_s_ at 1000 inversions. Direct measurements from cephalopod genomes (n=4 species, **Methods** - *Inversion breakpoint inference)* show that inversions in this clade occur at a rate of 4.29-29.2 inversions per million years^42^, suggesting a timeframe of 34.2-233.1 million years to reach the highly entropic state, for a chromosome of 1000 genes. These results suggest that achieving an irreversible mixed state after chromosome fusion depends on the chromosome size, but can be relatively quick on the macro-evolutionary time scale especially for the smaller chromosomes.

As a proxy for local regulatory linkages, we next sought to measure how many novel neighborhoods can be explored using this chromosome inversion model (**Extended Data Fig. 5**, **Methods** - *Simulation of fusion-with-mixing*). We define a measure τ_50_ that quantifies how many inversion cycles are needed to explore 50% of all possible neighborhoods. These interaction neighborhoods are defined either as direct neighbors, or all genes within windows of 2, 5, or 10 genes. Such neighborhoods can reflect regulatory compartments of animal genomes, such as topological associating domains (TADs) in which multiple genes can be coregulated^43,44^. Similar to τ_s_, τ_50_ increases with chromosome sizes, yet the scale is vastly different. To reach 50% of direct neighbor interactions for a fusion product of 100 genes, approximately 1,700 inversion cycles are required on average (25 times more than for τ_s_) (n=10 trials). Similarly, for a fusion product of 1,000 genes, 173,000 inversion cycles are needed (130 times larger than the respective τ_s_) (n=10 trials, **Extended Data Fig. 5**). Through direct measurement (mixing value *m^3^*)of ancient fusion-with-mixing events in the scallop *Pecten maximus* we confirmed that entropic saturation through chromosomal inversions has not been reached yet. These results suggest that for an average metazoan chromosome that has been retained over 600 or more million years, the vast majority of interactions are still unexplored.

Persistent topological properties can substantially reduce τ_50_ by allowing distal gene interactions to be maintained. When such neighborhoods are taken into account, τ_50_ is substantially reduced. Exploring 50% of all possible neighbors in 5-gene neighborhoods reduces the τ_50_ time by 93.5% (n=10 trials). This finding is important as it suggests that presence of persistent topologically associating domains (TADs) allows for faster evolutionary exploration of the regulatory environment given that genomes with interactions at the scale of TADs will explore the available combinations faster than genomes dominated by cis-interactions that can interact only with their immediate neighbors. Our expectation is therefore that non-bilaterian animal genomes that have been suggested to lack maintained loop extrusion through CTCF or similar insulator factors^45,46^, would be slow in the exploration of their regulatory potential, as opposed to genomes with the ability to establish and maintain more distal regulatory properties.

The long-term nature of continuous emergence of novel local neighborhoods on old or newly formed chromosomal elements describes an evolutionary process on the macro-evolutionary time scale (**Fig. 1**). However, whether and how local neighborhoods can be considered an irreversible FWM-like process, and the probabilities of a new sub-chromosomal FWM event given the macro-syntenic background, remain unexplored. The degree of irreversibility of local gene linkage mixing can therefore be seen as the cost of losing that regulatory constraint, as is the case for the Hox clusters^17^ and many other micro-syntenic clusters^15,41,47^. While the functionalities of the majority of these clusters are still unexplored, their colocalization property implies a different regulatory constraint^15,18^. Therefore, these micro-syntenic clusters, if evolving under persistent regulatory constraint, will be ‘un-mixable’ into their original states. Our results on genome rearrangements (**Methods** - *Inversion Breakpoint Inference*) and previous work^48–50^ highlight that inversion breakpoints tend to happen at insulator regions, i.e., TAD boundaries, suggesting high selective constraint to keep genes and their regulatory landscape within compartments together.

Different genomic compositions, such as repetitiveness or the chromosome size and composition after algebraic and non-algebraic changes, impact how much regulatory novelty can still be evolutionarily explored within each chromosome. The larger chromosomes are more likely to still harbor unexplored neighborhoods even after hundreds of millions of years after chromosomal element fusion, whereas small chromosomal elements are more likely to exhaust their combinatorial potential. To quantify how much modern genomes can be represented by both chromosomal and sub-chromosomal mixed states, we next describe a topological approach to this problem.

### Topologically mixed states that define distinct animal genome evolutionary trajectories

Comparative genomics analyses often focus on a single genomic scale at a time. To study FWM and other chromosomal changes simultaneously it is necessary to implement all genomic scales and features in one framework^31^: from single genes, to micro-syntenies and small structural changes, to whole chromosome composition and organization. Each level evolves under certain meiotic, gene-regulatory, or other functional constraints. To study this, we implemented an ‘evolutionary genome topology’ approach, which, in mathematical terms, allows us to view genomes and FWM processes on chromosomes as topological operations.

In this manuscript, we calculate pairwise distances between BCnS orthologs, but this approach can be generalized to any homologous features in genomes, such as: genes, conserved non-coding elements (CNEs), or repeat families and their insertions (**Methods** - *Forms of genome topological analysis*). This allows us to implement two topological approaches. The first approach compares the topology of many genomes simultaneously, which we term ‘multi-genome topology’ (MGT) and allows the analyses of evolutionary trajectories across species. The second approach, which we call ‘multi-locus topology’ (MLT), allows the simultaneous analysis of the topology evolution of loci within and between clades of genomes. In the MLT analyses, the interaction values for pairs of homologous loci are phylogenetically averaged, while the MGT uses distance measures from individual genomes (**Methods** - *Evolutionary topology implementation*). Dimensionality reduction and projection methods, such as uniform manifold approximation and projection (UMAP)^51^ or t-SNE^52^, are then used to visualize the interaction matrix of whole-chromosome differences between genomes, or the within-genome topology of single or multiple genomes (**Methods** - *Evolutionary topology implementation*). Eigenvector and singular value decomposition approaches are the key concepts for FWM generalization across genomic scales of organization, as they can be used as a measure of mixing (**Supplementary Information 3**). Raw orthology information and positional data can be stored and processed with graph database approaches (**Methods** - *Graph database integration*, **Extended Data Fig. 6**) for the downstream dimensionality and manifold projection approaches (**Fig. 4**, **Extended Data Fig. 7**, **Extended Data Fig. 8**).

**Fig. 4.**
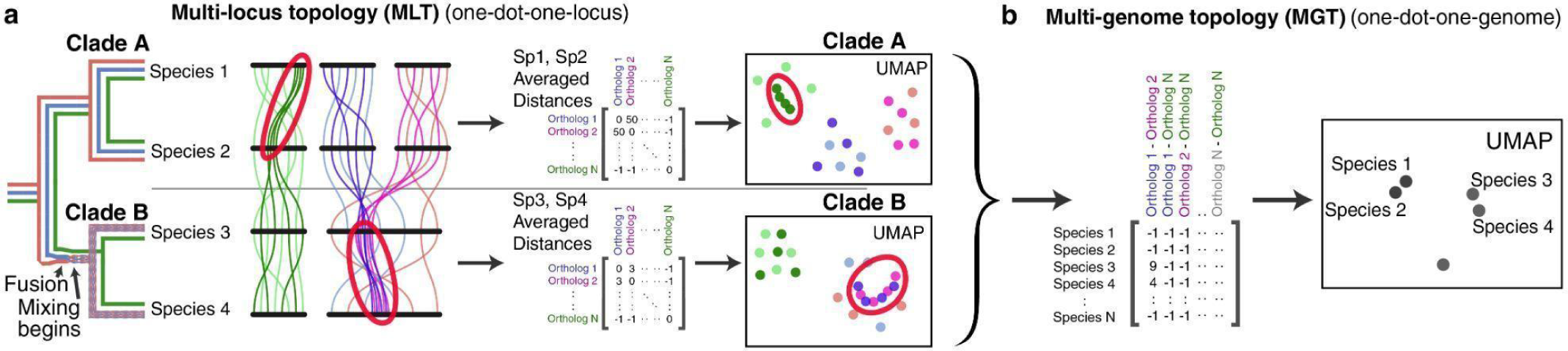
FWM representation on topological spaces. **a**, Mixing dynamics and gene colocalization can be approximated by averaged pairwise distances between orthogroups in each genome for a given clade to produce manifold projects for each clade. This distance matrix is then parsed through a manifold reduction algorithm like UMAP to visualize orthologous regions that are more prone to mixing. Specifically, FWM of chromosomes will result in their orthogroups to be inseparable on the UMAP. Similar observations can be made for mixed sub-chromosomal regions (red oval in Clade A UMAP). **b**, Orthogroup pair distances across species can further be used to identify species with similar colocalization distributions.

The multi-genome topology UMAP (**Fig. 5a**) reflects key genomic groupings. The genomes that are least derived from the ancestral BCnS state, mainly mollusc and cnidarian genomes, form a cluster of species on the UMAP (**Extended Data Fig. 9a-d**). The central hub of genomes is composed of species that have some chromosome rearrangements (**Extended Data Fig. 9e**), but are not dramatically rearranged from the ancestral animal genome. The derived states where the ancestral chromosomal element composition has been most heavily modified are found in the species clouds located at the edges of the plot (**Extended Data Fig. 9f,g,i**). These distinct clouds are connected via intermediate genomic configurations, suggesting that some genomes’ configurations are in less-derived states and are still on the way to the advanced differentiation in terms of their BCnS ALG composition and mixing. For example, the genomes of a tick (*Dermacentor silvarum*), a beetle (*Tribolium castaneum*), mosquitoes and fruit flies can be linked by many intermediate genomes, forming a trajectory (**Extended Data Fig. 9h**). These analyses recapitulate the two major outcomes (modes) of chromosome evolution: consolidation and dissociation. However, these states can now be refined and seen as being composed of multiple very distinct evolutionary trajectories (**Fig. 5**, **Extended Data Fig. 10b**).

**Fig. 5.**
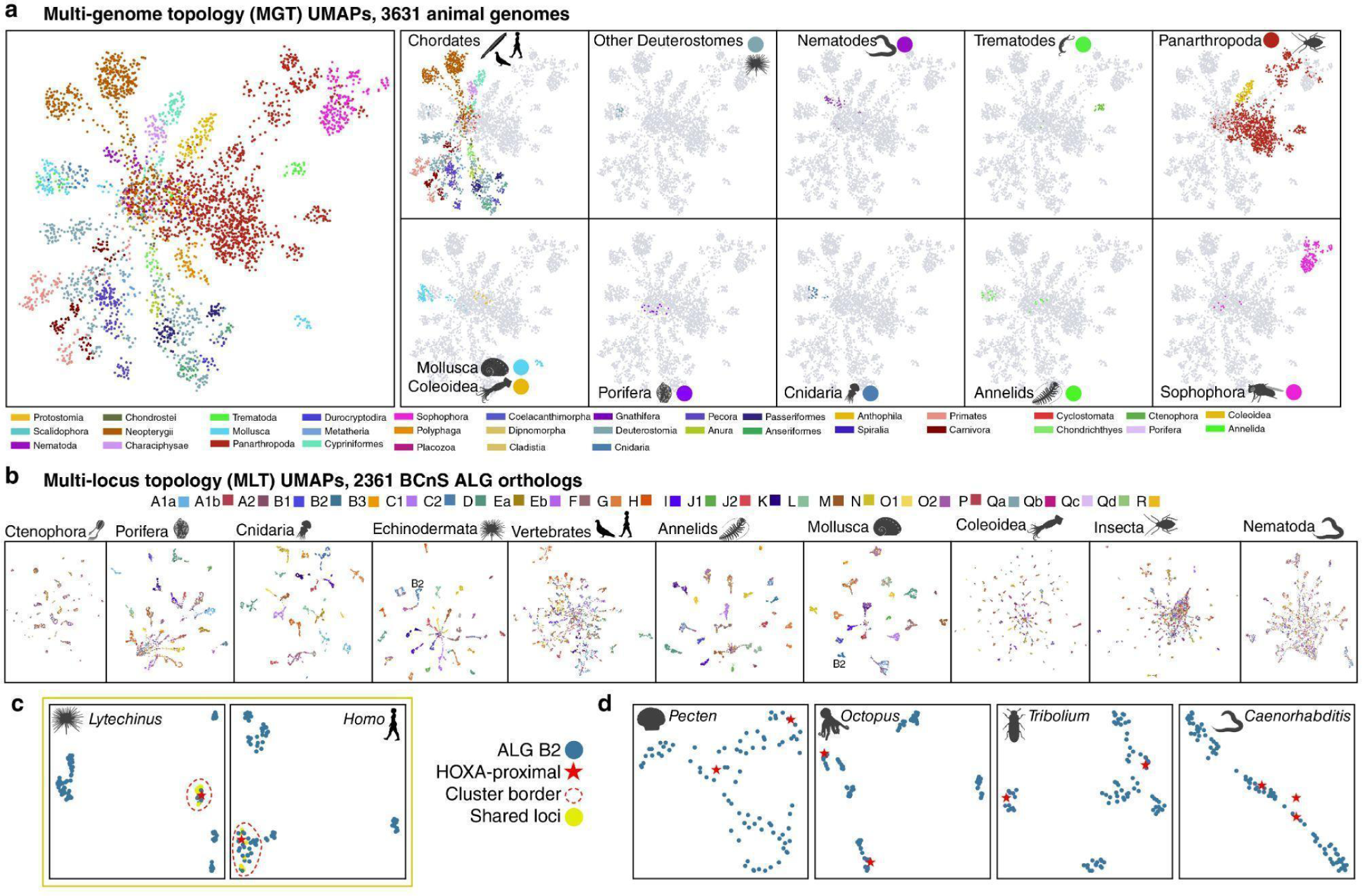
Evolutionary topology of chromosomal elements. **a**, A multi-genome topology (MGT) projection of 3631 chromosome-scale animal genomes (n= 2,291 species), in which each dot represents one genome. We use the genes from the previously-defined BCnS ALGs as anchors in the genomes of all chromosome-scale genomes, showing distinct colocalizations that are often clade specific. **b**, Multi-locus topology (MLT) projections for individual clades reflects evolutionary properties of chromosomes. One dot represents one orthologous protein shared across animals (n=2,361 BCnS ALGs). **c**, Sub-chromosomal topology around the Hox cluster orthologs for BCnS ALG B2 genes allows the identification of a set of co-mixed orthologs. **d**, Chromosomal-level topology around the Hox cluster in several animal groups.

The different trajectories within even closely related species are exemplified by applying the multi-genome topology approach to individual animal clades (**Extended Data Fig. 10**). Closely related species, even after tens of millions of years of divergence, occupy close positions in the UMAP space (**Extended Data Fig. 10a**). True oysters (n=18 genomes, 9 species), which have a reduced karyotype of 1n=10, form a cluster distant from the other mollusc genomes (n=108 genomes, 80 species), other clades with unique paths to karyotype reduction (limpets: n=6 genomes, 5 species, karyotype 1n=9, scaphopods: n = 2 genomes, 2 species, karyotype 1n=9), additionally form distinct clusters separate from one another. Similarly, the major clades in Diptera (n=509 genomes, 275 species), form clusters in UMAP space (**Extended Data Fig. 10b**). We recently introduced the use of macrosyntenic FWM to resolve ancient, recalcitrant phylogenetic nodes^3^, and others have used microsynteny similarity to resolve more recent evolutionary relationships^53^. The MGT approach, because it simultaneously incorporates both chromosomal and sub-chromosomal localization data, has clear potential to be applied to resolve outstanding phylogenetic questions.

Our findings from the MGT approach suggests that irreversible chromosomal changes preclude a direct evolutionary path between trajectories, and more importantly, back to the ancestral state. We hypothesized that accumulation of irreversible topologically mixed states defines the directionality of this macro-evolutionary process.

To test the degree and stability of mixing at the sub-chromosomal level, we investigated the evolutionary genome topology within two or more genomes in specific clades using the MLT approach. To do this we averaged, with phylogenetic weighting (**Methods** - *Evolutionary topology implementation*), the linear distance matrices of species within various clades and computed the sub-chromosomal, whole-genome (**Fig. 5b**) and individual chromosomal UMAPs (**Fig. 5c**). The homologous chromosomes form clearly distinct regions, which are separate entities in the manifold projection. Within each chromosome, local feature clusters emerge as well. While the maximum resolution of our analysis is the 2367 BCnS ALGs present mapped to the genome, these features reflect local conserved micro-syntenies, enhancer-promoter interactions, and other interactions which can be further explored at the limits of orthology detection.

The MLT manifold representation also recovers known FWM events, such as the A2⊗N and A1b⊗B3 in cnidarians (**Fig. 5b -** Cnidaria) relative to clades without these fusions, such as non-cephalopod molluscs (**Fig. 5b -** Mollusca). Similarly, the manifold representation can be used to investigate the degree of FWM. For example, the dots comprising the cnidarian A2⊗N ancestral FWM are highly mixed on the manifold representation (**Fig. 5b -** Cnidaria), while the poor mixing between B2 and Ea⊗Eb iN echiNoderms (N=21 geNomes, 18 species) is also clear (**Fig. 5b -** Echinodermata). Similar to chromosome-level FWM events, we can also use local clustering of features on the manifold, or their connectivity, to estimate the degree of local FWM. For highly preserved micro-syntenic linkages, such as Hox, (**Fig. 5c**), their unlinking would imply substantially increasing the distances of their individual components on the manifold. For example, we find that the posterior Hox genes tend to be located in a separate topological cluster in protostomes (**Fig. 5d**).

The evolutionary topology approach also identifies the presence of loci in clade-specific linkages^54^. These linked loci remain similarly positioned, not necessarily adjacent, relative to one another in 2D or 3D space against the backdrop of inter- and intra-chromosomal changes. Our approach provides sets of mixed states that define each clade (**Methods**). Pairs of loci can define the archetypal genome for a clade, and are topologically linked, if (1) the pairs are unique to this clade - that is these two sequences are only present on the same chromosome within genomes in this clade, (2) the pairs are fixed at a stable distance in this clade, regardless of whether they are close or distant, or (3) the distances between the pairs are very small in genomes in this clade (*cis-* interactions). It is therefore possible that type 3 pairs (small inter-locus distance, or *cis*-) pairs are also type 2 pairs (stable inter-locus distance), but not vice versa. Type 3 pairs can generally be considered to reflect widely studied micro-syntenic regions, or constraints that act on very short (*cis-*) distances. Such regions can be easily identified via comparisons of both chromosome- and non chromosome-scale genomes^15,41,47^. Type 2 pairs, on the other hand, can only be uncovered by averaging dozens to hundreds of chromosome-scale genomes. Type 2 pairs therefore may reflect more distal entanglement(**Supplementary Information 1**). Extensive mixing, leading to entangled pairs, of many gene enhancer-promoter regulatory links within a defined genomic region (e.g., a topological domain) can be seen to make the domain ‘resistant’ to random translocations that would break one or more of such regulatory links.

There are generally few locus pairs unique to any given animal clade (type 1 locus pairs), reflecting the degree of dispersion and convergent colocalizations during metazoan evolution. On the contrary, there are many locus pairs that are characteristically stable (type 2) or close (type 3) in any given clade. For example, in Spiralia there are 2,398 pairs stable in the clade, and 2,471 especially close pairs. While the test for type 1 pairs is binary, the stringency of our search for type 2 and type 3 pairs (one-sided μ-2σ cutoffs, ≥ 50% of species with this pair, **Methods**) mean that these numbers are conservative estimates of the number of identifiable markers. We note that the most strongly stable pairs (type 2) are also close pairs (type 3), and these novel linkages appear to be rarely lost in genomes within the clade once the loci become linked (**Extended Data Fig. 11c,f**). These results suggest that within-chromosomal mixing and constraint, rather than karyotype-level changes alone, are dominant forces in creating new clade-specific chromosomal landscapes. Moreover, the presence of clade-specific stable pairs suggests that novel sub-chromosomal linkages have a degree of irreversibility, and changes in them appear to be selected against.

Our approach reveals emergent patterns in stable pair formation that reflect different evolutionary histories of chromosomes. For example, BCnS ALG B2 that contains the Hox cluster and BCnS ALG A1 that contains the pharyngeal cluster^55^ as well as many homeobox genes^27^ show contrasting patterns between deuterostomes and spiralians (**Extended Data Fig. 11b,c,e,f)**. In spiralians, both chromosomes are characterized exclusively by the accumulation of within-chromosomal stable pairs (BCnS ALGs B2 or A1, **Extended Data Fig. 11b,e,f)**. With A1 forming almost 5 times the number of novel, spiralian-specific, stable pairs. In contrast, in deuterostomes, also inter-ALG pairings can be observed to contribute to both B2 and A1, with BCnS ALG B2 particularly enriched in novel inter-ALG pairings. This highlights different levels of neighborhood exploration and evolution of putatively novel regulatory constraints within these ancient chromosomal elements.

In summary, our evolutionary topology approach, currently based on pre-computed BCnS ortholog distances, can be used to place genomes on the manifold, identify their macro-evolutionary trajectory, and to identify evolutionarily mixed chromosomal and sub-chromosomal linkages. This analysis supports the view that saltatory changes in genomes (such as accumulation of algebraic FWMs or non-algebraic transitions) put genomes on distinct and irreversible evolutionary trajectories. Our results describe genome evolution as an iterative process with each step consisting of one or several algebraic or non-algebraic events that lead to the irreversible, iterative erasure of ancestral states, with strong intra-chromosomal entanglement. We show that this process can be described in a mathematical topological context that reflects and predicts extensive evolutionary constraints that act at whole-chromosomal and regulatory linkage levels.

## Conclusions

In this paper, we describe and quantify evolutionary time scales of the processes that govern changes in chromosomal and sub-chromosomal homologies, and connect these patterns of genomic change to patterns of speciation and diversity seen in the paleontological record. As new chromosome-scale data is emerging for many phylogenetically informative clades, together with genome topological and regulatory data, we can determine the evolutionary trajectories that are set and defined by the present-day configurations of animal genomes. We propose that using the evolutionary genome topology approach, as described in this manuscript, will be crucial in understanding the evolution of mixing, loci linkage, and irreversibility across genomic scales. Our data reveals thousands of clade-specific stable sub-chromosomal configurations that can be used as synapomorphic characters in phylogenetic studies. We propose that the irreversibility of such mixed states is the main contributing factor to the observed genomic macro-evolutionary trends. Moreover, our data provides insights into the potential, at the macro-evolutionary time-scale, to evolve new regulatory linkages within available chromosomal elements. This approach also highlights the role of saltatory changes in chromosomal complement reorganization, via both consolidation and dissociation means, to increased innovation through exploration of novel regulatory interactions that are predicted in this manuscript to be explored over the span of hundreds of millions of years. Understanding the degree of and the mechanisms underlying their predicted irreversibility with functional studies in various metazoan model organisms will provide further evaluation to this hypothesis.

## Extended Data Figures

**Extended Data Fig. 1.**
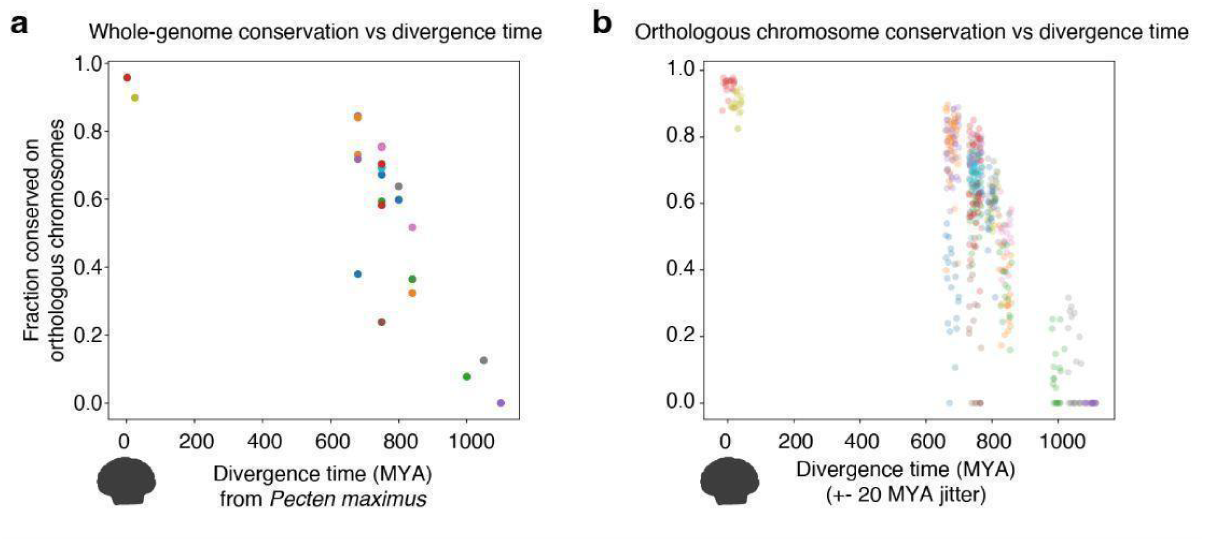
Time dependence of BCnS ALG dispersal. **a**, We compared the chromosome conservation of the whole *Pecten maximus* genome to the genomes of other organisms (n=22 species), as distantly related as the filasterean amoeba *Capsaspora owczarzaki*. The y-axis shows the total fraction of orthologs conserved in the genome. The decay of homology is a time dependent process, with a rate of slowed change between present day and 600 million years ago. **b**, The same data, but the conservation value of individual *Pecten* chromosomes relative to the other species are shown.

**Extended Data Fig. 2.**
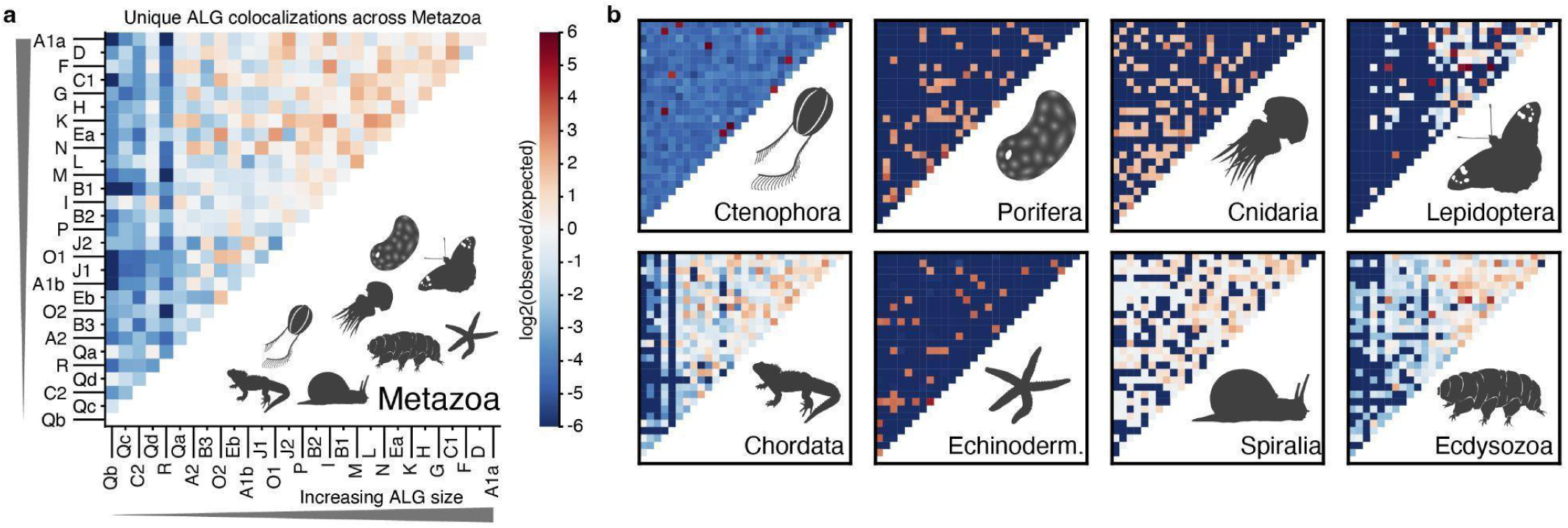
Phylogeny-aware counting of BCnS linkage group fusions. **a**, A general overview across metazoan genomes (n=3,631 genomes) shows depletion of smaller BCnS ALG fusions. **b**, BCnS fusion rates are different among animal clades and is a function of both chromosomal number within each clade and dispersion rate (see also **Fig. 3**).

**Extended Data Fig. 3.**
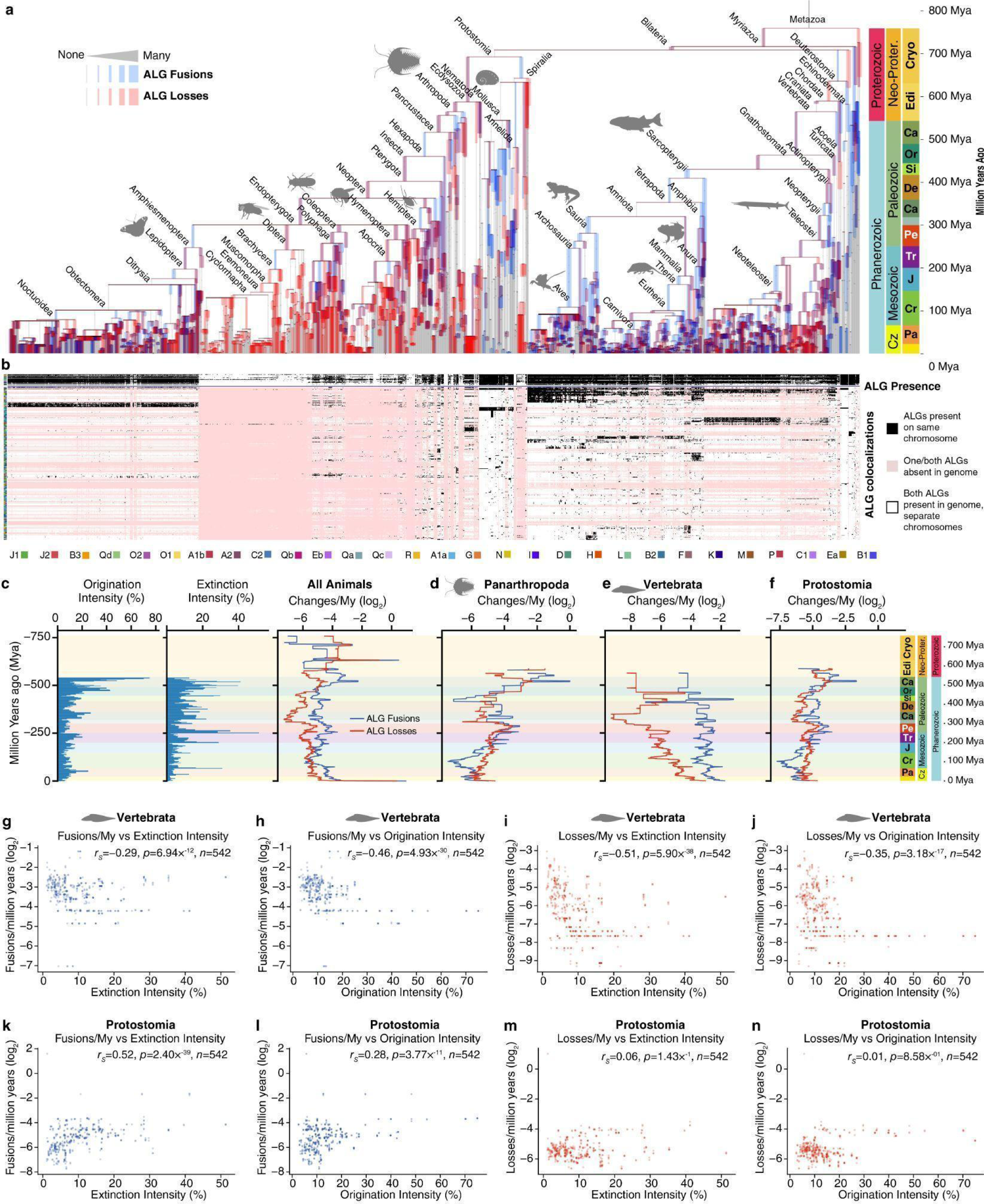
BCnS linkage profiling across metazoan chromosome-scale genomes. **a**, Chromosome evolutionary history encoded along the phylogenetic tree of 2262 species. BCnS ALG losses are red, and BCnS ALG fusions are blue. **b**, The grid encoded in **Fig. 3a**, where each column corresponds to a tree tip in panel a. Black pixels in the top row, *ALG Presence*, shows BCnS ALG persistence in the extant genomes. The lower rows, labeled *ALG colocalizations*, depict pairs of ALG colocalizations on the same chromosome (black), or on separate chromosomes (white) are depicted. Pink indicates a lack of detectability due to a lost BCnS ALG homology. **c**, Unedited panel shown in **Fig. 3b**, which shows the changes per million years across the evolutionary tree shown in panel a. **d**, The changes per million years in the Panarthropoda, **e**, Vertebrata, and **f**, Protostomia. **g**-**j**, plots of changes per million years versus the species origination or extinction intensity for the changes within Vertebrata. Statistics reported in the panels are one-tailed Spearman’s ranked correlation coefficients. **k-n**, the same plots as panels g-j for changes within Protostomia.

**Extended Data Fig. 4.**
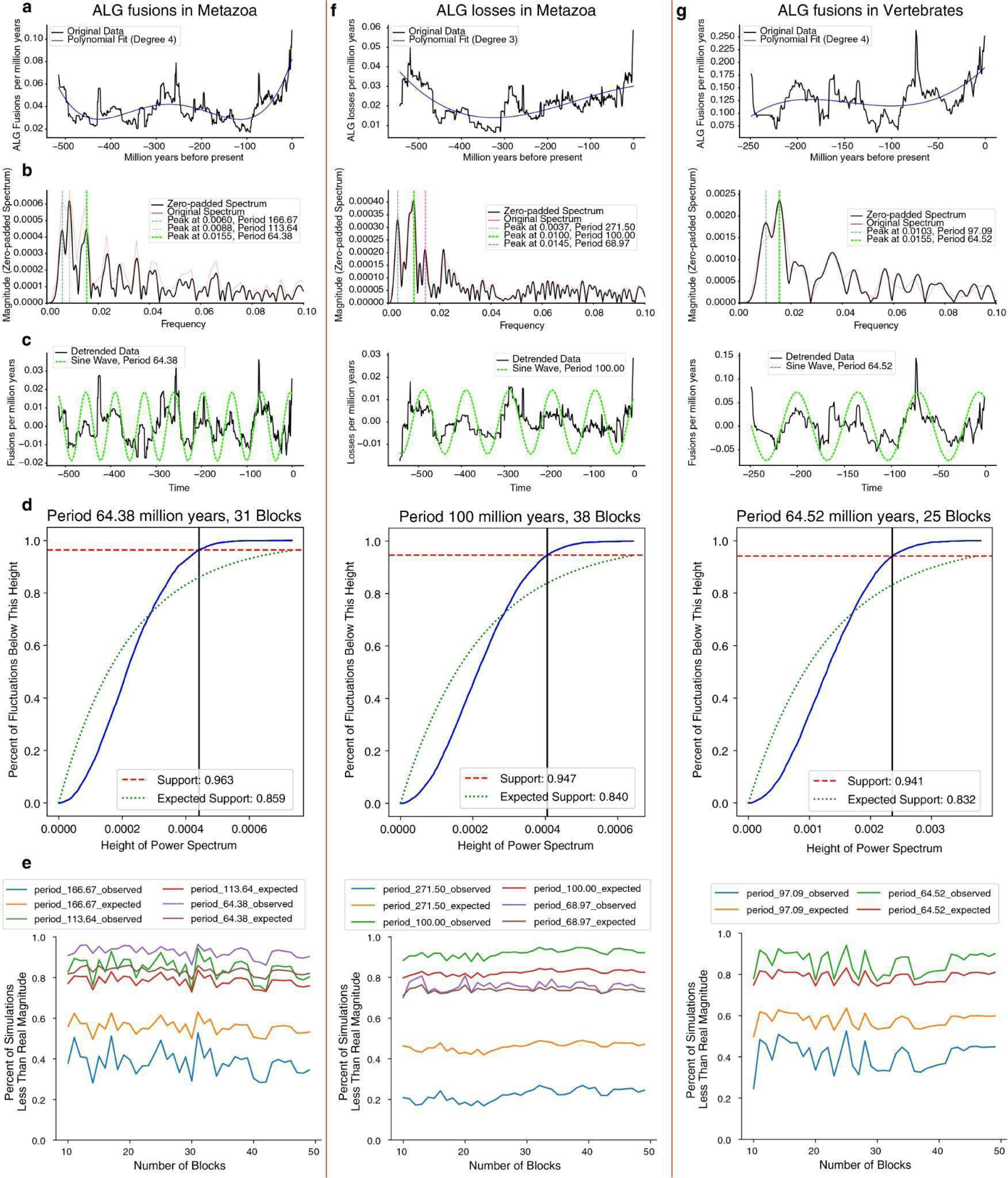
Periodicity analysis of BCnS ALG fusion or loss events. **a**, Rates of BCnS ALG fusions across the Metazoa in million-year bins, fitted with a quartic function. **b**, The power spectrum of the zero-padded discrete Fourier transform of the detrended dataset (black line) and the transformed unpadded power spectrum (red line). The three strongest peaks are highlighted with vertical dotted lines (periods of 166.67 million years, 113.64 million years, and 64.38 million years). **c**, The detrended dataset overlaid with a sine wave of a 64.38 million-year period. **d**, Example results from one Monte Carlo simulation for the 64.38 million year period. The datasets transitions were split into 31 blocks and shuffled. The green dotted line represents the exponential expectation function based on the average magnitudes in the power spectrum from the Monte Carlo trials. The vertical black line indicates the observed peak magnitude. The red support value is the percentage of simulated Fourier transforms with magnitudes less than the observed peak’s magnitude. **e**, The effect of the number of blocks on the observed support for the peak from the Monte Carlo simulations. **f**, Similar analysis as panels a-e for BCnS ALG losses in metazoans. **g**, Similar analysis as panels a-e for BCnS ALG fusions in Vertebrates.

**Extended Data Fig. 5.**
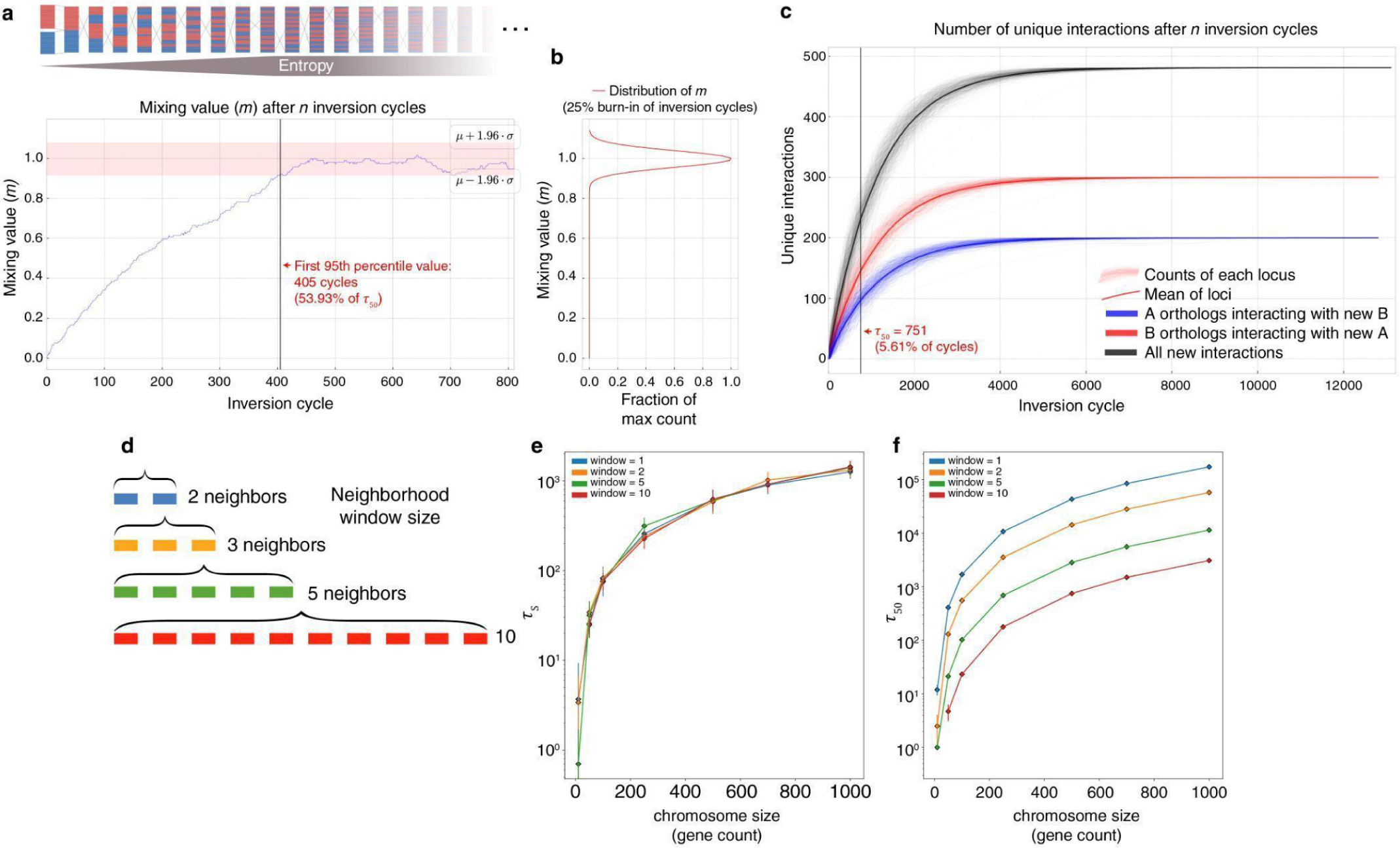
Time scale of mixing and genomic neighborhood exploration. **a**, Trace of the mixing value *m* (**Methods**) for two beads-on-a-string chromosomes after fusion. This simulation is for 300 genes in group A fusing with 200 genes in group B. **b**, The mixing value *m* reaches the mass of the distribution (μ-1.96σ < *m* < μ+1.96σ of *m* after 25% burn-in) of a well mixed state after 405 inversion cycles. **c**, Counting the number of unique direct (nearest neighbor) interactions or pairs during the simulation. For the gene and window parameters of panel a, half of all possible neighborhoods have been explored only after 751 inversion cycles. **d**, The window size parameter changes the window size of genes that can be considered for new interactions. Small number of neighbors n may best model genomes with cis-regulation, while a large number of neighbors may best model genomes with gene regulation grouped by TADs. **e**, Window size has little effect on the amount of time it takes for genes to fully explore the number of possible combinations. Error bars are the standard deviations of 10 simulations. **f**, More practically, however, the time to explore half or more of the combinations is reduced exponentially as the window size increases.

**Extended Data Fig. 6.**
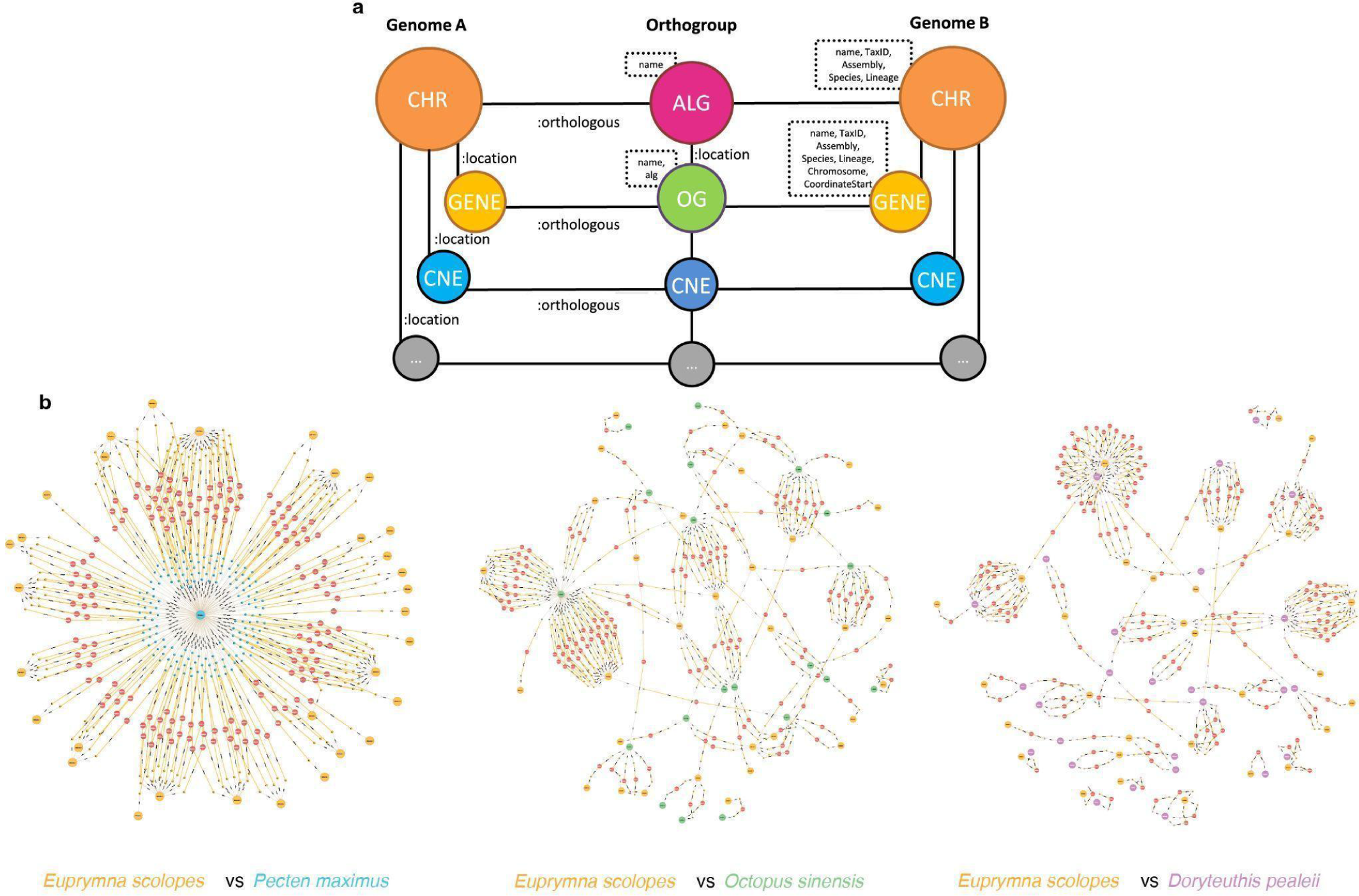
Graph database representation of orthologous connections between chromosomes, genes, and noncoding elements. **a**, data structure as implemented in the Neo4j database. Properties of different node types are shown inside dashed lines. **b**, example networks for different species groups highlight simpler (one-to-one chromosomal, with some gene dispersion) relationships between two squid species (Euprymna and Doryteuthis), and fission of the ancestral linkage groups (as represented by a Pecten chromosome) in Euprymna. Orthogroup nodes belonging to BCnS ALG A1a are in red color in middle size, large nodes are chromosomes, small nodes are genes, and nodes of each species are colored as in the species names shown.

**Extended Data Fig. 7.**
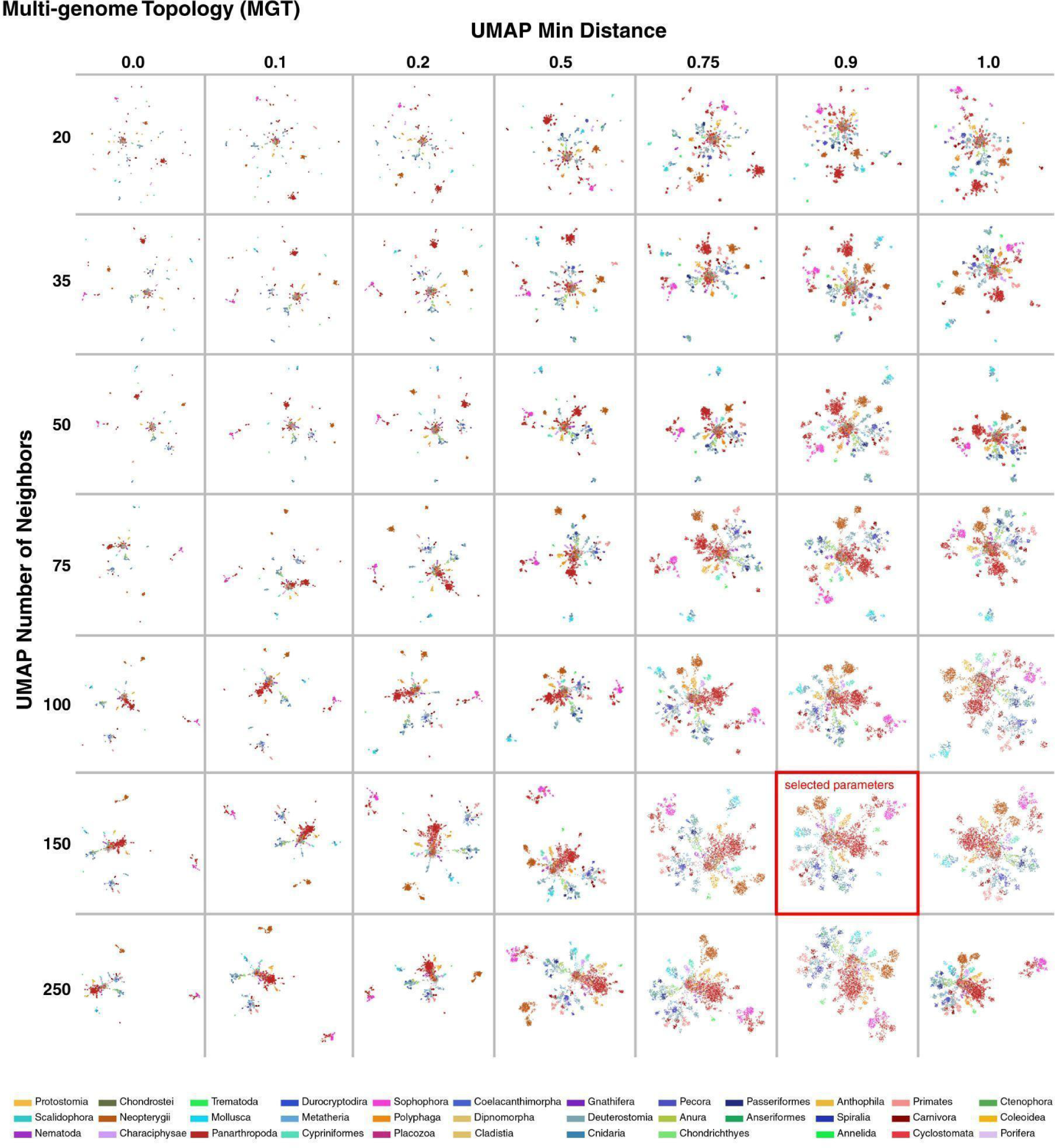
UMAP parameter space exploration. While local distribution of species on UMAPs change according to neighborhood and min distance parameters, the overall clade assignment and composition remains stable.

**Extended Data Fig. 8.**
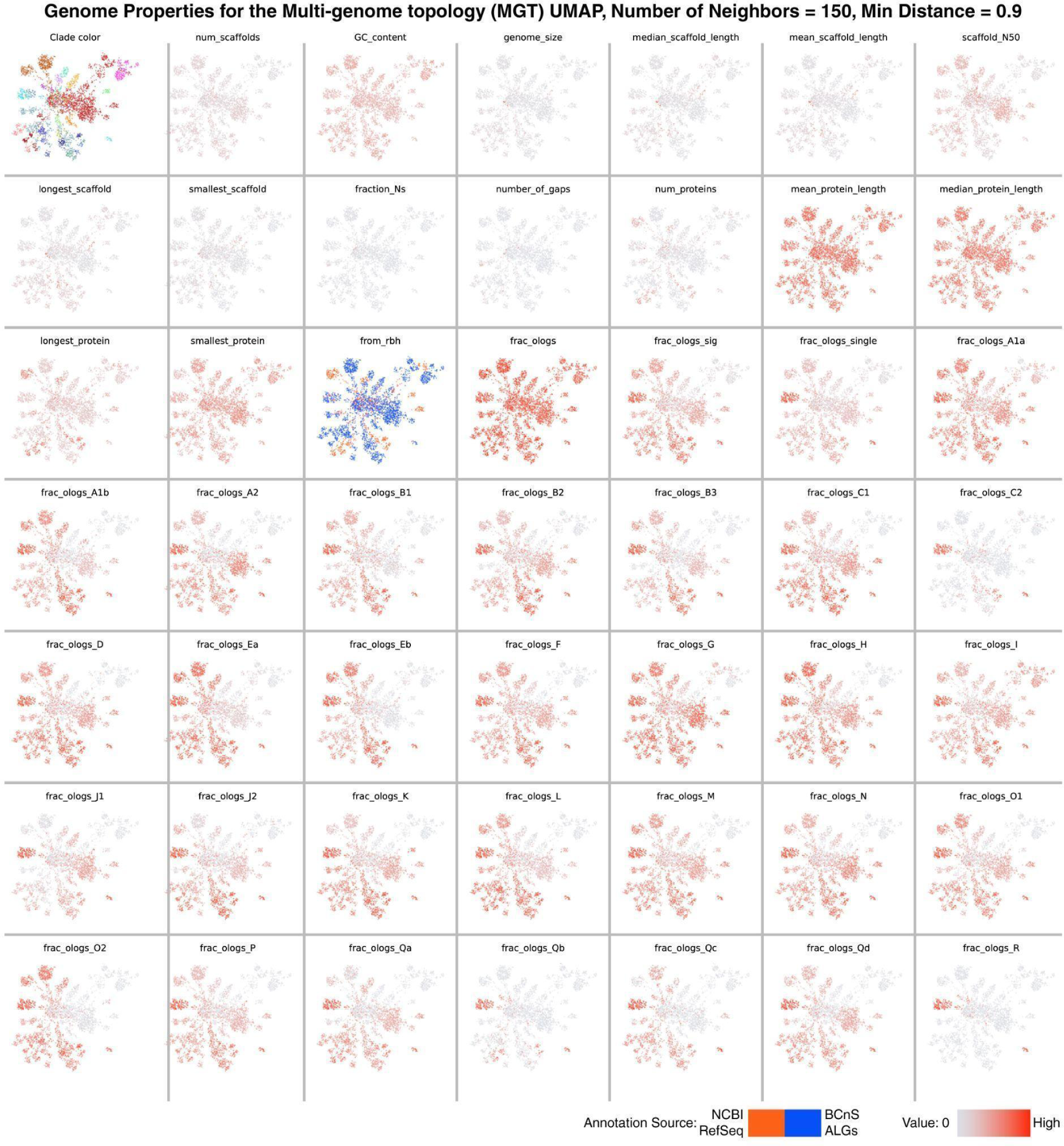
Genomic property localization on UMAP species embedding. The UMAP visualization of genomes shows localized evolutionary history of basic genome properties and BCnS ALG changes. For each subplot, the scale is varied depending on the range of the values of all the genomes.The fields of the figure are: *clade_color*, each dot is colored by the clade code from **Extended Data Fig. 7**; *num_scaffolds*, the number of scaffolds in the genome assembly fasta file; *GC_content*, the fraction of G or C nucleotides in the genome assembly fasta file; *genome_size*, the number of basepairs in the genome assembly fasta file; *median_scaffold_length*, the median length (basepairs) of all of the scaffolds in the genome assembly fasta file; *mean_scaffold_length*, same as the previous field, but the mean; *scaffold_N50*, the N50 of all the scaffolds in the genome assembly fasta file; *longest_scaffold*, the longest scaffold (basepairs) in the genome assembly fasta file; *fraction_Ns*, the number of Ns in the genome assembly fasta file divided by the total number of bases in the fasta file; *number_of_gaps*, The number of gaps, or continuous stretch of at least 10 N characters, in the genome assembly fasta file; *num_proteins*, the number of proteins present in the .chrom file for this genome; *mean_protein_length*, the mean length (amino acids) of the proteins present in the protein fasta file for this genome; *median_protein_length*, same as the previous field, but the median; *longest_protein*, the longest protein (amino acids) present in the protein fasta file for this genome; *smallest_protein*, the same as the last field, but the shortest protein; *from_rbh*, blue indicates that the genome was annotated with the 2361 BCnS ALG sequences (**Methods**), and orange indicates that the genome annotation was taken from NCBI; *frac_ologs*, the fraction of the 2361 BCnS ALG orthologs that were present on any scaffold in the genome; *frac_ologs_sig*, the fraction of the 2361 BCnS ALG orthologs that were present in a significant amount on any scaffold in the genome (*p* ≤ 0.05, one-sided Bonferroni-corrected Fisher’s exact test); *frac_ologs_single*, the fraction of the 2361 BCnS ALG orthologs significantly present on a single chromosome (*p* ≤ 0.05, one-sided Bonferroni-corrected Fisher’s exact test); *frac_ologs_{ALG name}*, similar to the previous two fields, this is the fraction of a single BCnS ALG that is significantly present (*p* ≤ 0.05, one-sided Bonferroni-corrected Fisher’s exact test) on one or more scaffolds in the genome.

**Extended Data Fig. 9.**
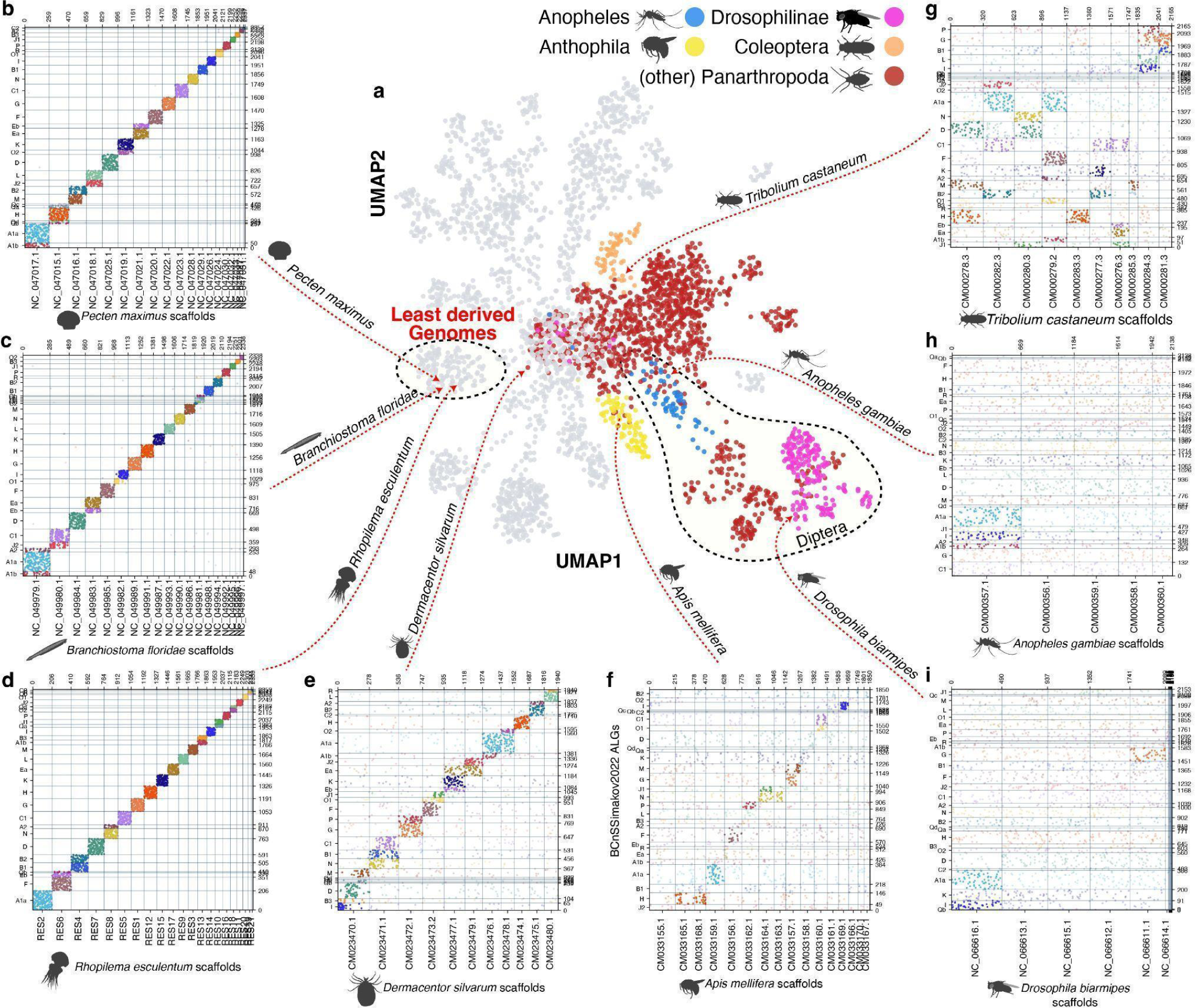
Multi-genome topology reflects genome evolutionary changes. **a**, UMAP from Figure 5 (n=3,631 genomes), with species within Insecta highlighted. The genomes with the fewest chromosomal tectonic changes since the ancestral myriazoan genome are circled. Genomes outside of this circle have varying degrees of tectonic changes from the ancestral BCnS ALGs. A clade with highly derived genomes, the Diptera, is also circled. Most chromosomes within the cluster of the least derived genomes contain single ALGs in **b**, *Pecten maximus*, **c**, *Branchiostoma floridae*, and **d**, *Rhopilema esculentum*. From the centroid of the UMAP mass there are both genomes with significant conservation of BCnS ALGs (*p* < 0.05, one-sided Bonferroni-corrected Fisher’s exact test) **e**, in the tick *Dermacentor silvarum* and derived genomes with poor species sampling. Well-represented clades with lineage-specific genomic changes form clusters and lobes, such as **f**, bees (Anthophila, *Apis mellifera*), and **g**, beetles (Coleoptera, *Tribolium castaneum*). The multi-genome topology also demonstrates how genomes with similar highly reduced karyotypes can be discerned, as seen in **h**, mosquitos(*Anopheles gambiae*), and **i**, drosophilid flies (*Drosophila biarmipes*).

**Extended Data Fig. 10.**
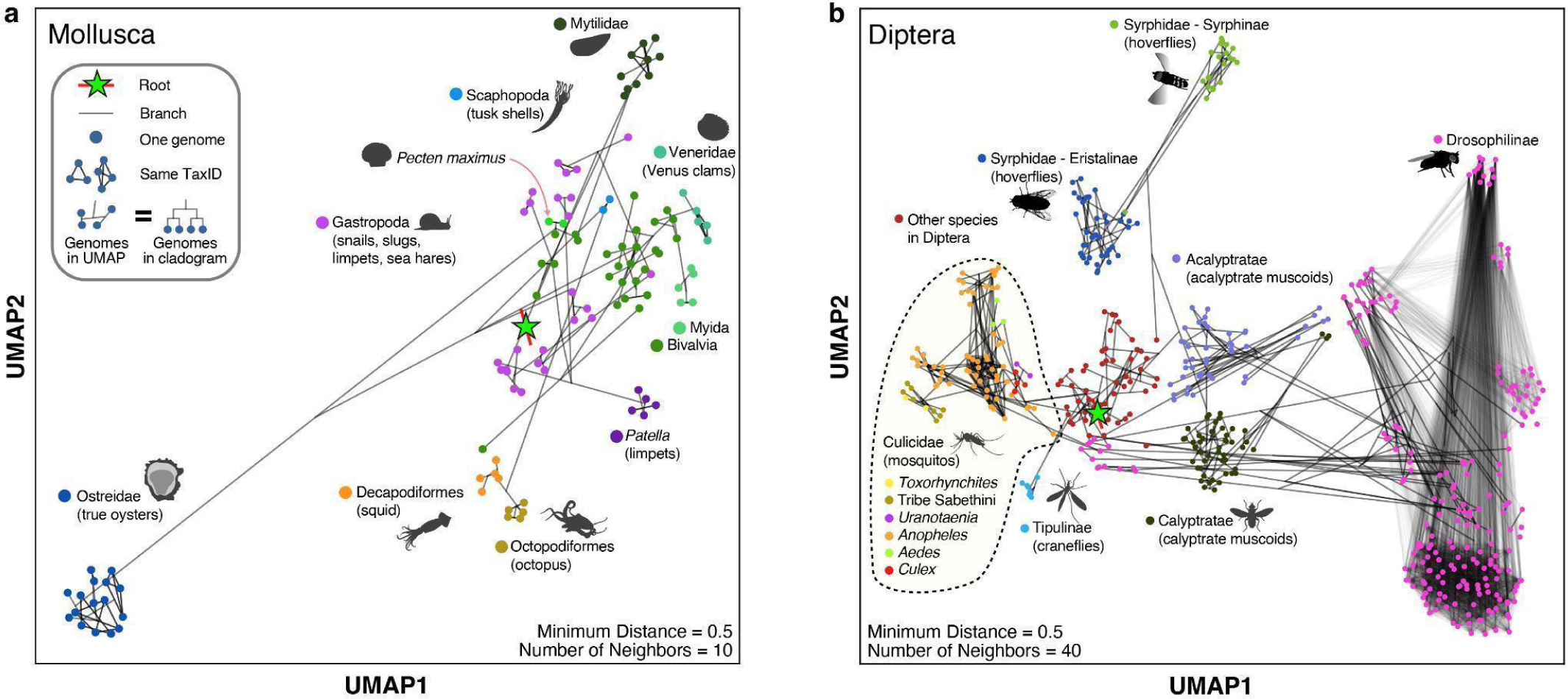
Multi-genome topology reflects genome phylogenetic changes. Both panels are UMAP embeddings of clades within the all-animal dataset. Individual genomes (dots) are colored according to phylogeny, and a cladogram is drawn such that branches between genomes and ancestral nodes are gray lines. **a**, A reimbedding of mollusc genomes (n=108 genomes, 80 species) captures major phylogenetic transitions. Lineage-specific divergences from the ancestral spiralian genome, such as the karyotype reductions in oysters and limpets, can be seen as individual clusters of closely related genomes. Furthermore, lineage-specific changes in squids and octopus since the coleoid cephalopod ancestor are also captured by this approach. **b**, Reimbedding of Diptera genomes (n=509 genomes, 275 species), circled in **Extended Data Fig. 9**. Major dipteran clades colocalize in UMAP space, capturing lineage-specific genomic changes.

**Extended Data Fig. 11.**
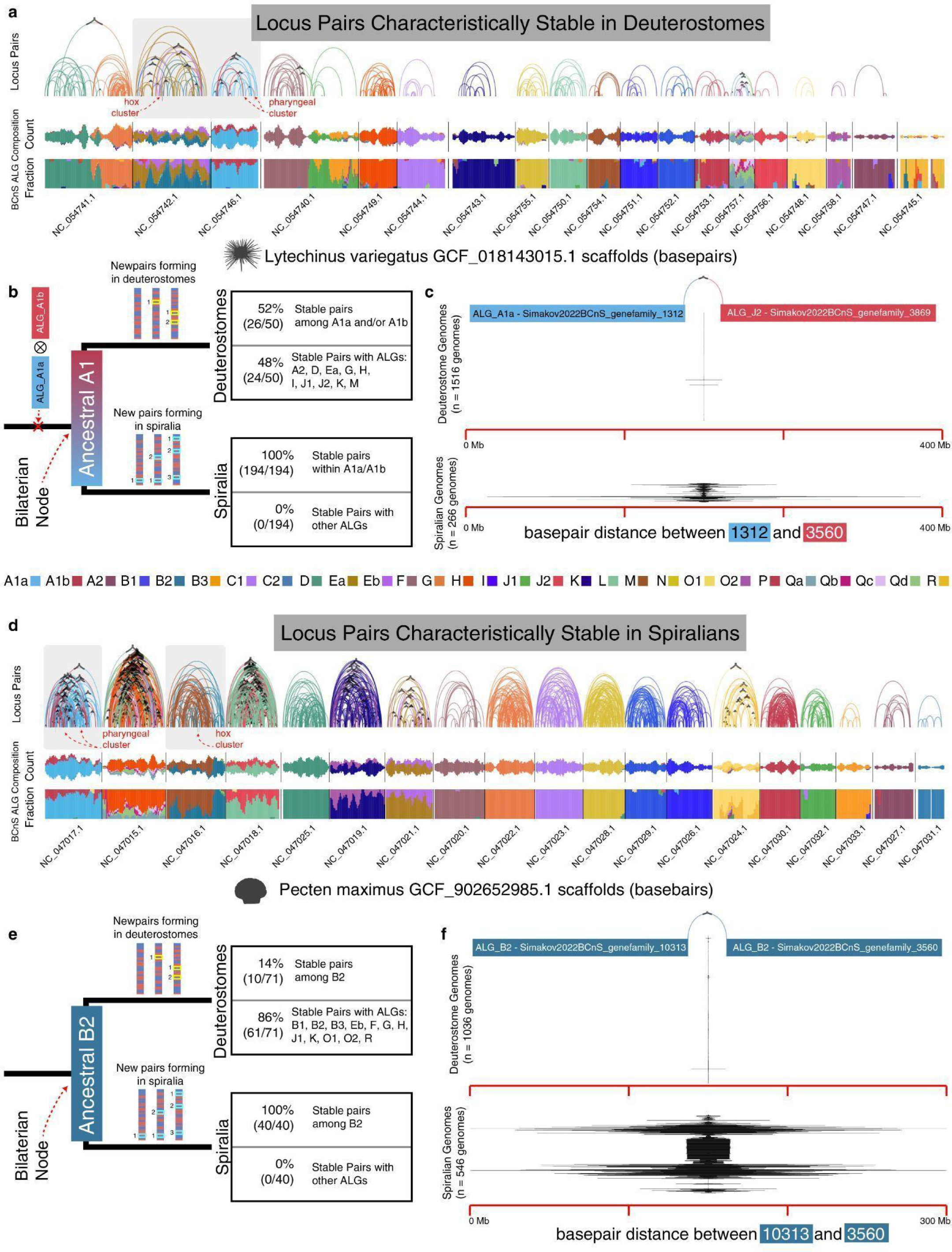
Evolutionary genome topology captures genome-defining features. **a**, The chromosomes of the urchin *Lytechinus variegatus*. The bottom row is the fraction of BCnS ALG gene identities along the chromosome. The middle row is the absolute count of BCnS ALG genes along the chromosomes. The colorful arcs on the top row are the pairs of BCnS ALG that are characteristically stable in the Deuterostomes. Arcs with a black peak on top are stable pairs where the two loci are from different BCnS ALGs. **b**, After ALG A1a and A1b fused onto one chromosome before the ancestor of the Bilateria, the evolutionary history of the ALGs progressed differently in the deuterostomes versus spiralians. **c**, The distance between one locus pair that is characteristically stable in the deuterostomes. The locus is tightly linked in deuterostome genomes, but varies in genomic distance widely, in non-deuterostome genomes. **d**, Locus pairs that are characteristically stable in spiralians, plotted in the context of the scallop *Pecten maximus*. **e**, There are more inter-ALG stable pairs that have formed in deuterostomes with BCnS ALG B2, the ALG associated with the Hox cluster, versus the spiralians. **f**, New characteristically stable pairs can form in lineage-specific ways, even from loci that were present on the same chromosome in the ancestral organism. This is a B2-B2 pair that became characteristically stable in deuterostomes, but is highly variable in spiralians.

## Methods

### Software requirements

The software requirements for these analyses are: bokeh v.3.3.4^83^, ETE3 v.3.1.3^57^, fastapy v.1.0.3^62^, matplotlib v.3.8.0^65^, networkx v.3.1^69^, numpy v.1.26.4^66^, pandas v.2.1.4^60,61^, PIL v.10.2.0^64^, scikit-learn v.1.4.1^67^, scipy v.1.12.0^68^, snakemake v.7.32.3^56^, and umap v.0.5.4^51^.

### Constructing a genome database

We sought to analyze a comprehensive dataset of all chromosome-scale animal genomes available through NCBI. For unannotated genomes we mapped the BCnS proteins to the chromosomes with miniprot v.0.12-r237^59^ with standard parameters.

### Identifying BCnS ALGs

We used hmmer v.3.4^63^ to compare the ancestral BCnS ALGs^1^ to the chromosome-scale genomes of extant animals. The HMM search was performed with hmmsearch from hmmer v.3.4^63^ with a maximum e-value inclusion cutoff of 1×10^5^ (parameter `--incE 1E-5`). For the unannotated genomes, the HMM search was not performed, as the genomes were already annotated with the BCnS ALG orthologs using miniprot. For each genome, the *p*-values of conservation between each pair of BCnS ALGs and extant chromosomes were calculated with a Bonferroni-corrected one-sided Fisher’s exact test^3^.

### Inferring ALG evolutionary history

First, we calculated basic statistics about the presence/absence of BCnS ALGs in extant genomes. One genome for each NCBI taxid was selected to represent each species in the dataset (n=2,291 genomes). To identify whether a BCnS ALG was present on the chromosome of an extant animal, we selected significant BCnS ALG-chromosome combinations (*p* ≤ 0.05, Bonferroni-corrected one-sided Fisher’s exact test). Colocalized ALGs were scored as such simply if they were significant and present on the same scaffold.

Using the same program, we next inferred the history of animal chromosome evolution from the BCnS ancestral linkage groups using a lineage-aware algorithm (n=3631 genomes). Briefly, the program works by reducing the information about BCnS ALG localization in extant genomes into a matrix that encodes whether two BCnS ALGs are colocalized on the same chromosome in that genome. After matrix construction, for each genome the program performs a depth-first search from tip (species) to the animal root and infers the branch along which specific chromosome fusion or loss events occurred. The algorithm pseudo code is:

#### Inputs

- *d*: DataFrame with information about ALG presence and colocalization on single chromosomes.
- *t*: NCBI taxid string for a species in *d*.
- *n*: Node representing the inference point for ALGs. Can be a species or ancestral node.
- *min_M_*: Threshold for determining missing ALGs. We use 0.8. This means that 80% of the genomes being considered must be missing this ALG to consider it lost from that species set. This value below 100% allows for the detection of ALGs that are still present, but difficult to detect, due to the evolutionary history of the chromosomes in that clade.
- *min_N_*: Threshold for determining non-colocalized ALG pairs. We use 0.5. This means that if 50% of the species in this dataset do not have an ALG pair colocalized on the same chromosome, we mark that the ALGs are not colocalized on the branch leading up to the most basal shared node for this clade. If ALGs are colocalized on the same chromosome in the branch leading up to a clade, we would expect to find the colocalization in 100% of species. However, due to dispersion or the limitations of Fisher’s exact test in genomes with few chromosomes, it is possible that using a cutoff of 100% will produce false negatives for ALG colocalizations. Therefore a cutoff below 100% is necessary to find colocalizations. Because the algorithm below polarizes observed colocalizations in one species against a sister clade, it instead measures the percent of species in the sister clade that do not contain the ALG colocalization in question. If convergent fusions were not possible, we would optimally test that 0% of species in the sister clade contain the ALG fusion in question to locate the branch on which the fusion occurred. However, the algorithm must accommodate the possibility of convergent fusions, lineage-specific fusions, the effects of dispersion, and the unequal sampling across the phylogenetic tree. For these reasons, to mark two ALGs as not being colocalized in a given clade, we first check that the ALGs are detectable in that clade in at least 50% of species, and that at least 50% of those species do not have the two ALGs colocalized to the same chromosome.

#### Algorithm 1

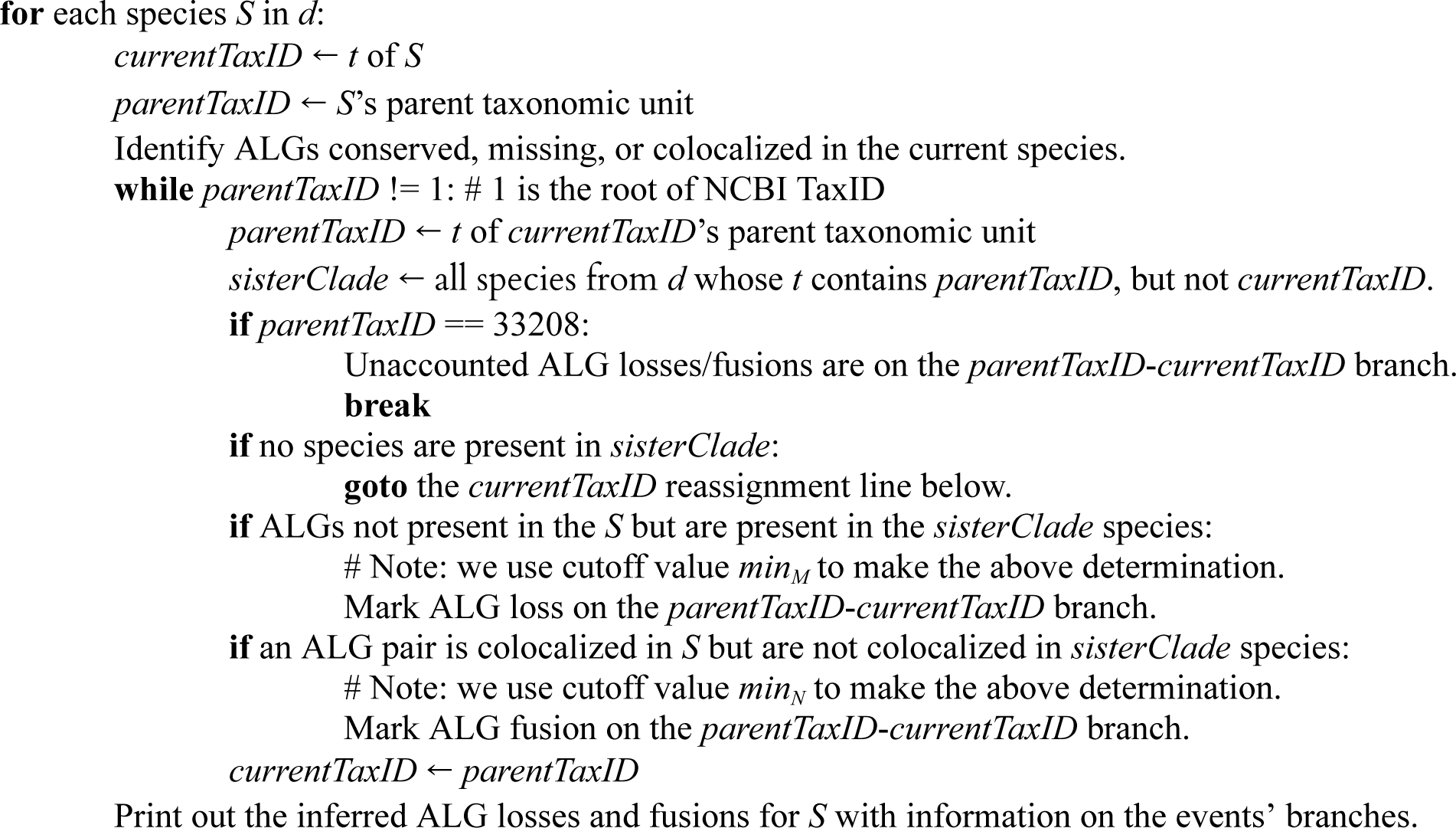

The final result is a string representation of ALG changes for each species.

One calculated value for each change is a measure of uniqueness of this specific change to this evolutionary branch. In our previous work^1,3,27^ we identified major algebraic chromosomal changes that happened on specific evolutionary branches, which became fixed before the crown group of that clade and consequently this evolutionary state can be observed in all extant species in that clade. For example, the fusion of BCnS ALGs A1b and A1a occurred in the ancestor of the Bilateria^1^, and 98.3% of the inferred A1a-A1b fusion events independently inferred from the viewpoint of individual extant species occurred on the branch leading up to Bilateria. Similarly, the known Ea-Eb fusion is captured in this analysis, and 91.1% of all inferred Ea-Eb fusions occurred on this branch. This field can be used as a cutoff to select the most confidently-called ALG change events.

Another value calculated is the measure of the number of species in this clade in which this change could be detected. For example, despite the fact that 98.3% of A1a-A1b events detected were in the branch leading up to Bilateria, this change was only detectable in 54.5% of the species in this clade. This value is reflective of species in which A1a or A1b are no longer detectable due to dispersal.

### Dispersal characterization

We sought to characterize the degree of BCnS ALG dispersal since the myriazoan ancestor, as well as the rate of dispersal over that time period. For each genome, we measured the total fraction of the 2367 BCnS ALG orthologs that were identifiable in the genome, the fraction of 2367 BCnS ALG orthologs that were significantly grouped on any scaffold (*p* ≤ 0.05, Bonferroni-corrected one-sided Fisher’s exact test), the fraction of 2367 BCnS ALG orthologs that were significantly grouped on the largest chromosome (*p* ≤ 0.05, Bonferroni-corrected one-sided Fisher’s exact test), and the fraction each BCnS ALG’s orthologs that are significantly on any chromosome (*p* ≤ 0.05, Bonferroni-corrected one-sided Fisher’s exact test). The genomes were then binned into five categories of decay.

To show the relationship between evolutionary time elapsed and the effects of dispersal, we compared genomes in our dataset (n=22 genomes, 22 species) to the chromosomes of the scallop *Pecten maximus*, which have undergone few inter-chromosomal changes since the myriazoan ancestor. Comparisons between extant species allow for estimation of the amount of dispersal over a given divergence time. We collected estimated divergence times for species in our analyses with TimeTree v.5^70^, and for species with no estimates on TimeTree, we selected the most closely related node in the tree with a time estimate. The value of conservation between *Pecten maximus* and other species was the total fraction of genes on the *Pecten maximus* chromosomes that were significantly conserved on any chromosome in the other species (*p* ≤ 0.05, Bonferroni-corrected one-sided Fisher’s exact test).

### ALG evolution simulations

To estimate whether any given category of ALG fusion, or dispersal in any clade in the tree happened more often than could be expected by random chance in an identical evolutionary scenario, we developed an ALG evolution Monte Carlo simulation.

To estimate a background rate of each type of event on every branch (n=6,376 branches, 6364 taxid nodes), the Monte Carlo simulation was run as follows: for each iteration of the Monte Carlo simulation a random seed value was selected, and the 29 BCnS ALG labels were shuffled. From these labels, the previously described algorithm (**Methods** - *Inferring ALG evolutionary history*) was used to infer the branches on which changes to ALGs occurred. The topology of the cladogram on which the changes were inferred was identical to that used for previous analyses. The results for 10,000 Monte Carlo simulations were summed to estimate the background rate of any given ALG change event happening on any given branch.

### Correlation inference with species diversity

We then measured the one-tailed Spearman’s ranked correlation coefficient between the ALG fusion and loss rates over time versus the species extinction and origination intensity^26^. Briefly, a vector was constructed in which one unit represented one million years. We recursed over the phylogenetic tree and recorded the mean rate of ALG fusions or losses that occurred in each million-year bin for all species in the clade in question.

### Chromosome evolution periodicity analysis

To determine whether changes in chromosomal evolutionary rates have significant underlying periods, we conducted Fourier analyses and Monte Carlo simulations, following the methods described by Rohde and Muller (2005)^26^. Our dataset included rates of ALG fusions and losses across vertebrates and protostomes. We truncated the oldest time for each clade to between 250 to 525 million years ago depending on the age of the clade, and removed the most recent million-year bin, to minimize the chance of over- or under-estimating rates of change. Cubic or quartic polynomials were fitted to the rates, and detrended datasets were created by subtracting these functions. Discrete Fourier transforms were then performed on both raw and zero-padded data to identify the strongest peaks in the spectral power distribution for further analysis.

To estimate false discovery rates for the identified peaks, we employed Monte Carlo simulations of the Fourier analysis through random walks through clustered transitions (type ‘W’), wherein the original dataset was partitioned into vectors and shuffled. We performed simulations with varying block sizes (10 to 50 blocks). Each simulation comprised 5000 random trials, and the results were summarized by reporting the percentage of trials with peak magnitudes matching those observed in the real data. This approach allowed us to assess the periodicity and potential significance of chromosomal changes in evolutionary history.

### Simulation of fusion-with-mixing

We sought to simulate the mixing process of chromosomal fusion-with-mixing (FWM) events on single (simplified) chromosomes. During the simulations, we measured the rate and amount of gene mixing entropy over time on simplified chromosomes.

Such a chromosome consists of genes assigned to two linkage groups (LGs) A and B. Initially, the two groups are each present in contiguous blocks, only joined to each other at one A-to-B transition. This configuration emulates the state of a chromosome after a recent fusion or translocation event, and before any chromosome inversions have spanned into the genes that were originally on two separate chromosomes. Next, the chromosome is subjected to a process of stochastic inversions.

In each chromosome inversion cycle, both inversion breakpoints are chosen with a uniform distribution. We only allow inversions between genes, therefore we can treat the length of the chromosome as a vector of length *L*, one shorter than the number of genes on the chromosome, *L* = *A* + *B* − 1. Therefore, each value between 0 and *L* represents the cutpoint between a single gene. These two random variables range between [0, *L*], and the distance between the two random variables, *d*, has a linear relationship because of the max value imposed by *L*. Given the range of possible values in [0, *L*], the probability density function for *d* can be expressed as 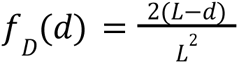. Therefore, the probability of yielding an inversion of length *d* linearly decreases.

At the beginning of the simulation, and after each inversion cycle, we calculate the mixing value *m* — a measurement of the entropy of the A and B genes that were previously on separate chromosomes. For a definition of *m*, see Supplementary Information Section 13.1.2, page 81, in Schultz et al. 2023^3^. Each simulation is run until the value *m* reaches a state in which further inversion events do not cause *m* to change dramatically — in other words after a burn-in period when *m* is exploring the main mass of the probability distribution of *m* in an ideally-mixed genome. After collecting this distribution, it is possible to estimate how many inversions it would take to reach a certain entropy value *m* observed in reality.

An additional feature of the simulation is that the novelty of local gene neighborhoods are measured after each inversion cycle. Every time there is an inversion, there is the possibility for two or more genes to come into close 2D proximity to one another for the first time. The simulation keeps track of newly encountered genes in windows of two to ten genes. This feature allows estimation of the number of inversions required to fully explore the space of possible gene combinations after the chromosome fusion event. The simulation offers the ability to collate the results for many runs and calculate averages or compare the mixing and novel locus encounter behavior under different parameter sets.

We ran chromosome simulations for four different window sizes `W = 1, 2, 5, 10` by executing the simulations with A, B gene counts: `(A, B) = (4, 6), (20, 30), (45, 55), (100, 150), (200, 300), (300, 400), (450, 550)`. Each combination of of chromosome size tuples `(A, B)` and window sizes `W` was simulated 10 times, unless 2 · *W* > *A* + *B*, (effectively, this means that `(A, B) = (4, 6)` was not run with `W = 10`). Mean, standard deviation, median, 1^st^ and 3^rd^ quantiles, as well as minimum and maximum values were calculated from 11 runs with each parameter set.

These simulations can also be run using chromosome sizes, and A and B group sizes, inferred from chromosomal orthology. This mode of operation can infer how far along a chromosome is on the process of mixing through inversions by comparing the observed *m* to the distribution of *m* over the course of the simulation. We used *Pecten maximus* chromosomes that underwent ancient fusion-with-mixing events to demonstrate the varied evolutionary history of these events. First, we identified which *Pecten maximus* fusion with mixing events. Consistent with previous results^1,3^, we found five chromosomes that are composed of two BCnS ancestral linkage groups that have fused and mixed. Among the five chromosomes there are two chromosomes with bilaterian synapomorphies (<u>NC_047017.1</u> - A1a⊗A1b, <u>NC_047021.1</u> - Ea⊗Eb), two chromosomes with spiraliaN syNapomorphies (<u>NC_047018.1</u> - J2⊗L, <u>NC_047019.1</u> - K⊗O2), aNd oNe chromosome with a scallop syNapomorphy (<u>NC_047016.1</u> - B2⊗M).

### Inversion breakpoint inference

We sought to infer rates at which chromosomal inversions become fixed by reconstructing the evolutionary history of homologous chromosomes in groups of closely related species.

Briefly, the analysis uses paf-format whole genome alignment files generated with minimap2 v.2.28^72^, subsetting the alignments to contain homologous chromosomes that have been identified using odp v.0.3.3^3^ or D-GENIES v.1.5.0^73^. Only alignments of a minimal length of 100kb and quality score of 30 are considered. Collinearity breaks are then identified as discontinuities in the sorted pairwise alignment position table. Alignments are considered contiguous if there is a gap smaller than 5 million basepairs between them, and they share the same alignment orientation. This gap parameter can be adjusted for different species combinations to minimize falsely identified breakpoints.

To further study whether collinearity breakpoints overlap with insulation boundaries, breakpointer2 searches for relationships between breakpoints and regions between TADs and other 3D chromosomal topological features. The hypothesis that is tested is whether the breakpoints tend to occur between TADs or other 3D topological features. We define regions that occur between TADs or other 3D topological features as regions with low insulation scores. The null hypothesis is that breakpoints do not tend to occur between TADs and 3D topological features, and we model this distribution of randomly sampled insulation scores. To prepare the data for these analyses, the Hi-C reads were aligned to reference genomes with chromap v.0.2.3^74^ with a quality cutoff of 0. Then pairix v.0.3.7^75,76^ was used to bgzip compress the `.pairs` files from chromap. The program cooler v.0.9.1^77^ was used to generate .cool matrices with bin sizes of 5000 bp, and 100,000 bp. The .cool files were then normalized with HiCNormalization from HiCExplorer v.3.7.2^78^. The normalized `.cool` files were balanced using cooler balance^77^. The normalized and balanced `.cool` files were converted into the .fance format using FAN-C v.0.9.26^79^, and insulation scores were generated with FAN-C for windows of 100 kbp, 250 kbp, 500 kbp, 750 kbp, and 1 Mbp. The resulting insulation scores, in `.bed` format, are used as the input for analyzing breakpoints with insulation score.

To complete the analysis of breakpoints and insulation scores, first the actual breakpoint coordinate is computed as the midpoint between the stop and the start of the previous and the next alignment region, respectively. The insulation scores at such locations and for each chromosome are then compared to the random sample across the same chromosome, with the sample size being the same as the number of observed breakpoints. The median insulation score for the random samples are calculated for 100,000 trials to generate a distribution of insulation score medians for a given chromosome. The medians of the observed breakpoint insulation scores are then tested against the random sampling median distribution using a one-tailed permutation test. The one-tailed permutation test yields a false discovery rate of finding an observed median insulation score smaller than the randomized background distribution. In other words, a false discovery rate of less than 0.05 represents breakpoints that occur in the lowest 5% of possible insulation scores for an identically sized sample, and likely represent regions between TADs or other topological features that have high insulation scores.

We characterized the breakpoints of the octopus species *Octopus vulgaris*(<u>GCA_951406725.2</u>)^71^, *Octopus sinensis* (<u>GCA_006345805.1</u>)^80^, *Octopus bimaculoides* (<u>GCA_001194135.2</u>)^8^, and *Amphioctopus fangsiao* (https://figshare.com/s/fa09f5dadcd966f020f3)^81^. Using the inferred phylogenetic relationship for these four species^81^, and by comparing the `.bed` output of breakpoint2 using bedtools v.2.31.1^82^, we inferred shared and derived inversions along each evolutionary branch for these species. Inversion breakpoints found to exist on two sister species, but not in outgroup species, were assumed to be shared, derived inversions that occurred on the branch leading to the crown group of the two sister species.

### Forms of genome topological analysis

We implemented two forms of topological analysis in this study. In the first type, the topology of entire genomes is encoded for comparison. We term this analysis paradigm as ‘multi-genome topology’, or MGT. This allows not only for interspecies comparisons, but also for comparisons of multiple haplotypes or individuals within a given species. Each sample is a single genome, and when visualized each genome is represented by a single point. Each sample has a vector that contains all of the possible pairwise interactions for an orthology set. The values of the pairwise interactions could be basepair distance, basepair distance divided by chromosome length, Euclidean distance in 3D space, or other weight values derived from these or other sources. In this manuscript, we use the linear distance in basepairs for the distance measure. For measurements of linear distance along a chromosome, ortholog pairs on separate chromosomes should be represented as a number much larger than is possible on real chromosomes. Negative numbers should not be used to encode orthologs on separate chromosomes, as phylogenetically weighting those values could inadvertently bring the values close to a positive value that could be find by bona fide ortholog pairs on the same chromosome. We colloquially refer to visualizations and 2D projections of this matrix as ‘multi-genome topology’ (MGT) plots.

In the second form of topological analysis each sample is a single locus of a single genome, or of a phylogenetically weighted average from a collection of genomes. We refer to this analysis paradigm as ‘multi-locus topology’, or MLT. This analysis enables comparison of changes within genomes, such as colocalizations of loci, chromosomal tectonic changes, new or broken gene linkages. In other words, this analysis paradigm allows for simultaneous analysis of the concepts of colinearity, microsynteny, macrosynteny, and other chromosomal relationship concepts. Each sample is a single locus, and when visualized each locus is represented by a single point. In this manuscript, we use a 2367 x 2367 matrix of the BCnS ALG orthologs as the loci. Each sample, a single BCnS ALG ortholog, has a vector that contains the distance of this BCnS ALG to all other orthologs. This matrix is symmetric. The same rules for representing ortholog pairs on separate chromosomes described for multi-genome topology should also be followed for this approach, multi-locus topology. We refer to 2D projections of this matrix as One-Dot-One-Locus (ODOL) topology plots.

### Evolutionary topology implementation

The multi-locus topology (MLT) matrix creation and visualization procedure differs slightly from that of the multi-genome topology matrices. Each multi-locus topology matrix is a symmetric *N* × *N* matrix, where *N* is the total number of orthologous loci across the entire dataset. The goal of this matrix is to express the topology of a genome for a single genome or collection of species, and to be able to compare topologies between species or clades. For this reason, it is necessary to average the distances between ortholog pairs for all of the samples in the comparison.

To perform phylogenetic weighting of the matrix, we first implement a directed, acyclic graph (DAG) reflecting the structure of the evolutionary relationships between species. In the phylogenetic weighting algorithm, the *N* × *N* distance matrix of each node is calculated by normalizing the contribution of different species depending on their relatedness. This prevents clades with many representative genomes, such as the many sequenced genomes of drosophilids or vertebrates, from biasing the averaged weights. In this approach, ortholog pairs that exist on separate chromosomes and are encoded as large values (999,999,999,999) are not treated differently in the weighting algorithm.

A more detailed explanation of the algorithm is below:

1. We construct the DAG, and its inverse graph, from NCBI taxid lineage information, and the branch lengths between taxids in the lineages were encoded as 1s. In this graph, nodes are extant species or ancestors of one or more species. Edges represent branches on the phylogenetic tree, and the value of the edge is the length of the branch. The root is determined and is defined as the node that has no parents. Below, we will refer to a node of the DAG as *DAG*[*node*1], and the edge weight between two nodes as *DAG*[*node*1][*node*2].
2. After constructing the DAG, the branch lengths are normalized such that the total distance from the root to any leaf is 1. The reason this is done is to enable matrix multiplication to be used to perform the phylogenetic averaging. Therefore, both cladograms and phylograms can be used as input. The algorithm finds the longest path from the root and scales the branch lengths accordingly. Let *P_longest_* be the longest path from the root to a leaf with ℓ nodes (hence ℓ − 1 edges). We define

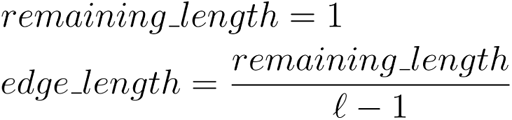

For each child *c* of node *n*:

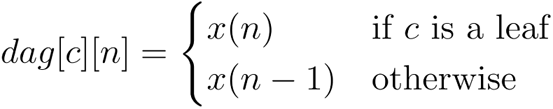

The algorithm then recursively adjusts the branch lengths for the subtree of each child node. This is all performed in the function `adjacency_dag.normalize_branch_lengths()`.

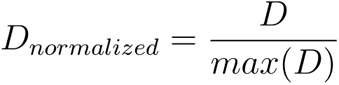
3. A distance matrix is generated for each pair of tips (*t_i_, t_j_*), where each tip *t_i_* is one of the species in the tree that is being averaged. The distance between two tips (*t_i_, t_j_*) is the sum of the weights along the path from *t_i_* to the last common ancestor node shared with *t_j_*. The distance matrix, *D*, is then normalized to be between the maximum weight recorded in the matrix.
4. The phylogenetic weight calculation is as follows: The total distance for each sample: 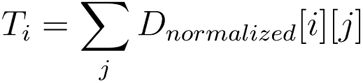 The weight vector for the samples: 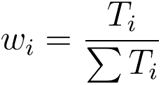
5. The phylogenetic weight matrix is calculated by calculating the dot product of the transposed multi-genome topology (MGT) *matrix^T^* and the weight vector *w*.

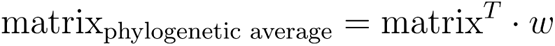
6. The final vector is transformed into the *N × N* distance matrix based on the predetermined mapping of MGT matrix indices to ortholog IDs.

### Graph database integration

We used the Neo4j Community Edition server v.5.20.0 (https://neo4j.com) to build a graph database of orthology at multiple levels across the species utilized in this study, which consists of relationships, nodes, and node properties, all of which can be used for various types of semantic queries. The BCnS ALGs and orthogroups (separate ALG and OG nodes), the orthologous proteins from each species (gene nodes), and the scaffolds for each species (chromosome nodes) were all loaded as different node types into the graph. The *orthologous* relationship in our graph database connects the chromosome nodes with the ALG nodes, and the gene nodes with the OG nodes. The *location* relationship connects the gene nodes with the chromosome nodes, and the OG nodes with the ALG nodes. Both gene and chromosome nodes have the following properties: *TaxID*, which is the NCBI taxid of the species; *Assembly*, which is the version of the genome assembly; *Species*, which is the scientific name of the species; *Lineage*, which is the complete set of NCBI taxids forming the taxonomic lineage of the organism starting from the species ID; and *name*, which is the name of the gene or the chromosome. In cases when a chromosome node is actually not a real chromosome but a smaller scaffold that may still be present in a chromosome level genome assembly, a node is still labeled as a chromosome. However, we note that we only downloaded the chromosomal scaffolds for the genomes we collected from NCBI. In addition, the gene nodes have the properties *Chromosome* and *CoordinateStart*, which define the chromosome the gene is located on and the genomic coordinate where the gene starts. Both gene and chromosome nodes are labeled with a string with the first field as the *CHR* or *GENE*, plus the NCBI taxid of the source organism. For instance, nodes labeled as CHR9606 and GENE9606 belong to *Homo sapiens*. This taxid identifier can be used to query nodes with the same taxonomic source. A schematic of the graph database and an example illustration comparing the orthology of genes from an ALG between three different pairs of species are shown in **Extended Data Fig. 6a,b**.

The graph was visualized with Neo4j Browser v.5.15.0. We further designed a tool using the Neo4j Cypher language to generate queries to retrieve subgraphs that satisfy different topological questions. The query forms provide the user with options such as species and ALG names, to generate Cypher codes that can be copied or downloaded and run on our browser to explore the subgraph of interest. The graph data structure is also available as a Neo4j image for loading it locally. We have also included the graph as a GraphML file which is the common format for exchanging graph structure data that can be used in other graph database softwares, as a JSON file, and as a CSV file, all of which may also be loaded into other graph data structures for exploration.

## Acknowledgements

D.T.S., D.D., and O.S. were supported by the European Research Council’s Horizon 2020: European Union Research and Innovation Programme, grant No. 945026. D.D. was also supported by the Vienna Haus des Meeres Verein: Rupert Riedl Prize. F.S. was supported by the Austrian Science Fund grant P32190 to O.S. The authors thank Nils Carqueville and Jacob Winnikoff for scientific discussions. The computation for this work was performed using the Life Science Compute Cluster (LiSC) of the University of Vienna.

## Author Contributions

O.S. and D.T.S. designed the scientific objectives of the study. Various parts of the code were written by D.T.S., A.B., F.S., and D.D. D.T.S., A.B., and F.S. created figures and tables. D.T.S. and O.S. wrote the manuscript under input from all other authors. All of the authors contributed to the interpretation, presentation and writing of the Article and the Supplementary Information.

## Supplementary information

The online version contains supplementary information.

## Materials & Correspondence

Correspondence to: D.T. Schultz, O. Simakov

## Data Availability

The genomes used are available through the INSDC database (ENA, NCBI, DDBJ) accession numbers: *Abramis brama* (GCA_022829085.1), *Abrostola tripartita* (GCA_905340225.1), *Abrostola triplasia* (GCA_946251915.1), *Acanthisitta chloris* (GCA_016904835.1), *Acanthochromis polyacanthus* (GCA_021347895.1, GCF_021347895.1), *Acanthococcus lagerstroemiae* (GCA_031841125.1), *Acanthopagrus latus* (GCA_013436515.1, GCA_020080195.1, GCA_904848185.1, GCF_904848185.1), *Acanthosoma haemorrhoidale* (GCA_930367205.1), *Accipiter gentilis* (GCA_929443795.1, GCA_929443795.2, GCF_929443795.1), *Acentria ephemerella* (GCA_943193645.1), *Achatina immaculata* (GCA_009760885.1), *Achlya flavicornis* (GCA_947623365.1), *Acholoe squamosa* (GCA_949317995.1), *Acinonyx jubatus* (GCF_027475565.1, GCA_027475565.2), *Acipenser oxyrinchus oxyrinchus* (GCA_030684275.1), *Acipenser ruthenus* (GCA_902713425.2, GCF_902713425.1, GCA_010645085.2, GCF_010645085.2, GCA_902713435.2), *Acleris cristana* (GCA_948252455.1), *Acleris emargana* (GCA_927399475.2), *Acleris holmiana* (GCA_949316455.1), *Acleris literana* (GCA_946894065.1), *Acleris sparsana* (GCA_923062465.1), *Acomys cahirinus* (GCA_029890205.1), *Acomys dimidiatus* (GCA_907164435.1), *Acomys kempi* (GCA_907164505.1), *Acomys percivali* (GCA_907169655.1), *Acomys russatus* (GCF_903995435.1, GCA_903995435.1), *Acridotheres tristis* (GCA_027559615.1), *Acrobasis consociella* (GCA_963555685.1), *Acrobasis repandana* (GCA_963576875.1), *Acrobasis suavella* (GCA_943193695.1), *Acrocephalus scirpaceus scirpaceus* (GCA_910950805.1), *Acrocera orbiculus* (GCA_947359355.1), *Acronicta aceris* (GCA_910591435.1), *Acronicta leporina* (GCA_947256265.1), *Acronicta psi* (GCA_946251955.1), *Acropora hyacinthus* (GCA_020536085.1), *Acropora millepora* (GCF_013753865.1, GCA_013753865.1), *Acrossocheilus wenchowensis* (GCA_029620275.1), *Acutogordius australiensis* (GCA_030555515.1), *Acyrthosiphon pisum* (GCF_005508785.2, GCA_005508785.2), *Adalia bipunctata* (GCA_910592335.1), *Adalia decempunctata* (GCA_951802165.1), *Adelphocoris suturalis* (GCA_030762985.1), *Adineta vaga* (GCA_021613535.1), *Aedes aegypti* (GCA_002204515.1, GCF_002204515.2), *Aedes aegypti aegypti* (GCA_025407655.1), *Aelia acuminata* (GCA_911387785.2), *Agapetus fuscipes* (GCA_951799405.1), *Agapostemon virescens* (GCA_028453745.1), *Agelaius phoeniceus* (GCA_020745825.3, GCF_020745825.1), *Agelas oroides* (GCA_949130485.1), *Agelastica alni* (GCA_950111635.1), *Agonopterix arenella* (GCA_927399405.1), *Agonopterix heracliana* (GCA_963693445.1), *Agonopterix subpropinquella* (GCA_922987775.1), *Agonum fuliginosum* (GCA_947534325.1), *Agrilus cyanescens* (GCA_947389935.1), *Agrilus mali* (GCA_029378335.1), *Agriopis aurantiaria* (GCA_914767915.1), *Agriopis leucophaearia* (GCA_949125355.1), *Agriopis marginaria* (GCA_932305915.1), *Agriotes lineatus* (GCA_940337035.1), *Agriphila geniculea* (GCA_943789525.1), *Agriphila straminella* (GCA_950108535.1), *Agriphila tristella* (GCA_928269145.1), *Agrochola circellaris* (GCA_914767755.1), *Agrochola litura* (GCA_949152395.1), *Agrochola lota* (GCA_947364205.1), *Agrochola lychnidis* (GCA_963680765.1), *Agrochola macilenta* (GCA_916701695.1), *Agrotis clavis* (GCA_954870645.1), *Agrotis exclamationis* (GCA_950005045.1), *Agrotis ipsilon* (GCA_028554685.1), *Agrotis puta* (GCA_943136025.2), *Agrypnus murinus* (GCA_929113105.1), *Ahaetulla prasina* (GCA_028640845.1, GCF_028640845.1), *Ailuropoda melanoleuca* (GCA_002007445.3, GCA_029963865.1, GCF_002007445.2), *Albula goreensis* (GCA_022829145.1), *Alca torda* (GCA_008658365.1), *Alcis repandata* (GCA_949125135.1), *Aldabrachelys gigantea* (GCA_026122505.1), *Alectoris magna* (GCA_030625065.1), *Aleiodes testaceus* (GCA_963565655.1), *Alentia gelatinosa* (GCA_950022655.1), *Alitta virens* (GCA_932294295.1), *Allacma fusca* (GCA_947179485.1), *Alligator mississippiensis* (GCA_030867095.1, GCF_030867095.1), *Allophyes oxyacanthae* (GCA_932294325.1), *Alloplasta piceator* (GCA_946863875.1), *Allygus modestus* (GCA_963675035.1), *Alosa alosa* (GCF_017589495.1, GCA_017589495.2), *Alosa sapidissima* (GCF_018492685.1, GCA_018492685.1, GCA_019202745.1), *Alsophila aescularia* (GCA_946251855.1), *Alviniconcha strummeri* (GCA_963584105.1), *Amaurobius ferox* (GCA_951213105.1), *Amazona aestiva* (GCA_017639355.1), *Amblyjoppa fuscipennis* (GCA_963575735.1), *Amblyraja radiata* (GCA_010909765.2, GCF_010909765.2), *Amblyteles armatorius* (GCA_933228735.2), *Ambystoma mexicanum* (GCA_002915635.3), *Ameiurus melas* (GCA_012411365.1), *Amia calva* (GCA_017591415.1), *Ammodytes dubius* (GCA_026122265.1), *Ammodytes marinus* (GCA_949987685.1), *Ammospiza caudacuta* (GCA_027887145.1, GCF_027887145.1), *Ammospiza nelsoni* (GCF_027579445.1, GCA_027579445.1), *Amorpha juglandis* (GCA_949126905.1), *Amphiduros pacificus* (GCA_949316495.1), *Amphilophus citrinellus* (GCA_013435755.1), *Amphimallon solstitiale* (GCA_963170755.1), *Amphipoea lucens* (GCA_947508005.1), *Amphipoea oculea* (GCA_945859645.1), *Amphiprion clarkii* (GCA_027123335.1), *Amphiprion ocellaris* (GCA_022539595.1, GCF_022539595.1), *Amphiprion percula* (GCA_003047355.2), *Amphipyra berbera* (GCA_910594945.2), *Amphipyra tragopoginis* (GCA_905220435.1), *Anabas testudineus* (GCA_900324465.2, GCF_900324465.2, GCA_900324465.3), *Anableps anableps* (GCA_014839685.1), *Anadara kagoshimensis* (GCA_021292105.1), *Anania crocealis* (GCA_949315895.1), *Anania fuscalis* (GCA_950371115.1), *Anania hortulata* (GCA_963576865.1), *Anarsia innoxiella* (GCA_947563765.1), *Anas platyrhynchos* (GCF_003850225.1, GCF_015476345.1, GCA_015476345.1, GCA_008746955.3, GCA_003850225.1, GCA_900411745.1, GCA_030704485.1), *Anas platyrhynchos platyrhynchos* (GCA_002743455.1), *Anaspis maculata* (GCA_949128115.1), *Anastrepha ludens* (GCF_028408465.1, GCA_028408465.1), *Anastrepha obliqua* (GCA_027943255.1, GCF_027943255.1), *Ancistrocerus nigricornis* (GCA_916049575.1), *Andrena bicolor* (GCA_960531205.1), *Andrena bucephala* (GCA_947577245.1), *Andrena camellia* (GCA_029168295.1, GCA_029448645.1), *Andrena chrysosceles* (GCA_963855975.1), *Andrena dorsata* (GCA_929108735.1), *Andrena fulva* (GCA_946251845.1), *Andrena haemorrhoa* (GCA_910592295.1), *Andrena hattorfiana* (GCA_944738655.2), *Andrena minutula* (GCA_929113495.1), *Andrena trimmerana* (GCA_951215215.1, GCA_952773225.1), *Anguilla anguilla* (GCA_018320845.1, GCF_013347855.1, GCA_013347855.1), *Anguilla japonica* (GCA_025169545.1), *Anguilla rostrata* (GCA_018555375.2), *Anisus vortex* (GCA_949126835.1), *Anolis carolinensis* (GCF_000090745.2, GCA_000090745.2), *Anomalospiza imberbis* (GCA_031753505.1), *Anomoia purmunda* (GCA_951828415.1), *Anopheles albimanus* (GCF_013758885.1, GCA_000349125.2, GCA_015501965.1, GCA_013758885.1), *Anopheles aquasalis* (GCA_943734665.1, GCF_943734665.1), *Anopheles arabiensis* (GCF_016920715.1, GCA_016920715.1), *Anopheles atroparvus* (GCA_015501955.1), *Anopheles bellator* (GCF_943735745.2, GCA_943735745.2), *Anopheles coluzzii* (GCA_016097195.1, GCA_016508615.1, GCA_943734685.1, GCA_016920705.1, GCA_016097205.1, GCA_016097185.1, GCA_016097225.1, GCF_016920705.1, GCA_016097175.1, GCA_016097095.1, GCF_943734685.1), *Anopheles coustani* (GCA_943734705.2, GCF_943734705.1), *Anopheles cruzii* (GCA_943734635.1, GCF_943734635.1), *Anopheles darlingi* (GCF_943734745.1, GCA_943734745.1), *Anopheles funestus* (GCA_943734845.1, GCF_943734845.2, GCA_003951495.1), *Anopheles funestus-like sensu Spillings et al. (2009)* (GCA_016170035.1), *Anopheles gambiae* (GCA_943734735.2), *Anopheles gambiae str. PEST* (GCF_000005575.2, GCA_000005575.1), *Anopheles longipalpis* (GCA_016170015.1), *Anopheles maculipalpis* (GCA_943734695.1, GCF_943734695.1), *Anopheles marshallii* (GCA_943734725.1, GCF_943734725.1), *Anopheles merus* (GCF_017562075.2, GCA_017562075.2), *Anopheles moucheti* (GCA_943734755.1, GCF_943734755.1), *Anopheles nili* (GCA_943737925.1, GCF_943737925.1), *Anopheles parensis* (GCA_016254315.1), *Anopheles rivulorum* (GCA_018908115.1), *Anopheles sp. NFL-2015* (GCA_016171315.1), *Anopheles stephensi* (GCF_013141755.1, GCA_023078585.1, GCA_003448955.2, GCA_013141755.1, GCA_017562265.1, GCA_020882675.1, GCA_003448975.2), *Anopheles vaneedeni* (GCA_016170025.1), *Anopheles ziemanni* (GCF_943734765.1, GCA_943734765.2), *Anoplius nigerrimus* (GCA_914767735.1), *Anoplopoma fimbria* (GCA_027596085.2, GCF_027596085.1), *Anorthoa munda* (GCA_945859665.1), *Anser cygnoides* (GCA_026259575.1, GCA_025388735.1), *Anser cygnoides domesticus* (GCA_030515125.1), *Anser indicus* (GCA_025583725.1), *Antechinus flavipes* (GCF_016432865.1, GCA_016432865.2), *Antedon bifida* (GCA_963402885.1), *Antennarius maculatus* (GCA_013358685.1), *Anthidium xuezhongi* (GCA_022405125.1), *Anthocharis cardamines* (GCA_905404175.1), *Anthonomus grandis grandis* (GCA_022605725.3, GCF_022605725.1), *Anthonomus grandis thurberiae* (GCA_030068095.1), *Anthophora plumipes* (GCA_951804975.1), *Anticlea derivata* (GCA_947579855.1), *Antitype chi* (GCA_947359405.1), *Antrozous pallidus* (GCA_027563665.1), *Apalone spinifera* (GCA_030068395.1), *Apamea anceps* (GCA_951799955.1), *Apamea crenata* (GCA_949629185.1), *Apamea epomidion* (GCA_947507525.1), *Apamea monoglypha* (GCA_911387795.2), *Apamea sordens* (GCA_945859715.1), *Apeira syringaria* (GCA_934044485.1), *Apeltes quadracus* (GCA_021346845.1), *Aphelinus atriplicis* (GCA_020882685.1), *Aphelinus certus* (GCA_020882725.1), *Aphidius gifuensis* (GCA_014905175.1, GCF_014905175.1), *Aphidoletes aphidimyza* (GCA_030463065.1), *Aphis gossypii* (GCA_917880025.4, GCA_020184165.1, GCA_020184175.2, GCF_020184175.1), *Aphrocallistes beatrix* (GCA_963281255.1), *Aphrophora alni* (GCA_963513935.1), *Apis cerana* (GCA_020141525.2, GCA_029531695.1, GCA_029169275.1), *Apis cerana cerana* (GCA_011100585.1), *Apis mellifera* (GCF_000002195.4, GCF_003254395.2, GCA_000002195.1, GCA_003254395.2), *Apis mellifera carnica* (GCA_013841245.2), *Apis mellifera ligustica* (GCA_019321825.1), *Apis mellifera mellifera* (GCA_003314205.2), *Aplidium turbinatum* (GCA_918807975.1), *Aplocera efformata* (GCA_921293045.1), *Aplochiton taeniatus* (GCA_017639675.1), *Aplysina aerophoba* (GCA_949841015.1), *Apocheima hispidaria* (GCA_947579745.1), *Apoda limacodes* (GCA_946406115.1), *Apodemus sylvaticus* (GCF_947179515.1, GCA_947179515.1), *Apoderus coryli* (GCA_911728435.2), *Apolygus lucorum* (GCA_009739505.2), *Apomyelois bistriatella* (GCA_947044815.1), *Aporia crataegi* (GCA_912999735.1), *Aporophyla lueneburgensis* (GCA_932294355.1), *Aporophyla nigra* (GCA_947507805.1), *Aporrectodea icterica* (GCA_963556255.1), *Apotomis betuletana* (GCA_932273695.1), *Apotomis capreana* (GCA_947623375.1), *Apotomis turbidana* (GCA_905147355.2), *Aproaerema taeniolella* (GCA_949987775.1), *Apus apus* (GCA_020740795.1, GCF_020740795.1), *Aquila chrysaetos chrysaetos* (GCF_900496995.4, GCA_900496995.4), *Ara ararauna* (GCA_028858755.1, GCA_028858555.1), *Aradus depressus* (GCA_963662175.1), *Archilochus colubris* (GCA_023079485.1), *Archips crataeganus* (GCA_947859365.1), *Archips xylosteana* (GCA_947563465.1), *Archivesica marissinica* (GCA_014843695.1), *Archocentrus centrarchus* (GCA_007364275.2, GCF_007364275.1), *Arctocephalus townsendi* (GCA_028646355.1), *Argentina silus* (GCA_951799395.1), *Argiope bruennichi* (GCF_947563725.1, GCA_947563725.1, GCA_015342795.1), *Argynnis bischoffii washingtonia* (GCA_033884665.1), *Argyresthia albistria* (GCA_963457715.1), *Argyresthia goedartella* (GCA_949825045.1), *Argyrosomus regius* (GCA_946959425.1), *Aricia agestis* (GCA_944452695.1, GCA_905147365.1, GCF_905147365.1), *Aricia artaxerxes* (GCA_937612035.1), *Arion vulgaris* (GCA_020796225.1), *Artemia franciscana* (GCA_032884065.1), *Artemia sinica* (GCA_027921565.1), *Arvicanthis niloticus* (GCF_011762505.1, GCA_011762505.1), *Arvicola amphibius* (GCA_903992535.2, GCF_903992535.2), *Ascaris suum* (GCA_013433145.1), *Ascidia mentula* (GCA_947561715.1), *Aselliscus stoliczkanus* (GCA_033961575.1), *Aspidapion aeneum* (GCA_963576565.1), *Aspiorhynchus laticeps* (GCA_023376895.1), *Astatotilapia calliptera* (GCA_900246225.3, GCF_900246225.1, GCA_900246225.5), *Asterias amurensis* (GCA_032118995.1), *Asterias rubens* (GCF_902459465.1, GCA_902459465.3), *Asteroscopus sphinx* (GCA_949699075.1), *Astyanax mexicanus* (GCA_023375845.1, GCA_023375835.1, GCA_000372685.2, GCF_023375975.1, GCA_019721115.1, GCA_023375975.1, GCF_000372685.2), *Atethmia centrago* (GCA_905333075.3), *Athalia rosae* (GCF_917208135.1, GCA_917208135.1), *Atherix ibis* (GCA_958298945.1), *Athetis lepigone* (GCA_033675125.1), *Athrips mouffetella* (GCA_947532105.1), *Athripsodes cinereus* (GCA_947579605.1), *Auanema freiburgensis* (GCA_030370435.1), *Augochlora pura* (GCA_028453695.1), *Augochlorella aurata* (GCA_028455555.1), *Autographa gamma* (GCA_905146925.1), *Autographa pulchrina* (GCA_905475315.1), *Aythya baeri* (GCA_026413565.1), *Aythya fuligula* (GCA_009819795.1, GCF_009819795.1), *Aythya marila* (GCA_029042245.1), *Azorinus chamasolen* (GCA_963576725.1), *Babyrousa celebensis* (GCA_028533215.1), *Baccha elongata* (GCA_951217065.1), *Bacillus rossius redtenbacheri* (GCA_032445375.1), *Bactericera cockerelli* (GCA_024516035.1), *Bactrocera correcta* (GCA_027475135.1), *Bactrocera dorsalis* (GCF_023373825.1, GCA_023373825.1, GCA_030710565.1), *Bactrocera neohumeralis* (GCA_024586455.2, GCF_024586455.1), *Bactrocera tryoni* (GCA_016617805.2, GCF_016617805.1), *Balaenoptera acutorostrata* (GCA_949987535.1, GCF_949987535.1), *Balaenoptera musculus* (GCF_009873245.2, GCA_009873245.3), *Balaenoptera ricei* (GCF_028023285.1, GCA_028023285.1), *Balanococcus diminutus* (GCA_959613365.1), *Balearica regulorum gibbericeps* (GCA_011004875.1), *Barbatula barbatula* (GCA_947034865.1), *Barbus barbus* (GCA_023566175.1, GCA_936440315.1), *Bathymodiolus brooksi* (GCA_963680875.1), *Bathymodiolus septemdierum* (GCA_963383655.1), *Beaufortia kweichowensis* (GCA_019155185.1), *Bedellia somnulentella* (GCA_963576735.1), *Bellardia bayeri* (GCA_950370525.1), *Bellardia pandia* (GCA_916048285.2), *Belonocnema kinseyi* (GCA_010883055.1, GCF_010883055.1), *Bembecia ichneumoniformis* (GCA_910589475.2), *Bemisia tabaci* (GCA_918797505.1), *Beris chalybata* (GCA_949128065.1), *Beris morrisii* (GCA_951812415.1), *Betta splendens* (GCA_024678985.1, GCA_013403625.1, GCF_900634795.4, GCA_900634795.4), *Bibio marci* (GCA_910594885.2), *Bicyclus anynana* (GCA_947172395.1, GCF_947172395.1), *Bimastos eiseni* (GCA_959347315.1), *Biomphalaria glabrata* (GCF_947242115.1, GCA_947242115.1), *Biston betularia* (GCA_905404145.2), *Biston stratarius* (GCA_950106695.1), *Blastobasis adustella* (GCA_907269095.1), *Blastobasis lacticolella* (GCA_905147135.1), *Blastomussa wellsi* (GCA_947652115.1), *Blennius ocellaris* (GCA_963422515.1), *Blera fallax* (GCA_946965025.1), *Blomia tropicalis* (GCA_029204025.1), *Bolinopsis microptera* (GCA_026151205.1), *Boloria euphrosyne* (GCA_951802675.2), *Boloria selene* (GCA_905231865.2), *Bombina bombina* (GCA_027579735.1, GCF_027579735.1), *Bombus affinis* (GCA_024516045.2, GCF_024516045.1), *Bombus breviceps* (GCA_014825925.1), *Bombus campestris* (GCA_905333015.3), *Bombus haemorrhoidalis* (GCA_014825975.1), *Bombus hortorum* (GCA_905332935.1), *Bombus huntii* (GCF_024542735.1, GCA_024542735.1), *Bombus hypnorum* (GCA_911387925.2), *Bombus ignitus* (GCA_014825875.1), *Bombus pascuorum* (GCF_905332965.1, GCA_905332965.1), *Bombus pratorum* (GCA_930367275.1), *Bombus pyrosoma* (GCA_014825855.1, GCF_014825855.1), *Bombus sylvestris* (GCA_911622165.2), *Bombus terrestris* (GCA_910591885.2, GCA_000214255.1, GCF_910591885.1, GCF_000214255.1), *Bombus turneri* (GCA_014825825.1), *Bombus vestalis* (GCA_963556215.1), *Bombylius discolor* (GCA_939192795.1), *Bombylius major* (GCA_932526495.1), *Bombyx mandarina* (GCA_030267445.2), *Bombyx mori* (GCA_014905235.2, GCF_014905235.1, GCA_027497135.1, GCA_027497115.1, GCA_026075555.1, GCA_030269925.2), *Borostomias antarcticus* (GCA_949987555.1), *Bos gaurus* (GCA_014182915.2), *Bos gaurus x Bos taurus* (GCA_946052875.1), *Bos grunniens* (GCA_027580245.1, GCA_005887515.3), *Bos grunniens x Bos taurus* (GCA_009493645.1, GCA_009493655.1), *Bos indicus* (GCA_030271805.1, GCA_026262375.1, GCA_002933975.1, GCA_000247795.2, GCA_030270995.1, GCA_030271795.1, GCA_030267355.1, GCA_030272505.1, GCA_030270855.1, GCF_000247795.1, GCA_030270935.1, GCA_030270945.1, GCA_026262465.1, GCA_030272475.1, GCA_030272235.1, GCA_030270865.1, GCA_030272135.1, GCA_026262515.1, GCA_030270805.1, GCA_029378735.1, GCA_030270725.1, GCA_030270715.1, GCA_030270705.1, GCA_030270685.1, GCA_030270955.1, GCA_030269815.1, GCA_030269505.1, GCA_030267375.1, GCA_026262545.1, GCA_030272495.1, GCA_030270825.1, GCA_029378745.1, GCA_030267345.1, GCA_030270845.1), *Bos indicus x Bos taurus* (GCA_003369685.2, GCA_003369695.2, GCF_003369695.1), *Bos javanicus* (GCF_032452875.1, GCA_032452875.1), *Bos mutus* (GCA_027580195.1, GCA_002968435.1), *Bos taurus* (GCA_028973685.2, GCA_034097375.1, GCF_002263795.3, GCF_000003205.7, GCA_905123515.1, GCA_030378505.1, GCA_002263795.4, GCA_021234555.1, GCA_000003055.5, GCA_000003205.6, GCA_905123885.1, GCF_000003055.6, GCA_947034695.1, GCA_021347905.1), *Bos taurus x Bison bison* (GCA_018282465.1, GCA_018282365.1, GCA_030254855.1), *Bostrychus sinensis* (GCA_020698575.1), *Brachylomia viminalis* (GCA_937001585.2), *Brachymystax lenok tsinlingensis* (GCA_031305455.1, GCA_030435695.1), *Brachyopa scutellaris* (GCA_949775065.1), *Brachypalpus laphriformis* (GCA_945910035.1), *Brachyptera putata* (GCA_907164805.1), *Brachypterus glaber* (GCA_958510825.1), *Bradysia coprophila* (GCA_014529535.1, GCA_014529535.2, GCF_014529535.1), *Branchellion lobata* (GCA_947562095.1), *Branchiostoma floridae* (GCA_000003815.2, GCF_000003815.2), *Branchiostoma floridae x Branchiostoma belcheri* (GCA_022788385.1, GCA_019207075.1, GCA_019207045.1), *Branchiostoma floridae x Branchiostoma japonicum* (GCA_013266295.2, GCA_015852565.1), *Branchiostoma lanceolatum* (GCA_927797965.1), *Branchipolynoe longqiensis* (GCA_030323885.1), *Brassicogethes aeneus* (GCA_921294245.1), *Brenthis daphne* (GCA_947093115.1), *Brenthis hecate* (GCA_951806755.1), *Brenthis ino* (GCA_921882275.1), *Brienomyrus brachyistius* (GCF_023856365.1, GCA_023856365.1), *Bruchidius siliquastri* (GCA_949316355.1), *Brugia pahangi* (GCA_012070555.1), *Buathra laborator* (GCA_934046635.1), *Bubalus bubalis* (GCF_003121395.1, GCF_019923925.1, GCA_003121395.1, GCA_019923925.1), *Bubalus carabanensis* (GCA_029407905.2, GCF_029407905.1), *Bucorvus abyssinicus* (GCA_009769605.1), *Budorcas taxicolor* (GCF_023091745.1, GCA_023091745.2), *Bufo bufo* (GCA_905171765.1, GCF_905171765.1), *Bufo gargarizans* (GCF_014858855.1, GCA_014858855.1), *Bufotes viridis* (GCA_033119425.1), *Bugulina stolonifera* (GCA_935421135.1), *Bungarus multicinctus* (GCA_023653725.1), *Bursaphelenchus mucronatus* (GCA_025436335.1), *Cabera pusaria* (GCA_954871355.1), *Caenorhabditis afra* (GCA_963570955.1), *Caenorhabditis briggsae* (GCA_021491975.1, GCA_022453885.1, GCA_029581135.1, GCF_000004555.2, GCA_000004555.3), *Caenorhabditis doughertyi* (GCA_963572265.1), *Caenorhabditis drosophilae* (GCA_963572285.1), *Caenorhabditis elegans* (GCA_020450165.1, GCA_004526295.1, GCA_029748435.1, GCA_000975215.1, GCA_022984815.1, GCA_028201515.1), *Caenorhabditis imperialis* (GCA_963572205.1), *Caenorhabditis inopinata* (GCA_003052745.1), *Caenorhabditis japonica* (GCA_963572235.1), *Caenorhabditis latens* (GCA_002259235.3), *Caenorhabditis nigoni* (GCA_002742825.1, GCA_027920645.1, GCA_001643685.2), *Caenorhabditis remanei* (GCF_010183535.1, GCA_001643735.4, GCA_002259225.3, GCA_010183535.1), *Caenorhabditis sp. 8 KK-2011* (GCA_963572245.1), *Caenorhabditis tropicalis* (GCA_016735795.1), *Caenorhabditis uteleia* (GCA_963573275.1), *Cairina moschata* (GCA_018104995.1), *Cairina moschata domestica* (GCA_009194515.1), *Calamotropha paludella* (GCA_927399485.1), *Caligus rogercresseyi* (GCA_013387185.1), *Callimorpha dominula* (GCA_949752705.1), *Callinectes sapidus* (GCA_020233015.1), *Calliopum simillimum* (GCA_951812925.1), *Calliphora vicina* (GCA_958450345.1), *Calliphora vomitoria* (GCA_942486065.2), *Callitettix versicolor* (GCA_022606455.1), *Callithrix jacchus* (GCF_011100555.1, GCF_009663435.1, GCA_011078405.1, GCA_013138085.1, GCA_011100555.2, GCA_009663435.2, GCA_000004665.1, GCF_000004665.1, GCA_011100535.2), *Calypte anna* (GCA_003957555.2, GCF_003957555.1), *Camarhynchus parvulus* (GCA_902806625.1, GCA_901933205.1, GCF_901933205.1), *Camelus dromedarius* (GCA_000803125.3, GCF_000803125.2), *Camelus ferus* (GCF_009834535.1, GCA_009834535.1), *Campaea margaritaria* (GCA_912999815.1), *Campoletis raptor* (GCA_948107755.1), *Camptogramma bilineatum* (GCA_958496255.1), *Canis lupus* (GCA_905319855.2), *Canis lupus dingo* (GCF_003254725.2, GCA_012295265.2, GCA_003254725.2), *Canis lupus familiaris* (GCA_000002285.4, GCA_004886185.2, GCF_014441545.1, GCF_011100685.1, GCF_005444595.1, GCF_000002285.5, GCA_008641055.3, GCA_031010295.1, GCA_011100685.1, GCA_014441545.1, GCA_031010635.1, GCA_012044875.1, GCA_005444595.1, GCA_012045015.1, GCA_031010765.1, GCA_031165255.1, GCA_013276365.2), *Cantharis flavilabris* (GCA_949152465.1), *Cantharis lateralis* (GCA_963170105.1), *Cantharis nigra* (GCA_958510845.1), *Cantharis rufa* (GCA_947369205.1), *Cantharis rustica* (GCA_911387805.1), *Capitulum mitella* (GCA_030062745.1), *Capra aegagrus* (GCA_000978405.1), *Capra hircus* (GCF_001704415.2, GCA_000317765.2, GCA_026652205.1, GCA_001704415.2, GCA_004361675.1, GCA_015443085.1), *Capricornis sumatraensis* (GCA_032405125.1), *Caprimulgus europaeus* (GCA_907165065.1), *Carabus problematicus* (GCA_963422195.1), *Caradrina clavipalpis* (GCA_932526535.1), *Caradrina kadenii* (GCA_947462355.1), *Carassius auratus* (GCA_003368295.1, GCA_014332655.1, GCA_019720715.2, GCF_003368295.1), *Carassius carassius* (GCA_963082965.1, GCF_963082965.1), *Carassius gibelio* (GCA_023724075.1, GCA_019843895.2), *Carcharodon carcharias* (GCA_017639515.1, GCF_017639515.1), *Carcina quercana* (GCA_910589575.2), *Carcinoscorpius rotundicauda* (GCA_011833715.1), *Caretta caretta* (GCF_023653815.1, GCA_023653815.1), *Carettochelys insculpta* (GCA_033958435.1), *Cariama cristata* (GCA_009819825.1), *Carpatolechia fugitivella* (GCA_951230895.1), *Carposina sasakii* (GCA_014607495.2), *Carterocephalus palaemon* (GCA_944567765.1), *Cataglyphis hispanica* (GCA_021464445.1, GCA_021464435.1, GCF_021464435.1), *Catalaphyllia jardinei* (GCA_022496165.2), *Catharus ustulatus* (GCA_009819885.2, GCF_009819885.2), *Catocala fraxini* (GCA_930367265.1), *Catocala nupta* (GCA_963675205.1), *Catocala sponsa* (GCA_963564715.1), *Catoptria pinella* (GCA_963556745.1), *Celastrina argiolus* (GCA_905187575.2), *Centropristis striata* (GCA_030273125.1, GCF_030273125.1), *Ceraclea dissimilis* (GCA_963576895.1), *Cerastis leucographa* (GCA_963082945.1), *Cerastis rubricosa* (GCA_949152445.1), *Cerastoderma edule* (GCA_947846245.1), *Ceratotherium simum cottoni* (GCA_021442165.1), *Ceratotherium simum simum* (GCA_023653735.1), *Cerceris rybyensis* (GCA_910591515.1), *Cercopithecus mitis* (GCA_028627265.1), *Ceriagrion tenellum* (GCA_963169105.1), *Certhia americana* (GCA_018697195.1), *Cervus canadensis* (GCF_019320065.1, GCA_019320065.1), *Cervus elaphus* (GCA_910594005.1, GCF_910594005.1), *Cervus elaphus hippelaphus* (GCA_002197005.1), *Cervus hanglu yarkandensis* (GCA_010411085.1), *Cervus nippon* (GCA_034195675.1), *Cetonia aurata* (GCA_949128085.1), *Ceutorhynchus assimilis* (GCA_917834065.1), *Chaenocephalus aceratus* (GCA_023974075.1), *Chaetogeoica ovagalla* (GCA_032441825.1), *Chalcis sispes* (GCA_949987625.1), *Chalcosyrphus nemorum* (GCA_949716465.1), *Channa argus* (GCA_018997905.1, GCA_033026475.1, GCA_004786185.1), *Channa maculata* (GCA_020496755.1), *Channa striata* (GCA_033026295.1), *Channallabes apus* (GCA_030522415.1), *Chanodichthys erythropterus* (GCA_024489055.1), *Chanos chanos* (GCA_902362185.1, GCF_902362185.1), *Charanyca ferruginea* (GCA_947361185.1), *Cheilinus undulatus* (GCA_018320785.1, GCF_018320785.1), *Cheilosia grossa* (GCA_963082955.1), *Cheilosia impressa* (GCA_948293265.1), *Cheilosia pagana* (GCA_936431705.1), *Cheilosia scutellata* (GCA_955612985.1), *Cheilosia soror* (GCA_949372485.1, GCA_948107745.1), *Cheilosia urbana* (GCA_946477585.1), *Cheilosia variabilis* (GCA_951230905.1), *Cheilosia vernalis* (GCA_949126925.1), *Cheilosia vulpina* (GCA_916610125.1), *Chelidonichthys spinosus* (GCA_029853015.1), *Chelmon rostratus* (GCA_017976325.1, GCF_017976325.1), *Chelon labrosus* (GCA_963514085.1), *Chelonia mydas* (GCF_015237465.2, GCA_015237465.2), *Chelonus formosanus* (GCA_028641665.1), *Cherax quadricarinatus* (GCA_026875155.2), *Chesias legatella* (GCA_947359385.1), *Cheumatopsyche charites* (GCA_024721215.1), *Chilo suppressalis* (GCA_902850365.2), *Chiloscyllium plagiosum* (GCF_004010195.1, GCA_004010195.1), *Chionomys nivalis* (GCF_950005125.1, GCA_950005125.1), *Chironomus riparius* (GCA_917627325.4), *Chironomus striatipennis* (GCA_026123125.1), *Chironomus tentans* (GCA_963573255.1), *Chiroxiphia lanceolata* (GCA_009829145.1, GCF_009829145.1), *Chlorocebus sabaeus* (GCA_000409795.2, GCF_000409795.2), *Chloroclysta siterata* (GCA_932294275.1), *Chloroclystis v-ata* (GCA_963691955.1), *Chlorops oryzae* (GCA_020466095.1), *Choloepus didactylus* (GCF_015220235.1, GCA_015220235.1), *Chondrosia reniformis* (GCA_947172415.1), *Choreutis nemorana* (GCA_949316135.1), *Chorisops tibialis* (GCA_963669355.1), *Choristoneura fumiferana* (GCA_025370935.1), *Chroicocephalus ridibundus* (GCA_030820635.1), *Chrysemys picta bellii* (GCA_000241765.5, GCA_011386835.2, GCF_000241765.5), *Chrysocyon brachyurus* (GCA_028533335.1), *Chrysodeixis includens* (GCA_903961255.1, GCA_941860345.1), *Chrysolina americana* (GCA_958502065.1), *Chrysolina haemoptera* (GCA_958298965.1), *Chrysolina oricalcia* (GCA_944452925.2), *Chrysomallon squamiferum* (GCA_012295275.1), *Chrysomela aeneicollis* (GCA_027562985.1), *Chrysopa pallens* (GCA_020423425.1), *Chrysoperla carnea* (GCA_905475395.1, GCF_905475395.1), *Chrysoteuchia culmella* (GCA_910589605.1), *Chrysotoxum bicinctum* (GCA_911387755.1), *Chydorus sphaericus* (GCA_030141595.1), *Chymomyza fuscimana* (GCA_949987675.1), *Ciconia maguari* (GCA_017639555.1), *Cinclus cinclus* (GCA_963662255.1), *Ciona intestinalis* (GCF_000224145.3, GCA_000224145.2), *Cirrhinus molitorella* (GCA_033026305.1), *Cistogaster globosa* (GCA_937654795.1), *Clarias fuscus* (GCA_030347435.1), *Clarias gariepinus* (GCA_024256465.1, GCF_024256425.1, GCA_024256435.1, GCA_024256425.2), *Clavelina lepadiformis* (GCA_947623445.1), *Clepsis dumicolana* (GCA_963691665.1), *Clistopyga incitator* (GCA_947507545.1), *Cloeon dipterum* (GCA_949628265.1), *Clonorchis sinensis* (GCA_003604175.2), *Clostera curtula* (GCA_905475355.2), *Clupea harengus* (GCA_900700415.2, GCF_900700415.2), *Clusia tigrina* (GCA_920105625.2), *Cnaphalocrocis medinalis* (GCA_014851415.1), *Coccinella septempunctata* (GCA_907165205.1, GCF_907165205.1), *Coelioxys conoideus* (GCA_947623535.1), *Coelopa frigida* (GCA_017309665.1), *Coelopa pilipes* (GCA_947389925.1), *Coendou prehensilis* (GCA_028534165.1), *Coenobita brevimanus* (GCA_032717465.1), *Coenonympha glycerion* (GCA_963855885.1), *Coilia nasus* (GCA_027475355.1, GCA_007927625.1), *Colaptes auratus* (GCA_015227895.2), *Coleophora deauratella* (GCA_958295455.1), *Coleophora flavipennella* (GCA_947284805.1), *Colias croceus* (GCA_905220415.1, GCF_905220415.1), *Colius striatus* (GCA_028858725.1, GCA_028858625.1), *Collichthys lucidus* (GCA_004119915.2), *Colobus guereza* (GCA_021498455.1, GCA_030247045.1), *Coloeus monedula* (GCA_013407035.1), *Cololabis saira* (GCA_033807715.1), *Colostygia pectinataria* (GCA_951394395.1), *Columba livia* (GCA_001887795.1, GCA_028654425.1), *Conchocele bisecta* (GCA_029237695.1), *Conger conger* (GCF_963514075.1, GCA_029692045.1, GCA_963514075.1), *Congeria kusceri* (GCA_027627225.1), *Conistra vaccinii* (GCA_948150665.1), *Conogethes punctiferalis* (GCA_031163375.1), *Conomelus anceps* (GCA_948455865.1), *Conops quadrifasciatus* (GCA_949752815.1), *Conus canariensis* (GCA_033310375.1), *Conus ventricosus* (GCA_018398815.1), *Cordilura impudica* (GCA_963682025.1), *Cordylochernes scorpioides* (GCA_030710605.1), *Coregonus clupeaformis* (GCA_020615455.1, GCA_018398675.1, GCF_018398675.1, GCF_020615455.1), *Coregonus sp. ‘balchen’* (GCA_902810595.1), *Coregonus ussuriensis* (GCA_031479575.1), *Corella eumyota* (GCA_963082875.1), *Coremacera marginata* (GCA_914767935.1), *Coreoperca whiteheadi* (GCA_011952105.1), *Coronaproctus castanopsis* (GCA_032883995.1), *Corticium candelabrum* (GCA_963422355.1), *Corvus cornix cornix* (GCA_000738735.6, GCF_000738735.6), *Corvus hawaiiensis* (GCF_020740725.1, GCA_020740725.1), *Corvus moneduloides* (GCA_009650955.1, GCF_009650955.1), *Corythoichthys intestinalis* (GCF_030265065.1, GCA_030265065.1), *Cosmia pyralina* (GCA_946251885.1), *Cosmia trapezina* (GCA_905163495.3), *Cosmorhoe ocellata* (GCA_963675405.1), *Cotesia glomerata* (GCF_020080835.1, GCA_020080835.1), *Cottoperca gobio* (GCA_900634415.1, GCF_900634415.1), *Cottus gobio* (GCA_023566465.1), *Coturnix japonica* (GCF_001577835.2, GCA_001577835.2), *Crambus lathoniellus* (GCA_949710035.1), *Craniophora ligustri* (GCA_905163465.1), *Cranoglanis bouderius* (GCA_026119655.1, GCA_023630585.1), *Crassostrea angulata* (GCA_025612915.2, GCF_025612915.1, GCA_025765675.3), *Crassostrea ariakensis* (GCA_020458035.1, GCA_020567875.1), *Crassostrea gigas* (GCA_011032805.1, GCF_902806645.1, GCA_902806645.1, GCA_025765685.3), *Crassostrea hongkongensis* (GCA_015776775.1), *Crassostrea nippona* (GCA_033439105.1), *Crataerina pallida* (GCA_949710015.1), *Crepidodera aurea* (GCA_949320105.2), *Cricetulus griseus* (GCA_003668045.2, GCF_003668045.3), *Crioceris asparagi* (GCA_958507055.1), *Criorhina berberina* (GCA_917880715.2), *Criorhina ranunculi* (GCA_951813785.1), *Crocallis elinguaria* (GCA_907269065.1), *Cromileptes altivelis* (GCA_013133815.1, GCA_019925165.1), *Crossocerus cetratus* (GCA_963675795.1), *Crotalus viridis viridis* (GCA_003400415.2), *Cryptocephalus moraei* (GCA_946251905.1), *Cryptocephalus primarius* (GCA_963576515.1), *Cryptophagus acutangulus* (GCA_963556235.1), *Cryptoprocta ferox* (GCA_028646485.1), *Cryptosula pallasiana* (GCA_945261195.1), *Ctenicera cuprea* (GCA_958336395.1), *Ctenopharyngodon idella* (GCA_019924925.1, GCF_019924925.1), *Cuculus canorus* (GCF_017976375.1, GCA_017976375.1), *Culex pipiens molestus* (GCA_024516115.1), *Culex pipiens pallens* (GCF_016801865.2, GCA_016801865.2), *Culex quinquefasciatus* (GCA_015732765.1, GCF_015732765.1), *Cyaniris semiargus* (GCA_905187585.1), *Cybosia mesomella* (GCA_946251805.1), *Cyclophora albipunctata* (GCA_963082685.1), *Cyclophora punctaria* (GCA_951394245.1), *Cyclopterus lumpus* (GCA_963457625.1, GCF_009769545.1, GCA_009769545.1), *Cyclura pinguis* (GCA_030412105.1), *Cydalima perspectalis* (GCA_951394215.1), *Cydia amplana* (GCA_948474715.1), *Cydia fagiglandana* (GCA_963556715.1), *Cydia pomonella* (GCA_033807575.1), *Cydia splendana* (GCA_910591565.2), *Cydia strobilella* (GCA_947568885.1), *Cygnus atratus* (GCA_013377495.2, GCF_013377495.2), *Cygnus olor* (GCF_009769625.2, GCA_009769625.2), *Cynaeus angustus* (GCA_030157275.1), *Cynocephalus volans* (GCA_027409185.1), *Cynoglossus semilaevis* (GCA_000523025.1, GCF_000523025.1), *Cynopterus sphinx* (GCA_030015415.1), *Cyprinodon diabolis* (GCA_030533445.1), *Cyprinodon nevadensis mionectes* (GCA_030533455.1), *Cyprinus carpio* (GCA_018340385.1, GCA_000951615.2, GCF_018340385.1, GCF_000951615.1, GCA_027406505.1), *Cyrtorhinus lividipennis* (GCA_019603395.1), *Dallia pectoralis* (GCA_029465975.1), *Dama dama* (GCA_033118175.1, GCF_033118175.1), *Danaus plexippus* (GCA_018135715.1, GCF_018135715.1), *Danaus plexippus plexippus* (GCF_009731565.1, GCA_009731565.1), *Danio aesculapii* (GCF_903798145.1, GCA_903798145.1), *Danio kyathit* (GCA_903798195.1), *Danio rerio* (GCA_903798175.1, GCA_000002035.4, GCA_903798165.1, GCA_020184715.1, GCA_018400075.1, GCA_903684855.2, GCA_008692375.1, GCA_903684865.1, GCF_000002035.6, GCA_903798185.1, GCA_944039275.1), *Daphnia carinata* (GCF_022539665.2, GCA_022539665.4), *Daphnia galeata* (GCA_030770115.1), *Daphnia magna* (GCA_030254905.1, GCF_020631705.1, GCA_003990815.1, GCF_003990815.1, GCA_020631705.2), *Daphnia pulex* (GCA_021134715.1, GCA_028752575.1, GCA_023526725.1, GCA_028752595.1, GCF_021134715.1), *Daphnia pulicaria* (GCA_021234035.2, GCF_021234035.1, GCA_017493165.1, GCA_021234015.2), *Daphnia sinensis* (GCA_013167095.2), *Dascillus cervinus* (GCA_949768715.1), *Dascyllus trimaculatus* (GCA_024666655.1), *Dastarcus helophoroides* (GCA_028583605.1), *Dasypolia templi* (GCA_963555695.1), *Dasypus novemcinctus* (GCA_030445055.1, GCA_030445035.1, GCF_030445035.1), *Dasysyrphus albostriatus* (GCA_946251815.1), *Dasyurus viverrinus* (GCA_028643805.1, GCA_020854095.1), *Decapterus maruadsi* (GCA_030347415.2), *Deilephila elpenor* (GCA_949752805.1), *Deilephila porcellus* (GCA_905220455.2), *Delia radicum* (GCA_021234595.1), *Delphinus delphis* (GCA_949987515.1, GCF_949987515.1, GCA_030062865.1), *Dendrolimus kikuchii* (GCA_019925095.2), *Dendrolimus pini* (GCA_949752895.1), *Dendrolimus punctatus* (GCA_012273795.1), *Dendropsophus ebraccatus* (GCA_027789765.1), *Denticeps clupeoides* (GCA_900700375.1, GCF_900700375.1, GCA_900700375.2), *Dermacentor silvarum* (GCA_013339745.2, GCF_013339745.2), *Dermatophagoides farinae* (GCA_024713945.1), *Dermochelys coriacea* (GCA_009764565.4, GCF_009764565.3), *Deroplatys truncata* (GCA_030765065.1), *Desmodus rotundus* (GCA_022682495.1, GCF_022682495.1, GCA_022682395.1), *Diabrotica balteata* (GCA_918026665.1), *Diabrotica virgifera virgifera* (GCF_917563875.1, GCA_917563875.2), *Diachrysia chrysitis* (GCA_932294365.1), *Diadema setosum* (GCA_033980235.1), *Diadumene lineata* (GCA_918843875.1), *Diaperis boleti* (GCA_963583935.1), *Diaphora mendica* (GCA_949125395.1), *Diaphorina citri* (GCA_030643865.1), *Diarsia brunnea* (GCA_949774965.1), *Diarsia dahlii* (GCA_949775195.1), *Diarsia mendica* (GCA_949316265.1), *Diarsia rubi* (GCA_932274075.1), *Diatraea saccharalis* (GCA_918026875.4), *Diceros bicornis* (GCA_028533375.1), *Diceros bicornis minor* (GCA_020826845.1, GCF_020826845.1), *Dicrocoelium dendriticum* (GCA_944474145.2), *Dicycla oo* (GCA_948252095.1), *Dicyrtomina minuta* (GCA_949802685.1), *Diglossa brunneiventris* (GCA_019023105.1), *Diloba caeruleocephala* (GCA_947459985.1), *Dilophus febrilis* (GCA_958336335.1), *Diogma glabrata* (GCA_963693315.1), *Diorhabda carinata* (GCA_029229535.1), *Diorhabda carinulata* (GCA_026250575.1, GCF_026250575.1), *Diorhabda elongata* (GCA_026230145.1), *Diorhabda sublineata* (GCA_026230105.1, GCF_026230105.1), *Diplazon laetatorius* (GCA_963662125.1), *Diplosoma virens* (GCA_963680785.1), *Diprion similis* (GCA_021155765.1, GCF_021155765.1), *Dircenna loreta* (GCA_963555665.1), *Discoglossus pictus* (GCA_027410445.1), *Dissostichus eleginoides* (GCA_031216635.1), *Ditula angustiorana* (GCA_963691745.1), *Dolichopus griseipennis* (GCA_963082915.1), *Dolichovespula media* (GCA_911387685.1), *Dolichovespula saxonica* (GCA_911387935.1), *Dolichovespula sylvestris* (GCA_918808275.2), *Dolomedes plantarius* (GCA_907164885.2), *Dorcus hopei* (GCA_033060865.1), *Dorcus parallelipipedus* (GCA_958336345.1), *Doryrhamphus excisus* (GCF_030265055.1, GCA_030265055.1), *Doryteuthis pealeii* (GCA_023376005.1), *Dreissena polymorpha* (GCA_020536995.1, GCF_020536995.1), *Drepana falcataria* (GCA_945859725.1), *Drepanosiphum platanoidis* (GCA_948098885.1), *Dromaius novaehollandiae* (GCA_016128335.2), *Dromiciops gliroides* (GCA_019393635.1, GCF_019393635.1), *Drosophila albomicans* (GCF_009650485.2, GCA_009650485.2), *Drosophila americana* (GCA_030788265.1), *Drosophila ananassae* (GCA_017639315.2, GCA_030555595.1, GCF_017639315.1), *Drosophila athabasca* (GCA_003185025.1, GCA_008121215.1, GCA_008121225.1), *Drosophila biarmipes* (GCA_025231255.1, GCF_025231255.1), *Drosophila bifasciata* (GCA_009664405.1), *Drosophila bipectinata* (GCA_030179905.1), *Drosophila busckii* (GCF_001277935.1, GCA_011750605.1, GCA_001277935.1, GCF_011750605.1), *Drosophila elegans* (GCA_011057505.1), *Drosophila funebris* (GCA_958295475.1), *Drosophila gunungcola* (GCA_025200985.1, GCF_025200985.1), *Drosophila histrio* (GCA_958299025.2), *Drosophila immigrans* (GCA_963583835.1, GCA_963584065.1), *Drosophila innubila* (GCA_004354385.2, GCF_004354385.1, GCA_004354385.1), *Drosophila kepulauana* (GCA_023558455.1), *Drosophila kikkawai* (GCA_030179895.1), *Drosophila lowei* (GCA_008121275.1), *Drosophila mauritiana* (GCA_004382145.1, GCF_004382145.1), *Drosophila melanogaster* (GCA_029775775.1, GCA_020141795.1, GCA_029775175.1, GCA_029774715.1, GCA_029774615.1, GCA_029774635.1, GCA_029775755.1, GCA_029774655.1, GCA_029774675.1, GCA_029774695.1, GCA_029775815.1, GCA_020142005.1, GCA_029775195.1, GCF_000001215.4, GCA_020141985.1, GCA_020141835.1, GCA_029775795.1, GCA_020142055.1, GCA_020142045.1, GCA_020141815.1, GCA_020142025.1, GCA_020141765.1, GCA_020141845.1, GCA_020141935.1, GCA_024500395.1, GCA_029774375.1, GCA_029774395.1, GCA_020142105.1, GCA_020142085.1, GCA_029774415.1, GCA_029774435.1, GCA_029774455.1, GCA_029775475.1, GCA_029774475.1, GCA_002300595.1, GCA_002310755.1, GCA_002310775.1, GCA_003397115.2, GCA_003401685.1, GCA_003401735.1, GCA_003401745.1, GCA_003401795.1, GCA_000001215.4, GCA_003401805.1, GCA_020169495.1, GCA_029775115.1, GCA_029775135.1, GCA_020141485.1, GCA_020141745.1, GCA_003401925.1, GCA_003401975.1, GCA_003402005.1, GCA_020141585.1, GCA_020141575.1, GCA_020141625.1, GCA_020141515.1, GCA_015832445.1, GCA_029775155.1, GCA_015852585.1, GCA_020141505.1, GCA_020141495.1, GCA_004798055.1, GCA_004798075.2, GCA_020141925.1, GCA_003402015.1, GCA_003401855.1, GCA_020141595.1, GCA_003402055.1, GCA_029775675.1, GCA_029775275.1, GCA_029775655.1, GCA_029775295.1, GCA_029775635.1, GCA_029775315.1, GCA_029775615.1, GCA_029775335.1, GCA_020141665.1, GCA_029775355.1, GCA_020141855.1, GCA_029775375.1, GCA_029775555.1, GCA_029775535.1, GCA_029775515.1, GCA_029775395.1, GCA_020141875.1, GCA_029775415.1, GCA_029775495.1, GCA_029775215.1, GCA_029775715.1, GCA_029775735.1, GCA_020141675.1, GCA_020141705.1, GCA_003401885.1, GCA_029775255.1, GCA_029775695.1, GCA_029775575.1, GCA_020141655.1, GCA_029775095.1, GCA_020141735.1, GCA_029775595.1, GCA_003401915.1, GCA_020141955.1, GCA_029775235.1, GCA_029775455.1), *Drosophila miranda* (GCF_003369915.1, GCA_003369915.2, GCF_000269505.1, GCA_000269505.2), *Drosophila nasuta* (GCA_023558535.1, GCF_023558535.1), *Drosophila nebulosa* (GCA_024703675.1), *Drosophila novamexicana* (GCA_030788195.1), *Drosophila obscura* (GCA_963584055.1, GCA_963583975.1), *Drosophila pallidifrons* (GCA_023558445.1), *Drosophila persimilis* (GCA_020698555.1), *Drosophila phalerata* (GCA_951394115.2), *Drosophila pseudoobscura* (GCF_009870125.1, GCA_004329205.1, GCA_009870125.2), *Drosophila pseudoobscura pseudoobscura* (GCA_000001765.3, GCA_020698565.1), *Drosophila santomea* (GCA_016746245.2, GCF_016746245.2), *Drosophila sechellia* (GCF_004382195.2, GCA_004382195.2), *Drosophila simulans* (GCA_029774515.1, GCA_029774535.1, GCA_016746395.2, GCA_029774575.1, GCA_029774995.1, GCA_029775075.1, GCA_029774955.1, GCA_029775035.1, GCA_029775015.1, GCA_029774895.1, GCA_029775055.1, GCA_029774915.1, GCA_000259055.1, GCF_000259055.1, GCA_029774975.1, GCA_029774595.1, GCA_000259045.1, GCF_016746395.2, GCA_029774875.1, GCF_000754195.2, GCA_029774495.1, GCA_029774935.1, GCA_029774555.1, GCA_029774735.1, GCA_004382185.1, GCA_000754195.3, GCA_029774835.1, GCA_029774815.1, GCA_029774795.1, GCA_000820565.1, GCA_029774755.1, GCA_029774855.1, GCA_029774775.1), *Drosophila subobscura* (GCF_008121235.1, GCA_008121235.1, GCA_963583985.1, GCA_963584095.1), *Drosophila subpulchrella* (GCA_014743375.2, GCF_014743375.2), *Drosophila sulfurigaster albostrigata* (GCA_023558435.1), *Drosophila sulfurigaster bilimbata* (GCA_023558465.1), *Drosophila sulfurigaster sulfurigaster* (GCA_023558475.1), *Drosophila suzukii* (GCF_013340165.1, GCA_013340165.1), *Drosophila takahashii* (GCA_030179915.1), *Drosophila teissieri* (GCA_016746235.2, GCF_016746235.2), *Drosophila triauraria* (GCA_014170255.2), *Drosophila virilis* (GCA_007989325.2, GCA_030788295.1, GCA_000004125.1), *Drosophila willistoni* (GCF_018902025.1, GCA_018902025.2), *Drosophila yakuba* (GCA_000005975.1, GCF_016746365.2, GCA_016746335.2, GCF_000005975.2, GCA_016746365.2), *Dryas iulia moderata* (GCA_019049465.1), *Drymonia ruficornis* (GCA_947859195.1), *Dryobates pubescens* (GCA_014839835.1, GCF_014839835.1), *Dryobota labecula* (GCA_947523025.1), *Dryobotodes eremita* (GCA_917490735.1), *Dryococelus australis* (GCA_029891345.1), *Dryomyza anilis* (GCA_951804985.1), *Dugong dugon* (GCA_030035585.1), *Dunckerocampus dactyliophorus* (GCA_027744805.2, GCF_027744805.1), *Dysmachus trigonus* (GCA_949715965.1), *Dyspetes luteomarginatus* (GCA_963669185.1), *Ecdyonurus torrentis* (GCA_949318235.1), *Echeneis naucrates* (GCA_900963305.2, GCF_900963305.1, GCA_031770045.1, GCA_900963305.1), *Echiichthys vipera* (GCA_963691815.1), *Echinococcus granulosus* (GCA_021556725.1), *Echinometra lucunter* (GCA_029962745.1), *Ecliptopera silaceata* (GCA_932527185.1), *Ectemnius continuus* (GCA_910591665.1), *Ectemnius lituratus* (GCA_910593735.2), *Ectropis crepuscularia* (GCA_963693475.1), *Ectropis grisescens* (GCA_017562165.1), *Eidolon dupreanum* (GCA_028627145.1), *Eilema caniola* (GCA_949126895.1), *Eilema depressum* (GCA_914767945.1), *Eilema griseolum* (GCA_963662185.1), *Eilema sororculum* (GCA_914829495.1), *Electrona antarctica* (GCA_951216825.1), *Electrophaes corylata* (GCA_947095575.1), *Electrophorus electricus* (GCA_013358815.1, GCF_013358815.1), *Elegia similella* (GCA_947532085.1), *Elephas maximus* (GCA_033060105.1, GCA_032718585.1, GCA_032718755.1), *Elephas maximus indicus* (GCF_024166365.1, GCA_024166365.1), *Eleutherodactylus coqui* (GCA_019857665.1), *Elmis aenea* (GCA_947652605.1), *Elophila nymphaeata* (GCA_955850985.1), *Elysia crispata* (GCA_963854125.1), *Emmelina monodactyla* (GCA_916618145.1), *Empis stercorea* (GCA_949752835.1), *Emys orbicularis* (GCA_028017835.1), *Endomychus coccineus* (GCA_958510875.1), *Endotricha flammealis* (GCA_905163395.2), *Engystomops pustulosus* (GCA_019512145.1), *Ennomos erosarius* (GCA_963580505.1), *Ennomos fuscantarius* (GCA_905220475.3), *Ennomos quercinarius* (GCA_910589525.2), *Entelurus aequoreus* (GCA_033978785.1), *Entomobrya proxima* (GCA_029691765.1), *Entosphenus tridentatus* (GCA_014621495.2), *Ephestia elutella* (GCA_018467065.1), *Epiblema foenella* (GCA_963556455.1), *Epicampocera succincta* (GCA_932526305.1), *Epinephelus cyanopodus* (GCA_026686955.1), *Epinephelus fuscoguttatus* (GCF_011397635.1, GCA_011397635.1), *Epinephelus lanceolatus* (GCA_005281545.1, GCF_005281545.1, GCA_017165655.1), *Epinephelus moara* (GCA_006386435.1, GCF_006386435.1), *Epinotia bilunana* (GCA_947049275.1), *Epinotia brunnichana* (GCA_963854355.1), *Epinotia demarniana* (GCA_945867215.1), *Epinotia nisella* (GCA_932294315.1), *Epinotia ramella* (GCA_947578815.1), *Epirrhoe alternata* (GCA_963565295.1), *Epirrhoe tristata* (GCA_951394285.1), *Epirrita christyi* (GCA_951392215.1), *Epistrophe eligans* (GCA_951394125.1), *Epistrophe grossulariae* (GCA_929447395.1), *Epistrophella euchroma* (GCA_947049315.1), *Episyrphus balteatus* (GCA_945859705.1, GCF_945859705.1), *Erinaceus europaeus* (GCA_950295315.1, GCF_950295315.1), *Eriocheir sinensis* (GCA_013436485.1, GCA_024679095.1, GCA_023348025.1, GCF_024679095.1, GCA_033459325.1), *Eriosoma lanigerum* (GCA_013282895.1), *Eristalinus aeneus* (GCA_955652365.1), *Eristalinus sepulchralis* (GCA_944738645.1), *Eristalis arbustorum* (GCA_916610145.1), *Eristalis pertinax* (GCA_907269125.1), *Eristalis tenax* (GCA_905231855.2), *Erithacus rubecula* (GCA_903797595.2), *Erpetoichthys calabaricus* (GCA_900747795.4, GCF_900747795.2), *Erynnis tages* (GCA_905147235.1), *Erythrolamprus reginae* (GCA_031021105.1), *Eschrichtius robustus* (GCA_028021215.1), *Esox lucius* (GCF_000721915.3, GCA_011004845.1, GCF_004634155.1, GCF_011004845.1, GCA_004634155.1, GCA_000721915.3), *Esperia sulphurella* (GCA_947086405.1), *Etheostoma cragini* (GCF_013103735.1, GCA_013103735.1), *Etheostoma nigrum* (GCA_029931835.1), *Etheostoma perlongum* (GCA_026937815.1), *Etheostoma spectabile* (GCA_008692095.1, GCF_008692095.1), *Ethmia dodecea* (GCA_963855545.1), *Eubalaena glacialis* (GCA_028564815.2, GCF_028564815.1, GCA_028571275.2), *Eubasilissa splendida* (GCA_031772225.1), *Eublepharis macularius* (GCF_028583425.1, GCA_028583425.1), *Euclidia mi* (GCA_944739405.2), *Eucosma cana* (GCA_951800055.1), *Eudemis profundana* (GCA_947034925.1), *Eudonia lacustrata* (GCA_947562085.1), *Eudonia mercurella* (GCA_963082485.1), *Eudonia truncicolella* (GCA_949315975.1), *Eueides isabella* (GCA_019049475.1), *Eugnorisma glareosa* (GCA_947578425.1), *Eulemur mongoz* (GCA_028534055.1), *Euleptes europaea* (GCF_029931775.1, GCA_029931775.1), *Eulithis prunata* (GCA_918843925.1), *Eulithis testata* (GCA_947507515.1), *Eumerus sabulonum* (GCA_951905685.1), *Eupeodes corollae* (GCA_945859685.1, GCF_945859685.1), *Eupeodes latifasciatus* (GCA_920104205.1), *Eupeodes luniger* (GCA_951509635.1), *Eupithecia abbreviata* (GCA_943735965.1), *Eupithecia centaureata* (GCA_944547425.1), *Eupithecia dodoneata* (GCA_947044415.1), *Eupithecia exiguata* (GCA_947086465.1), *Eupithecia insigniata* (GCA_947859395.1), *Eupithecia inturbata* (GCA_963662085.1), *Eupithecia subfuscata* (GCA_963564075.1), *Eupithecia subumbrata* (GCA_949316285.1), *Eupithecia tripunctaria* (GCA_955876795.1), *Eupithecia vulgata* (GCA_946478455.1), *Euplexia lucipara* (GCA_921972225.1), *Euproctis similis* (GCA_905147225.2), *Euprymna scolopes* (GCA_024364805.1), *Eupsilia transversa* (GCA_914767815.1), *Eurois occulta* (GCA_950022335.1), *Euspilapteryx auroguttella* (GCA_951802225.1), *Eustalomyia histrio* (GCA_949748255.1), *Euthynnus affinis* (GCA_029490765.1), *Eutrigla gurnardus* (GCA_963514095.1), *Euzophera pinguis* (GCA_947363495.1), *Exephanes ischioxanthus* (GCA_958510785.1), *Fabriciana adippe* (GCA_905404265.1), *Falcaria lacertinaria* (GCA_951449985.1), *Falco biarmicus* (GCA_023638135.1, GCF_023638135.1), *Falco cherrug* (GCA_023634085.1, GCF_023634085.1), *Falco naumanni* (GCA_017639655.1, GCF_017639655.2), *Falco peregrinus* (GCF_023634155.1, GCA_023634155.1, GCA_001887755.1), *Falco punctatus* (GCA_963210335.1), *Falco rusticolus* (GCF_015220075.1, GCA_015220075.1), *Felis catus* (GCA_000003115.1, GCA_016509815.2, GCF_018350175.1, GCA_018350175.1, GCA_000181335.6), *Felis chaus* (GCA_019924945.1), *Felis nigripes* (GCA_028533295.1, GCA_032458615.1), *Ferdinandea cuprea* (GCA_963576555.1), *Ficedula albicollis* (GCF_000247815.1, GCA_000247815.2), *Fissipunctia ypsillon* (GCA_947568875.1), *Folsomia candida* (GCA_020920555.1, GCA_020923455.1), *Formica selysi* (GCA_009859135.1), *Fragum fragum* (GCA_946902895.1), *Fragum sueziense* (GCA_963680895.1), *Fragum whitleyi* (GCA_948146395.1), *Frankliniella intonsa* (GCA_033675135.1), *Fringilla coelebs* (GCA_963513975.1), *Fringilla coelebs coelebs* (GCA_015532645.2), *Fundulus heteroclitus* (GCF_011125445.2, GCA_011125445.2), *Furcifer pardalis* (GCA_030440675.1), *Furcula furcula* (GCA_911728495.1), *Gadus chalcogrammus* (GCF_026213295.1, GCA_026213295.1), *Gadus macrocephalus* (GCA_031193875.1, GCA_031168955.1, GCF_031168955.1), *Gadus morhua* (GCA_902167405.1, GCF_902167405.1, GCA_010882105.1), *Galaxea fascicularis* (GCA_948470475.1), *Galleria mellonella* (GCA_958496185.1), *Gallus gallus* (GCA_024653025.1, GCA_024652985.1, GCA_025370635.1, GCA_024653045.1, GCA_016700215.2, GCF_016699485.2, GCF_016700215.2, GCA_002798355.1, GCA_024206055.2, GCA_030979905.1, GCA_000002315.5, GCA_027557775.1, GCA_016699485.1, GCA_030914275.1, GCA_030849555.1, GCF_000002315.6, GCA_027408225.1, GCA_024653035.1, GCA_024652995.1, GCA_030914265.1), *Gallus gallus gallus* (GCA_033088195.1), *Gambusia affinis* (GCA_019740435.1, GCF_019740435.1), *Gandaritis pyraliata* (GCA_947859175.1), *Gari tellinella* (GCA_922989275.2), *Gasterosteus aculeatus* (GCA_017751045.1), *Gasterosteus aculeatus aculeatus* (GCA_016920845.1, GCF_016920845.1), *Gasterosteus nipponicus* (GCA_014132575.2), *Gasteruption jaculator* (GCA_949825005.1), *Gastracanthus pulcherrimus* (GCA_949152435.1), *Gastrophryne carolinensis* (GCA_027917425.1, GCA_027917415.1), *Gastrophysa polygoni* (GCA_963576655.1), *Gavia stellata* (GCF_030936135.1, GCA_030936125.1, GCA_030936135.1), *Gavialis gangeticus* (GCA_030020295.1), *Geothlypis trichas* (GCA_009764595.1), *Geotrupes spiniger* (GCA_959613385.1), *Geotrypetes seraphini* (GCA_902459505.2, GCA_902459505.1, GCF_902459505.1), *Germaria angustata* (GCA_963681545.1), *Gerris lacustris* (GCA_951217055.1), *Gibbula magus* (GCA_936450465.1), *Gigantopelta aegis* (GCA_016097555.1, GCF_016097555.1), *Giraffa camelopardalis rothschildi* (GCA_017591445.1), *Giraffa tippelskirchi* (GCA_013496395.1), *Girardinichthys multiradiatus* (GCA_021462225.2, GCF_021462225.1), *Glaucopsyche alexis* (GCA_905404095.1), *Globia sparganii* (GCA_949316385.1), *Globicephala melas* (GCF_963455315.1, GCA_963455315.1), *Glyphotaelius pellucidus* (GCA_936435175.1), *Gnatocerus cornutus* (GCA_029298725.1), *Gobio gobio* (GCA_949357685.1, GCA_021464655.1), *Gobiocypris rarus* (GCA_023029165.1, GCA_018491645.1), *Gobius niger* (GCA_951799975.1), *Gonocerus acuteangulatus* (GCA_946811695.1), *Gopherus evgoodei* (GCA_007399415.1, GCF_007399415.2), *Gopherus flavomarginatus* (GCA_025201925.1, GCF_025201925.1), *Gordionus sp. m RMFG-2023* (GCA_954871325.1), *Gorilla gorilla gorilla* (GCA_028885475.1, GCF_029281585.1, GCA_900006655.3, GCF_000151905.2, GCA_008122165.1, GCA_000151905.3, GCF_008122165.1, GCA_028885495.1, GCA_029281585.1), *Gorsachius magnificus* (GCA_034008185.1), *Gortyna flavago* (GCA_963669345.1), *Gouania willdenowi* (GCA_900634775.1, GCF_900634775.1, GCA_900634775.2), *Gracilinanus agilis* (GCF_016433145.1, GCA_016433145.1), *Grampus griseus* (GCA_028646425.1), *Grapholita molesta* (GCA_022674325.1), *Griposia aprilina* (GCA_916610205.1), *Grus americana* (GCA_028858595.1, GCF_028858705.1, GCA_028858705.1), *Gymnocephalus cernua* (GCA_023631565.1), *Gymnocheta viridis* (GCA_956483585.1), *Gymnocypris eckloni scoliostomus* (GCA_027564155.1), *Gymnogyps californianus* (GCF_018139145.2, GCA_018139145.2), *Gymnoscelis rufifasciata* (GCA_929108375.1), *Gymnoscopelus braueri* (GCA_963280865.1), *Gymnoscopelus microlampas* (GCA_963454915.1), *Gymnosoma rotundatum* (GCA_916610165.2), *Gymnothorax javanicus* (GCA_029692085.1), *Gymnothorax reevesii* (GCA_029721435.1), *Gypaetus barbatus* (GCA_028022735.1), *Habrosyne pyritoides* (GCA_907165245.1), *Haemaphysalis longicornis* (GCA_013339765.2), *Haemonchus contortus* (GCA_000469685.2, GCA_007637855.2), *Haemorhous mexicanus* (GCA_027477595.1, GCF_027477595.1), *Haliaeetus albicilla* (GCA_947461875.1), *Halichondria panicea* (GCA_963675165.1), *Haliclystus octoradiatus* (GCA_916610825.1), *Halictus ligatus* (GCA_028454255.1), *Halictus quadricinctus* (GCA_028454245.1), *Halictus rubicundus* (GCA_028454235.1), *Halyzia sedecimguttata* (GCA_937662695.2), *Harmonia axyridis* (GCA_914767665.1, GCF_914767665.1, GCA_011033045.2), *Harmothoe impar* (GCA_947462335.1), *Harpalus rubripes* (GCA_963082715.1), *Harpalus rufipes* (GCA_951394225.1), *Harpia harpyja* (GCA_026419915.1, GCF_026419915.1), *Hecatera dysodea* (GCA_905332915.2), *Hedya salicella* (GCA_905404275.2), *Helarctos malayanus* (GCA_028533245.1), *Heliconius charithonia* (GCA_030704555.1), *Heliconius sara* (GCA_917862395.2), *Helicoverpa armigera* (GCF_023701775.1, GCA_023701775.1, GCA_026262555.1), *Helicoverpa assulta* (GCA_029618815.1), *Helicoverpa zea* (GCA_022343045.1, GCF_022581195.2, GCA_022581195.1), *Heligmosomoides bakeri* (GCA_947359475.1), *Heligmosomoides polygyrus polygyrus* (GCA_947396885.1), *Heliocidaris erythrogramma* (GCA_025617745.1), *Heliocidaris tuberculata* (GCA_025618425.1), *Heliothis peltigera* (GCA_958496145.1), *Hemaris fuciformis* (GCA_907164795.1), *Hemibagrus wyckioides* (GCA_019097595.1, GCF_019097595.1), *Hemicordylus capensis* (GCF_027244095.1, GCA_027244095.1), *Hemicrepidius niger* (GCA_963082805.1), *Hemiprocne comata* (GCA_020745705.1), *Hemiscyllium ocellatum* (GCA_020745735.1, GCF_020745735.1), *Hemistola chrysoprasaria* (GCA_947063395.1), *Hemithea aestivaria* (GCA_947507615.1), *Hepsetus odoe* (GCA_017165825.1), *Hermaeophaga mercurialis* (GCA_951812935.1), *Hermetia illucens* (GCF_905115235.1, GCA_905115235.1), *Herminia tarsipennalis* (GCA_945859575.2), *Hesperia comma* (GCA_905404135.1), *Hestina assimilis* (GCA_023373885.1), *Heterobilharzia americana* (GCA_944470555.2, GCA_944470545.2), *Heterocephalus glaber* (GCA_944319715.1, GCA_944319725.1), *Heterodera glycines* (GCA_004148225.2), *Heterohyrax brucei* (GCA_028571685.1), *Heteromyza rotundicornis* (GCA_951394025.1), *Heteronetta atricapilla* (GCA_011075105.1), *Heteronotia binoei* (GCA_032191835.1, GCF_032191835.1), *Heteropelma amictum* (GCA_959613375.1), *Heterotis niloticus* (GCA_018136845.1), *Hexactinellida sp. DTS-2022* (GCA_028753945.1), *Hexaprotodon liberiensis* (GCA_023065765.1), *Himalopsyche anomala* (GCA_031772345.1), *Hipparchia semele* (GCA_933228805.2), *Hippocampus abdominalis* (GCA_018466805.1), *Hippocampus zosterae* (GCA_026261805.1, GCF_025434085.1, GCA_025434085.3), *Hippoglossus hippoglossus* (GCF_009819705.1, GCA_009819705.1), *Hippoglossus stenolepis* (GCF_022539355.2, GCA_022539355.2), *Hippopotamus amphibius* (GCA_023065835.1), *Hippopotamus amphibius kiboko* (GCF_030028045.1, GCA_030028045.1), *Hippopus hippopus* (GCA_946811185.1), *Hipposideros larvatus* (GCA_031876335.1), *Hirtodrosophila cameraria* (GCA_949708635.1), *Hirundo rustica* (GCA_015227805.3, GCF_015227805.2), *Hofmannophila pseudospretella* (GCA_947369225.1), *Hololepta plana* (GCA_963695495.1), *Holothuria leucospilota* (GCA_029531755.1), *Holotrichia oblita* (GCA_023690525.1), *Homalodisca vitripennis* (GCF_021130785.1, GCA_021130785.2), *Homo sapiens* (GCA_016695395.2, GCA_948473225.1, GCA_000002125.2, GCA_948474725.1, GCA_021951015.1, GCA_016700455.2, GCA_001292825.2, GCA_003634875.1, GCA_000365445.1, GCA_948473255.1, GCA_022833125.2, GCA_001712695.1, GCA_024586135.1, GCA_002180035.3, GCA_000002115.2, GCA_000001405.29, GCA_948473215.1, GCA_020497115.1, GCA_001524155.4, GCA_000306695.2, GCA_021950905.1, GCA_000002135.3, GCA_948473235.1, GCA_000212995.1, GCA_000252825.1, GCA_948474675.1, GCA_000442335.2, GCA_948474735.1, GCA_015476435.1, GCA_016894425.1, GCA_002077035.3, GCF_000001405.40, GCA_018852605.2, GCF_000002125.1, GCF_000306695.2, GCA_018852615.2, GCA_018873775.2, GCA_020497085.1, GCA_020881995.2, GCA_011064465.2, GCA_014905855.1), *Hoplias malabaricus* (GCA_029633875.1), *Hoplodrina ambigua* (GCA_949774945.1), *Hoplodrina blanda* (GCA_949316365.1), *Horisme vitalbata* (GCA_951804965.1), *Hormaphis cornu* (GCA_017140985.1), *Hormiphora californensis* (GCA_020137815.1), *Hyalomma asiaticum* (GCA_013339685.2), *Hydra vulgaris* (GCA_022113875.1, GCA_024232925.1, GCF_022113875.1), *Hydractinia symbiolongicarpus* (GCF_029227915.1, GCA_029227915.2), *Hydraecia micacea* (GCA_914767645.1), *Hydriomena furcata* (GCA_912999785.1), *Hydrochoerus hydrochaeris* (GCA_015741225.1), *Hydrophis curtus* (GCA_019472885.1), *Hydrophis cyanocinctus* (GCA_019473425.1), *Hydrophis major* (GCA_033807585.1), *Hydropotes inermis* (GCA_020226075.1), *Hydrotaea cyrtoneurina* (GCA_958296145.1), *Hydrotaea diabolus* (GCA_963513945.1), *Hyla sarda* (GCA_029499605.1, GCF_029499605.1), *Hylaea fasciaria* (GCA_905147375.1), *Hylaeus volcanicus* (GCA_026283585.1, GCF_026283585.1), *Hyles euphorbiae* (GCA_023078785.2), *Hyles vespertilio* (GCA_009982885.2), *Hylobates pileatus* (GCA_021498465.1), *Hylyphantes graminicola* (GCA_023701765.1), *Hymenochirus boettgeri* (GCA_019447015.1), *Hymenolepis microstoma* (GCA_000469805.3), *Hymenopus coronatus* (GCA_030762935.1), *Hypanus sabinus* (GCF_030144855.1, GCA_030144855.1, GCA_030144785.1), *Hypena proboscidalis* (GCA_905147285.1), *Hypercompe scribonia* (GCA_949316085.1), *Hyperoodon ampullatus* (GCA_949752795.1), *Hyperoplus immaculatus* (GCA_949357725.1), *Hyperoplus lanceolatus* (GCA_026929865.2), *Hypomecis punctinalis* (GCA_949316475.1), *Hypomesus transpacificus* (GCF_021917145.1, GCA_021917145.1, GCA_021870715.1), *Hypophthalmichthys molitrix* (GCA_022817975.1), *Hypophthalmichthys nobilis* (GCA_019925145.1), *Hyppa rectilinea* (GCA_951799385.1), *Hypsopygia costalis* (GCA_937001555.2), *Icerya purchasi* (GCA_952773005.1), *Ichneumon xanthorius* (GCA_917499995.1), *Ichthyophis bannanicus* (GCA_033557465.1), *Ictalurus furcatus* (GCA_023701845.2, GCF_023375685.1, GCA_023375685.2), *Ictalurus punctatus* (GCF_001660625.3, GCA_001660625.3, GCA_004006655.3), *Ictidomys tridecemlineatus* (GCA_016881025.1, GCF_016881025.1), *Idaea aversata* (GCA_907269075.1), *Idaea biselata* (GCA_958496205.1), *Idaea dimidiata* (GCA_949358125.1), *Idaea straminata* (GCA_951213275.1), *Ilyophis sp. 1 JC-2022* (GCA_022702515.1), *Ilyophis sp. 2 JC-2022* (GCA_022702525.1), *Ilyophis sp. JC_2022_1* (GCA_022702415.1), *Incurvaria masculella* (GCA_946894095.1), *Indicator indicator* (GCA_027791375.2, GCF_027791375.1), *Iphiclides podalirius* (GCA_933534255.1), *Ips nitidus* (GCA_018691245.2), *Ischnura elegans* (GCF_921293095.1, GCA_921293095.2, GCA_921293095.1), *Isoperla grammatica* (GCA_945910005.1), *Istiophorus platypterus* (GCA_016859345.1), *Ixodes scapularis* (GCA_031841145.1), *Jaculus jaculus* (GCA_020740685.1, GCF_020740685.1), *Junco hyemalis* (GCA_003829775.2), *Karalla daura* (GCA_029224185.1), *Kogia breviceps* (GCF_026419965.1, GCA_026419965.1, GCA_026419985.1), *Kryptolebias hermaphroditus* (GCA_007896545.1), *Kryptolebias marmoratus* (GCF_001649575.2, GCA_001649575.2), *Labeo rohita* (GCA_022985175.1, GCF_022985175.1), *Labia minor* (GCA_963082975.1), *Labroides dimidiatus* (GCA_030710495.1), *Labrus mixtus* (GCA_963584025.1, GCF_963584025.1), *Lacanobia oleracea* (GCA_950371165.1), *Lacanobia wlatinum* (GCA_947578705.1), *Lacerta agilis* (GCF_009819535.1, GCA_009819535.1), *Lagenorhynchus albirostris* (GCF_949774975.1, GCA_949774975.1), *Lagopus muta* (GCA_023343835.1, GCF_023343835.1), *Lagorchestes hirsutus* (GCA_028533205.1), *Lagria hirta* (GCA_947359425.1), *Lama glama* (GCA_028534125.1), *Lamellibrachia columna* (GCA_963662155.1), *Lampris incognitus* (GCA_029633865.1, GCF_029633865.1, GCA_022059245.1), *Lampropteryx suffumata* (GCA_948098915.1), *Lamprotornis superbus* (GCA_015883425.2), *Laodelphax striatellus* (GCA_014465815.1, GCA_017141395.1), *Laothoe populi* (GCA_905220505.1), *Lapara coniferarum* (GCA_949316025.1), *Larimichthys crocea* (GCA_000972845.2, GCA_004352675.2, GCA_003845795.1, GCA_003711585.2, GCF_000972845.2, GCA_900246015.1), *Lasioglossum calceatum* (GCA_028455575.1), *Lasioglossum figueresi* (GCA_028455805.1), *Lasioglossum lativentre* (GCA_916610255.1), *Lasioglossum leucozonium* (GCA_028454225.1), *Lasioglossum malachurum* (GCA_028455605.1), *Lasioglossum marginatum* (GCA_028455795.1), *Lasioglossum morio* (GCA_916610235.2), *Lasioglossum oenotherae* (GCA_028455765.1), *Lasioglossum pauxillum* (GCA_028455745.1, GCA_933228785.1), *Lasioglossum vierecki* (GCA_028455595.1), *Lasioglossum zephyrum* (GCA_028455615.1), *Lasiommata megera* (GCA_928268935.1), *Lasius fuliginosus* (GCA_949152495.1), *Laspeyria flexula* (GCA_905147015.1), *Lateolabrax maculatus* (GCA_004023545.1, GCA_004028665.1, GCA_031216445.1), *Lates calcarifer* (GCA_001640805.2, GCF_001640805.2), *Latheticus oryzae* (GCA_030157265.1), *Lathronympha strigana* (GCA_949128165.1), *Latrodectus elegans* (GCA_030067965.1), *Leguminivora glycinivorella* (GCA_023078275.1, GCF_023078275.1), *Leistus spinibarbis* (GCA_933228885.1), *Lemur catta* (GCA_020740605.1, GCF_020740605.2), *Leontopithecus rosalia* (GCA_028533165.1), *Leopardus geoffroyi* (GCF_018350155.1, GCA_018350155.1), *Lepeophtheirus nordmannii* (GCA_963680855.1), *Lepeophtheirus salmonis* (GCA_016086655.3, GCA_905330665.1, GCF_016086655.3, GCA_902860245.1, GCA_016086655.4), *Lepidonotus clava* (GCA_936440205.1), *Lepisosteus oculatus* (GCA_000242695.1, GCF_000242695.1), *Leptidea sinapis* (GCA_905404315.1), *Leptinotarsa decemlineata* (GCA_024712935.1), *Leptobrachium ailaonicum* (GCA_032062155.1, GCA_018994145.1), *Leptobrachium leishanense* (GCA_009667805.1), *Leptodactylus fuscus* (GCA_031893055.1, GCA_031893025.1), *Leptodirus hochenwartii* (GCA_947310635.1), *Leptogaster cylindrica* (GCA_963082835.1), *Leptophobia aripa* (GCA_951799465.1), *Leptopilina boulardi* (GCA_032872485.1), *Leptopilina drosophilae* (GCA_032873175.1), *Leptopilina heterotoma* (GCA_032872495.1), *Leptopilina sp. ZJUHJH005* (GCA_032872475.1), *Leptopilina syphax* (GCA_032872505.1), *Leptopterna dolabrata* (GCA_954871275.1), *Leptura quadrifasciata* (GCA_963675555.1), *Lepturacanthus savala* (GCA_030544235.1), *Lepus europaeus* (GCA_033115175.1), *Lepus timidus* (GCA_009760805.1), *Lethenteron reissneri* (GCF_015708825.1, GCA_015708825.1), *Leucania comma* (GCA_958295575.1), *Leucaspius delineatus* (GCA_023718395.1), *Leuciscus idus* (GCA_021554675.1), *Leucoma salicis* (GCA_948253155.1), *Leucophora obtusa* (GCA_949987735.1), *Leucoraja erinacea* (GCA_028641065.1, GCF_028641065.1), *Leucozona laternaria* (GCA_932273885.2), *Leuctra nigra* (GCA_934045905.1), *Liasis olivaceus* (GCA_030867105.1, GCA_030867085.1), *Lichenostomus cassidix* (GCA_008360975.2), *Ligdia adustata* (GCA_947049295.1), *Limanda limanda* (GCF_963576545.1, GCA_963576545.1), *Limenitis camilla* (GCA_905147385.1), *Limnephilus auricula* (GCA_951813805.1), *Limnephilus lunatus* (GCA_917563855.2), *Limnephilus marmoratus* (GCA_917880885.1), *Limnephilus rhombicus* (GCA_929108145.2), *Limnoperna fortunei* (GCA_944474755.1), *Lineus longissimus* (GCA_910592395.2), *Linnaemya tessellans* (GCA_951800035.1), *Linnaemya vulpina* (GCA_963675445.1), *Lipophrys pholis* (GCA_963383615.1), *Liposcelis brunnea* (GCA_023512825.1), *Lithobates sylvaticus* (GCA_028564925.1), *Lithophane leautieri* (GCA_949152455.1), *Lithophane ornitopus* (GCA_948473465.1), *Lithophane semibrunnea* (GCA_947363905.1), *Lithophane socia* (GCA_947522985.1), *Lithosia quadra* (GCA_963576445.1), *Litomosoides sigmodontis* (GCA_963070105.1), *Lobophora halterata* (GCA_932526365.1), *Lochmaea capreae* (GCA_949126875.1), *Lochmaea crataegi* (GCA_947563755.1), *Locusta migratoria* (GCA_026315105.1), *Lomographa bimaculata* (GCA_948107665.1), *Lonchura striata domestica* (GCF_005870125.1, GCA_005870125.1), *Lophura swinhoii* (GCA_030408155.1), *Loxodonta africana* (GCA_033060095.1, GCA_032717415.1, GCA_030014295.1, GCA_032717405.1), *Lucilia cuprina* (GCA_022045245.1, GCF_022045245.1), *Lucinisca nassula* (GCA_963580285.1), *Luffia ferchaultella* (GCA_949709985.1), *Luidia sarsii* (GCA_949987565.1), *Lumbricus rubellus* (GCA_945859605.1), *Lumbricus terrestris* (GCA_949752735.1), *Luperina nickerlii* (GCA_963855955.1), *Luperina testacea* (GCA_927399505.1), *Lutjanus erythropterus* (GCA_020091685.1), *Lutra lutra* (GCF_902655055.1, GCA_902655055.2), *Lutzomyia longipalpis* (GCA_024334085.1, GCF_024334085.1), *Lycaena phlaeas* (GCA_905333005.2), *Lycaon pictus* (GCA_001883655.1, GCA_001887905.1), *Lycia hirtaria* (GCA_947563715.1), *Lycodopsis pacificus* (GCA_028022725.1), *Lycophotia porphyrea* (GCA_950005105.1), *Lymantria dispar* (GCA_963576585.1, GCA_032191425.1), *Lymantria monacha* (GCA_905163515.2), *Lynx canadensis* (GCA_007474595.2, GCF_007474595.2), *Lypha dubia* (GCA_947311025.1), *Lysandra bellargus* (GCA_905333045.1), *Lysandra coridon* (GCA_905220515.1), *Lytechinus pictus* (GCF_015342785.2, GCA_015342785.2), *Lytechinus variegatus* (GCA_018143015.1, GCF_018143015.1), *Macaca cyclopis* (GCA_026956025.1), *Macaca fascicularis* (GCF_000364345.1, GCA_011100615.1, GCF_012559485.2, GCA_000222185.1, GCA_000230815.1, GCA_012559485.3, GCA_000364345.1), *Macaca mulatta* (GCA_008058575.1, GCA_003339765.3, GCF_003339765.1, GCA_001270425.1, GCA_000230795.1, GCA_000002255.2, GCF_000772875.2, GCF_000002255.3, GCA_000772875.3), *Macaca thibetana thibetana* (GCA_024542745.1, GCF_024542745.1), *Macaria notata* (GCA_927399415.1), *Machimus atricapillus* (GCA_933228815.1), *Machimus rusticus* (GCA_951509405.1), *Macrobrachium nipponense* (GCA_015104395.1, GCA_015110555.1), *Macrochelys suwanniensis* (GCA_033349115.1, GCA_033296515.1), *Macrophya alboannulata* (GCA_949628255.1), *Macropis europaea* (GCA_916610135.2), *Macropodus opercularis* (GCA_030770545.1), *Macropus fuliginosus* (GCA_028583105.1), *Macropus giganteus* (GCA_028627215.1), *Mactra quadrangularis* (GCA_025267735.1), *Magallana gigas* (GCA_963853765.1), *Malachius bipustulatus* (GCA_910589415.1), *Malaclemys terrapin pileata* (GCA_027887205.1, GCF_027887155.1, GCA_027887155.1), *Malacosoma neustria* (GCA_963693495.1), *Malaya genurostris* (GCF_030247185.1, GCA_030247185.2), *Malthinus flaveolus* (GCA_950108345.1), *Malurus cyaneus samueli* (GCA_009741485.1), *Mamestra brassicae* (GCA_905163435.1), *Manduca sexta* (GCF_014839805.1, GCA_014839805.1), *Maniola hyperantus* (GCF_902806685.1, GCA_902806685.1, GCA_963576595.1, GCA_902806685.2), *Maniola jurtina* (GCA_905333055.1, GCF_905333055.1), *Manis pentadactyla* (GCA_030020395.1, GCF_030020395.1), *Mantis religiosa* (GCA_030765055.1), *Marasmarcha lunaedactyla* (GCA_923062675.1), *Marasmia exigua* (GCA_019059595.1), *Martes flavigula* (GCA_029410595.1), *Martes martes* (GCA_963455335.1), *Marthasterias glacialis* (GCA_911728455.2), *Mastacembelus armatus* (GCA_900324485.3, GCA_019455535.1, GCA_019455525.1, GCA_900324485.2, GCF_900324485.2), *Mastomys coucha* (GCF_008632895.1, GCA_008632895.1), *Matsumurasca onukii* (GCA_018831715.1), *Mauremys mutica* (GCF_020497125.1, GCA_020497125.1, GCA_032357905.1), *Mauremys reevesii* (GCA_016161935.1, GCF_016161935.1), *Maylandia zebra* (GCF_000238955.4, GCA_000238955.5), *Meandrina meandrites* (GCA_963693305.1), *Mechanitis mazaeus* (GCA_959347395.1), *Mechanitis messenoides* (GCA_959347415.1), *Meconema thalassinum* (GCA_946902985.2), *Mecyna flavalis* (GCA_949319885.1), *Meda fulgida* (GCA_030578275.1), *Megachile leachella* (GCA_963576845.1), *Megachile ligniseca* (GCA_945859555.1), *Megachile willughbiella* (GCA_945859595.2), *Megaleporinus macrocephalus* (GCA_021613375.1), *Megalobrama amblycephala* (GCF_018812025.1, GCA_018812025.1), *Megalops atlanticus* (GCA_019176425.1), *Megalops cyprinoides* (GCF_013368585.1, GCA_013368585.1), *Megalurothrips usitatus* (GCA_026979955.1), *Megamerina dolium* (GCA_963854835.1), *Meganola albula* (GCA_936450015.1), *Meghimatium bilineatum* (GCA_034231615.1), *Meiosimyza decipiens* (GCA_963680825.1), *Meiosimyza platycephala* (GCA_963662105.1), *Melanargia galathea* (GCA_920104075.1), *Melanchra persicariae* (GCA_947386135.1), *Melangyna quadrimaculata* (GCA_949320155.1), *Melanophora roralis* (GCA_963583895.1), *Melanostigma gelatinosum* (GCA_949748355.1), *Melanostoma mellinum* (GCA_914767635.1), *Melanostoma scalare* (GCA_949752695.1), *Melanotaenia boesemani* (GCA_017639745.1, GCF_017639745.1), *Melanotaenia duboulayi* (GCA_026261665.1), *Melanotus villosus* (GCA_963082815.1), *Meleagris gallopavo* (GCA_000146605.4, GCA_943295565.1, GCF_000146605.3, GCA_943294205.1), *Meles meles* (GCF_922984935.1, GCA_922984935.2, GCA_922984935.1), *Melieria crassipennis* (GCA_963668005.1), *Melinaea marsaeus rileyi* (GCA_918358865.1), *Melinaea menophilus n. ssp. AW-2005* (GCA_918358695.1), *Meliscaeva auricollis* (GCA_948107695.1), *Melitaea cinxia* (GCF_905220565.1, GCA_905220565.1), *Mellicta athalia* (GCA_905220545.2), *Melolontha melolontha* (GCA_935421215.2), *Melopsittacus undulatus* (GCA_012275295.1, GCF_012275295.1), *Melospiza georgiana* (GCA_028018845.1, GCF_028018845.1), *Membranipora membranacea* (GCA_914767715.1), *Mercenaria mercenaria* (GCA_021730395.1, GCA_014805675.2, GCF_021730395.1, GCF_014805675.1), *Meriones unguiculatus* (GCA_030254825.1, GCF_030254825.1), *Merodon equestris* (GCA_958301585.1), *Merops nubicus* (GCA_009819595.1), *Merzomyia westermanni* (GCA_949987695.1), *Mesoligia furuncula* (GCA_916614155.1), *Mesoplodon densirostris* (GCF_025265405.1, GCA_025265405.1), *Mesopsocus fuscifrons* (GCA_950004255.1), *Meta bourneti* (GCA_933210815.1), *Metalampra italica* (GCA_949699065.1), *Metallyticus violacea* (GCA_030762175.1), *Metaphire vulgaris* (GCA_018105865.1), *Metellina segmentata* (GCA_947359465.1), *Metopia argyrocephala* (GCA_963576795.1), *Metopolophium dirhodum* (GCF_019925205.1, GCA_019925205.1), *Metridium senile* (GCA_949775045.1), *Microcaecilia unicolor* (GCA_901765095.2, GCF_901765095.1, GCA_901765095.1), *Microcebus murinus* (GCF_000165445.2, GCA_000165445.3), *Microchirus variegatus* (GCA_963457635.1), *Microchrysa polita* (GCA_949715475.1), *Microctonus brassicae* (GCA_940306215.1, GCA_918226075.1), *Microplitis demolitor* (GCF_026212275.2, GCA_026212275.2), *Microplitis manilae* (GCA_029641195.1, GCA_030273425.1), *Microplitis mediator* (GCF_029852145.1, GCA_029852145.1), *Micropterix aruncella* (GCA_944548615.1), *Micropterus dolomieu* (GCA_021292245.1, GCF_021292245.1), *Micropterus salmoides* (GCA_022435785.1, GCA_019677235.1), *Microtus ochrogaster* (GCF_000317375.1, GCA_000317375.1), *Miltochrista miniata* (GCA_933228765.1), *Mimachlamys varia* (GCA_947623455.1), *Mimas tiliae* (GCA_905332985.1), *Mimumesa dahlbomi* (GCA_917499265.3), *Miopithecus talapoin* (GCA_028551445.1), *Mirounga angustirostris* (GCF_021288785.2, GCA_021288785.3), *Misgurnus anguillicaudatus* (GCF_027580225.1, GCA_027580225.1), *Mobula birostris* (GCA_030028105.1, GCA_030035685.1), *Molanna angustata* (GCA_963576475.1), *Molothrus ater* (GCF_012460135.2, GCA_012460135.3), *Monochamus saltuarius* (GCA_025584915.1), *Monodelphis domestica* (GCF_000002295.2, GCF_027887165.1, GCA_000002295.1, GCA_027887165.1), *Monomorium pharaonis* (GCA_013373865.2, GCF_013373865.1), *Monopis laevigella* (GCA_947458855.1, GCA_947359445.1), *Monoplex corrugatus* (GCA_030674185.1), *Montipora capitata* (GCA_949126865.1), *Morinoia aosen* (GCA_030386875.1), *Morus bassanus* (GCA_031468805.1, GCA_031468815.1), *Motacilla alba alba* (GCF_015832195.1, GCA_015832195.1), *Mugilogobius chulae* (GCA_016735935.1), *Mungos mungo* (GCA_028533875.1), *Muntiacus crinifrons* (GCA_020276665.1, GCA_020226055.1), *Muntiacus muntjak* (GCA_008782695.1), *Muntiacus reevesi* (GCA_020226045.1, GCA_008787405.2), *Muraenolepis orangiensis* (GCA_027704905.1), *Muricea muricata* (GCA_963855995.1), *Mus caroli* (GCA_900094665.2, GCF_900094665.2), *Mus musculus* (GCA_921999865.2, GCA_031761455.1, GCA_001624675.1, GCA_001624535.1, GCA_947599735.1, GCA_001632575.1, GCA_000001635.9, GCA_921998905.2, GCA_001624215.1, GCA_921998355.2, GCA_921998635.2, GCA_921998315.2, GCA_001632525.1, GCA_921998325.2, GCA_001624505.1, GCA_001632555.1, GCA_921998555.2, GCA_001624475.1, GCA_921997125.2, GCA_030265425.1, GCA_949316315.1, GCA_922000895.2, GCA_001632615.1, GCA_001624185.1, GCF_000001635.27, GCA_001624295.1, GCA_921997145.2, GCA_947593165.1, GCA_031763685.1, GCA_000002165.1, GCA_001624745.1), *Mus musculus castaneus* (GCA_921999005.2, GCA_001624445.1), *Mus musculus domesticus* (GCA_921998345.2, GCA_001624835.1), *Mus musculus molossinus* (GCA_921999095.2), *Mus musculus musculus* (GCA_921998335.2, GCA_001624775.1), *Mus pahari* (GCF_900095145.1, GCA_900095145.2), *Mus spretus* (GCA_921997135.2, GCA_001624865.1), *Muscardinus avellanarius* (GCA_963383645.1), *Muscidifurax raptorellus* (GCA_020010945.3), *Musotima nitidalis* (GCA_949126915.1), *Mustela erminea* (GCF_009829155.1, GCA_009829155.1), *Mustela lutreola* (GCF_030435805.1, GCA_030435805.1), *Mustela nigripes* (GCF_022355385.1, GCA_022355385.1), *Mya arenaria* (GCF_026914265.1, GCA_026914265.1), *Myathropa florea* (GCA_930367185.1), *Myiopsitta monachus* (GCA_017639245.1), *Myopa tessellatipennis* (GCA_943737955.2), *Myopa testacea* (GCA_949629155.1), *Myotis daubentonii* (GCF_963259705.1, GCA_963259705.1), *Myripristis murdjan* (GCF_902150065.1, GCA_902150065.1), *Myrrha octodecimguttata* (GCA_958510865.1), *Mystacides longicornis* (GCA_963576905.1), *Mythimna albipuncta* (GCA_929112965.1), *Mythimna ferrago* (GCA_910589285.2), *Mythimna impura* (GCA_905147345.3), *Mythimna lalbum* (GCA_949319445.1), *Mythimna loreyi* (GCA_029852875.1), *Mythimna pallens* (GCA_961205895.1), *Mythimna separata* (GCA_026898235.1, GCA_030763345.1, GCA_020882275.1, GCA_029852925.1), *Mythimna vitellina* (GCA_949316375.1), *Mytilisepta virgata* (GCA_028015205.1), *Mytilus coruscus* (GCA_017311375.1), *Mytilus edulis* (GCA_963676685.1, GCA_019925275.1, GCA_025215535.1, GCA_025276775.1), *Mytilus galloprovincialis* (GCA_025277285.1), *Myxocyprinus asiaticus* (GCA_019703515.2, GCF_019703515.2), *Myzus persicae* (GCA_029232275.1), *Naja naja* (GCA_009733165.1), *Napeogenes inachia* (GCA_959347355.1), *Napeogenes sylphis* (GCA_959347405.1), *Nasalis larvatus* (GCA_000772465.1), *Nasonia vitripennis* (GCF_000002325.3, GCA_009193385.2, GCF_009193385.2, GCA_000002325.2), *Nasua narica* (GCA_028533885.1), *Nebria brevicollis* (GCA_944738965.1), *Nebria salina* (GCA_944039245.1), *Necator americanus* (GCA_031761385.1), *Nematolebias whitei* (GCF_014905685.2, GCA_014905685.2), *Nematopogon swammerdamellus* (GCA_946902875.1), *Nematostella vectensis* (GCA_932526225.1, GCF_932526225.1, GCA_033964005.1), *Nemotelus nigrinus* (GCA_947369275.1), *Nemoura dubitans* (GCA_921293005.1), *Nemurella pictetii* (GCA_921293315.2), *Neoaliturus tenellus* (GCA_030545055.2), *Neoarius graeffei* (GCF_027579695.1, GCA_027579695.1), *Neoascia interrupta* (GCA_947623515.1), *Neoceratitis asiatica* (GCA_030068015.2), *Neoceratodus forsteri* (GCA_016271365.2), *Neocrepidodera transversa* (GCA_963243735.1), *Neodiprion fabricii* (GCF_021155785.1, GCA_021155785.1), *Neodiprion lecontei* (GCA_021901455.1, GCA_001263575.2, GCF_021901455.1), *Neodiprion pinetum* (GCF_021155775.1, GCA_021155775.1), *Neodiprion virginianus* (GCA_021901495.1, GCF_021901495.1), *Neofelis nebulosa* (GCF_028018385.1, GCA_028018385.1, GCA_030324275.1), *Neogale vison* (GCF_020171115.1, GCA_020171115.1), *Neoitamus cyanurus* (GCA_947538895.1), *Neolamprologus multifasciatus* (GCA_963576455.1), *Neomonachus schauinslandi* (GCA_002201575.2, GCF_002201575.2), *Neoneuromus ignobilis* (GCA_024320075.1, GCA_029203775.1), *Neosalanx taihuensis* (GCA_030340665.1), *Neostethus bicornis* (GCA_902685375.1), *Neotoxoptera formosana* (GCA_022818045.1), *Nephotettix cincticeps* (GCA_023375725.1), *Nephrocerus scutellatus* (GCA_947095585.1), *Nephrotoma appendiculata* (GCA_947310385.1), *Nephrotoma flavescens* (GCA_932526605.1), *Nephrotoma guestfalica* (GCA_963691935.1), *Neria commutata* (GCA_963457695.1), *Nerophis lumbriciformis* (GCA_033978685.1), *Nerophis ophidion* (GCA_033978795.1), *Nesoenas mayeri* (GCA_963082525.1), *Netelia dilatata* (GCA_946811545.1), *Netelia virgata* (GCA_951802705.1), *Nettapus auritus* (GCA_011076525.1), *Nezara viridula* (GCA_928085145.1), *Nibea albiflora* (GCA_902410095.1, GCA_014281875.1), *Nibea coibor* (GCA_023373845.1), *Nicrophorus investigator* (GCA_963457615.1), *Nilaparvata lugens* (GCA_014356525.1, GCA_015708395.1, GCF_014356525.2), *Nipponacmea schrenckii* (GCA_030562195.1), *Nippostrongylus brasiliensis* (GCA_030553155.1), *Noctua comes* (GCA_963082995.1), *Noctua fimbriata* (GCA_905163415.1), *Noctua janthe* (GCA_910589295.1), *Noctua janthina* (GCA_963576755.1), *Noctua pronuba* (GCA_905220335.1), *Nomada fabriciana* (GCA_907165295.1), *Nomada ferruginata* (GCA_963583965.1), *Nomada flava* (GCA_951802745.1), *Nomada fucata* (GCA_948146005.1), *Nomada hirtipes* (GCA_951802735.1), *Nomada lathburiana* (GCA_963667235.1), *Nomada panzeri* (GCA_951802685.1), *Nomada ruficornis* (GCA_951802695.1), *Nomascus leucogenys* (GCF_000146795.2, GCA_000146795.3, GCA_006542625.1, GCF_006542625.1), *Nomophila noctuella* (GCA_958496325.1), *Notamacropus eugenii* (GCA_028372415.1), *Nothobranchius furzeri* (GCA_014300015.1, GCF_001465895.1, GCA_001465895.2), *Notocelia uddmanniana* (GCA_905163555.1), *Notodonta dromedarius* (GCA_905147325.1), *Notodonta ziczac* (GCA_918843915.1), *Notolabrus celidotus* (GCA_009762535.1, GCF_009762535.1), *Notothenia rossii* (GCA_949606895.1), *Nowickia ferox* (GCA_936439885.1), *Nudaria mundana* (GCA_963556515.1), *Numida meleagris* (GCA_002078875.2, GCF_002078875.1), *Nycteola revayana* (GCA_947037095.2), *Nyctibius grandis* (GCA_013368605.1), *Nycticebus bengalensis* (GCA_023898255.1), *Nycticebus coucang* (GCF_027406575.1, GCA_027406575.1), *Nylanderia fulva* (GCA_024268025.1), *Nymphalis c-album* (GCA_963151315.1), *Nymphalis io* (GCA_905147045.1, GCF_905147045.1), *Nymphalis polychloros* (GCA_905220585.2), *Nymphalis urticae* (GCA_905147175.2), *Nymphula nitidulata* (GCA_947347705.1), *Nysson spinosus* (GCA_910591585.2), *Obolodiplosis robiniae* (GCA_028476595.1), *Ochlodes sylvanus* (GCA_905404295.2), *Ochotona princeps* (GCA_030435715.1, GCA_014633375.1, GCF_014633375.1, GCF_030435755.1, GCA_030435755.1), *Ochropacha duplaris* (GCA_951361185.1), *Ochropleura leucogaster* (GCA_958449745.1), *Ochropleura plecta* (GCA_905475445.1), *Octodonta nipae* (GCA_034190945.1), *Octopus bimaculoides* (GCA_001194135.2, GCF_001194135.2), *Octopus sinensis* (GCF_006345805.1, GCA_006345805.1), *Octopus vulgaris* (GCA_951406725.2), *Ocypus olens* (GCA_910593695.2), *Odocoileus hemionus* (GCA_020976825.1), *Odocoileus virginianus* (GCA_023699985.2), *Odontamblyopus lacepedii* (GCA_032888595.1), *Odontamblyopus rebecca* (GCA_030686955.1), *Odontesthes bonariensis* (GCA_027942865.1), *Odontocerum albicorne* (GCA_949825065.1), *Oedothorax gibbosus* (GCA_019343175.1), *Oegoconia quadripuncta* (GCA_949316235.1), *Oenanthe melanoleuca* (GCF_029582105.1, GCA_029582105.1), *Oikopleura dioica* (GCA_907165135.1), *Okapia johnstoni* (GCA_024291935.2), *Oligia fasciuncula* (GCA_963082905.1), *Oligia latruncula* (GCA_948474745.1), *Oligia strigilis* (GCA_951800025.1), *Omphaloscelis lunosa* (GCA_916610215.1), *Oncorhynchus gorbuscha* (GCA_021184085.1, GCA_017355495.1, GCF_021184085.1), *Oncorhynchus keta* (GCA_012931545.1, GCF_012931545.1, GCF_023373465.1, GCA_023373465.1), *Oncorhynchus kisutch* (GCF_002021735.2, GCA_002021735.2), *Oncorhynchus mykiss* (GCA_013265735.3, GCA_025558465.1, GCF_013265735.2, GCA_029834435.1, GCF_002163495.1, GCA_002163495.1, GCA_900005705.1), *Oncorhynchus nerka* (GCF_006149115.2, GCA_006149115.2), *Oncorhynchus tshawytscha* (GCA_002872995.1, GCF_002872995.1, GCA_002831465.1, GCA_018296145.1, GCF_018296145.1), *Onychomys arenicola* (GCA_949786405.1, GCA_949786425.1), *Onychomys leucogaster* (GCA_949786395.1, GCA_949786385.1), *Onychomys torridus* (GCA_949787125.1, GCF_903995425.1, GCA_903995425.1), *Onychostoma macrolepis* (GCA_012432095.1, GCF_012432095.1), *Ooceraea biroi* (GCA_003672135.1, GCF_003672135.1), *Operophtera brumata* (GCA_932527175.1), *Ophion costatus* (GCA_951751655.1), *Ophion ellenae* (GCA_963210295.1), *Ophion slaviceki* (GCA_944452715.1), *Ophonus ardosiacus* (GCA_943142095.1), *Opisthocomus hoazin* (GCA_030867145.1, GCA_030867165.1), *Opisthograptis luteolata* (GCA_931315375.2), *Orchestes rusci* (GCA_958502075.1), *Orcinus orca* (GCA_937001465.1, GCF_937001465.1), *Oreochromis aureus* (GCA_016625535.1, GCF_013358895.1, GCA_013358895.1), *Oreochromis niloticus* (GCF_001858045.2, GCA_000188235.2, GCF_000188235.2, GCA_001858045.3, GCA_013350305.1), *Orgyia antiqua* (GCA_916999025.1), *Ormia ochracea* (GCA_963402855.1), *Ornithorhynchus anatinus* (GCA_004115215.4, GCA_000002275.2, GCF_004115215.2), *Orthonevra nobilis* (GCA_963555765.1), *Orthosia gothica* (GCA_949775005.1), *Orthosia gracilis* (GCA_947562075.1), *Orthosia incerta* (GCA_948252205.1), *Oryctolagus cuniculus* (GCF_000003625.3, GCF_009806435.1, GCA_009806435.2, GCA_000003625.1), *Oryctolagus cuniculus cuniculus* (GCA_013371645.1), *Oryzias curvinotus* (GCA_023969325.1), *Oryzias javanicus* (GCA_003999625.1), *Oryzias latipes* (GCA_004348095.1, GCA_002234715.1, GCA_004347445.1, GCF_000313675.1, GCA_002234695.1, GCA_002234675.1, GCF_002234675.1, GCA_000313675.1, GCA_004348135.1), *Oryzias melastigma* (GCF_002922805.2, GCA_002922805.2), *Oscarella lobularis* (GCA_947507565.1), *Oscheius dolichura* (GCA_932521035.1), *Oscheius onirici* (GCA_932521025.1), *Oscheius sp. DF5120* (GCA_932521415.1), *Oscheius sp. JU1382* (GCA_932521405.1), *Oscheius tipulae* (GCA_013425905.1), *Osmerus eperlanus* (GCA_963692335.1), *Osmia bicornis bicornis* (GCA_907164935.1, GCF_907164935.1), *Ostrea edulis* (GCA_024362745.1, GCA_032173915.1, GCF_947568905.1, GCA_947568905.1), *Ostrinia nubilalis* (GCA_963855985.1), *Othius punctulatus* (GCA_951805005.1), *Otis tarda* (GCA_026413225.1), *Otocolobus manul* (GCA_028564725.2), *Ovis ammon polii* (GCA_028583565.1), *Ovis ammon polii x Ovis aries* (GCA_023701675.1), *Ovis aries* (GCF_002742125.1, GCF_016772045.2, GCA_022432835.1, GCA_022416775.1, GCA_002742125.1, GCA_011170295.1, GCA_016772045.2, GCA_017524585.1, GCA_018804185.1, GCA_019145175.1, GCF_000298735.2, GCA_024222265.1, GCA_030512445.1, GCA_024222175.1, GCA_022244695.1, GCA_022432845.1, GCA_022416725.1, GCA_022432825.1, GCA_022244705.1, GCA_022416685.1, GCA_022416695.1, GCA_022416745.1, GCA_022416755.1, GCA_022416785.1, GCA_022416915.1, GCA_033439445.1, GCA_022538005.1, GCA_000298735.2, GCA_000005525.1), *Ovis canadensis canadensis* (GCA_001039535.1), *Ovis orientalis* (GCA_014523465.1), *Oxytorus armatus* (GCA_958009045.1), *Oxyura jamaicensis* (GCA_011077185.1, GCF_011077185.1), *Pachypeltis micranthus* (GCA_020466155.1), *Pachypsylla venusta* (GCA_012654025.1), *Pachyrhynchus sulphureomaculatus* (GCA_019049505.1), *Pachyuromys duprasi* (GCA_028658305.1), *Paedocypris micromegethes* (GCA_028031935.1), *Paguma larvata* (GCA_030068075.1), *Palloptera scutellata* (GCA_958295655.1), *Pammene aurita* (GCA_947086415.1), *Pammene fasciana* (GCA_911728535.1), *Pan paniscus* (GCA_028858845.1, GCA_028858825.1, GCA_000258655.2, GCF_029289425.1, GCF_000258655.2, GCA_029289425.1, GCA_013052645.3), *Pan troglodytes* (GCF_000001515.7, GCA_028858805.1, GCF_028858775.1, GCA_000090855.1, GCA_000001515.5, GCA_002880755.3, GCA_028858775.1, GCF_002880755.1), *Pan troglodytes verus* (GCA_000002175.2, GCF_000002175.1), *Pandemis cinnamomeana* (GCA_932294345.1), *Pandemis corylana* (GCA_949127965.1), *Pandemis heparana* (GCA_963854515.1), *Pangasianodon gigas* (GCA_022758105.1), *Pangasianodon hypophthalmus* (GCA_027358585.1, GCF_009078355.1, GCA_009078355.1, GCA_016801045.1, GCF_027358585.1), *Pangasius djambal* (GCA_022985145.1), *Panopea generosa* (GCA_029582155.1), *Panorpa germanica* (GCA_963678705.1), *Pantala flavescens* (GCA_020796165.1), *Panthera leo* (GCA_008795835.1, GCF_018350215.1, GCA_018350215.1), *Panthera onca* (GCF_028533385.1, GCA_028533385.1), *Panthera tigris* (GCA_024034525.1, GCA_018350195.2, GCF_018350195.1), *Panthera tigris tigris* (GCA_021131075.1, GCA_021130815.1), *Panthera uncia* (GCF_023721935.1, GCA_023721935.1, GCA_028646445.1), *Panzeria rudis* (GCA_956483635.1), *Pao palembangensis* (GCA_015343265.1), *Papilio elwesi* (GCA_029849275.1, GCA_029641285.1), *Papilio machaon* (GCF_912999745.1, GCA_912999745.1), *Papio anubis* (GCA_008728515.2, GCF_000264685.3, GCA_008728515.1, GCF_008728515.1, GCA_000264685.2), *Papio papio* (GCA_028645565.1), *Paracanthobrama guichenoti* (GCA_018749465.1), *Paraescarpia echinospica* (GCA_020002185.1), *Paralichthys olivaceus* (GCA_001904815.2), *Paralithodes platypus* (GCA_013283005.1, GCA_032716605.1), *Parambassis ranga* (GCA_900634625.1, GCA_900634625.2, GCF_900634625.1), *Parapoynx stratiotata* (GCA_910589355.1), *Pararge aegeria* (GCA_905163445.1, GCF_905163445.1), *Parascotia fuliginaria* (GCA_963082885.1), *Parasteatoda lunata* (GCA_949128135.1), *Pardosa pseudoannulata* (GCA_032207245.1), *Parus major* (GCA_001522545.3, GCF_001522545.3), *Pasiphila rectangulata* (GCA_963082625.1), *Passer domesticus* (GCA_001700915.1), *Passerculus sandwichensis* (GCA_031885435.1), *Patella depressa* (GCA_948474765.1), *Patella pellucida* (GCA_917208275.1), *Patella ulyssiponensis* (GCA_963678685.1), *Patella vulgata* (GCA_932274485.2, GCF_932274485.2), *Patiria pectinifera* (GCA_029964075.1), *Pecten maximus* (GCA_902652985.1, GCF_902652985.1), *Pectinophora gossypiella* (GCF_024362695.1, GCA_024362695.1), *Pedicia rivosa* (GCA_963082725.1), *Pelecanus crispus* (GCA_030463565.1), *Pelobates cultripes* (GCA_933207985.1), *Pelochelys cantorii* (GCA_032595735.1), *Pelosia muscerda* (GCA_963691645.1), *Pemphredon lugubris* (GCA_933228935.1), *Penaeus chinensis* (GCA_019202785.2, GCA_016920825.1), *Penaeus monodon* (GCF_015228065.2, GCA_015228065.1), *Perca flavescens* (GCF_004354835.1, GCA_004354835.1), *Perca fluviatilis* (GCF_010015445.1, GCA_010015445.1), *Perca schrenkii* (GCA_025617405.1), *Perccottus glenii* (GCA_024416835.2), *Peribatodes rhomboidaria* (GCA_911728515.1), *Periophthalmus magnuspinnatus* (GCA_026225835.1, GCF_009829125.3, GCA_009829125.3), *Periparus ater* (GCA_032357785.1), *Periphyllus acericola* (GCA_949715065.1), *Perizoma affinitatum* (GCA_961405105.1), *Perizoma flavofasciatum* (GCA_958496245.1), *Perognathus longimembris pacificus* (GCA_023159225.1, GCF_023159225.1), *Peromyscus eremicus* (GCA_949786415.1, GCA_949786435.1, GCF_949786415.1), *Peromyscus leucopus* (GCF_004664715.2, GCA_004664715.2), *Peromyscus maniculatus bairdii* (GCF_003704035.1, GCA_003704035.3), *Peromyscus polionotus subgriseus* (GCA_003704135.2), *Petromyzon marinus* (GCA_010993605.1, GCF_010993605.1), *Petrophora chlorosata* (GCA_951640565.1), *Petrosia ficiformis* (GCA_947044365.1), *Phacochoerus africanus* (GCF_016906955.1, GCA_016906955.1), *Phaedon cochleariae* (GCA_918026855.4), *Phaenicophaeus curvirostris* (GCA_032191515.2), *Phalacrocorax aristotelis* (GCA_949628215.1), *Phalanger gymnotis* (GCA_028646595.1), *Phalera bucephala* (GCA_905147815.2), *Phasia obesa* (GCA_949628195.1), *Phenacoccus solenopsis* (GCA_009761765.1), *Phengaris arion* (GCA_963565745.1), *Pheosia gnoma* (GCA_905404115.1), *Pheosia tremula* (GCA_905333125.1), *Pherbina coryleti* (GCA_943735915.1), *Philereme vetulata* (GCA_918857605.2), *Philonthus cognatus* (GCA_932526585.2), *Philonthus spinipes* (GCA_963082785.1), *Phlebotomus papatasi* (GCF_024763615.1, GCA_024763615.2), *Phlogophora meticulosa* (GCA_905147745.2), *Phocoena sinus* (GCA_008692025.1, GCF_008692025.1), *Phodopus sungorus* (GCA_023856395.1), *Phoenicopterus ruber ruber* (GCA_009819775.1), *Pholidichthys leucotaenia* (GCA_020510985.1), *Pholis gunnellus* (GCA_910591455.2), *Phorcus lineatus* (GCA_921293015.1), *Phosphuga atrata* (GCA_944588485.1), *Phoxinus phoxinus* (GCA_949152265.1), *Phragmatobia fuliginosa* (GCA_932526445.1), *Phrynosoma platyrhinos* (GCA_020142125.1), *Phycodurus eques* (GCA_024500275.1), *Phyllopteryx taeniolatus* (GCA_024500385.1, GCA_019802525.1, GCA_019802545.1), *Phyllostomus discolor* (GCA_004126475.3, GCF_004126475.2), *Phyllotreta cruciferae* (GCA_917563865.1), *Phyllotreta striolata* (GCA_918026865.1), *Phymatocera aterrima* (GCA_963170745.1), *Phymorhynchus buccinoides* (GCA_017654935.2), *Physeter catodon* (GCF_002837175.3, GCA_002837175.5), *Phyto melanocephala* (GCA_941918925.1), *Pictodentalium vernedei* (GCA_031216915.1), *Picus viridis* (GCA_033816785.1), *Pieris brassicae* (GCF_905147105.1, GCA_905147105.1), *Pieris mannii* (GCA_028984075.1, GCA_029001895.1), *Pieris napi* (GCA_905231885.1, GCF_905475465.1, GCA_905475465.2), *Pieris rapae* (GCF_905147795.1, GCA_905147795.1), *Piezodorus guildinii* (GCA_023052935.1), *Piliocolobus tephrosceles* (GCF_002776525.5, GCA_002776525.5), *Pilophorus perplexus* (GCA_955831195.1), *Pinctada fucata* (GCA_028142955.1, GCA_028253585.1), *Pinctada imbricata* (GCA_002216045.1), *Pipistrellus pipistrellus* (GCA_903992545.1), *Pipistrellus pygmaeus* (GCA_949987585.1), *Pisaster ochraceus* (GCA_010994315.2), *Piscicola geometra* (GCA_943735955.1), *Pithecia pithecia* (GCA_028551515.1), *Plagiodera versicolora* (GCA_963584125.1), *Plagodis dolabraria* (GCA_963854805.1), *Planococcus citri* (GCA_950023065.1), *Platichthys flesus* (GCA_949316205.1), *Platichthys stellatus* (GCA_016801935.1), *Platycheirus albimanus* (GCA_916050605.2), *Platycnemis pennipes* (GCA_933228895.1), *Platygaster robiniae* (GCA_028476575.1), *Platypus cylindrus* (GCA_949748235.1), *Plazaster borealis* (GCA_021014325.1), *Plebejus argus* (GCA_905404155.3), *Plecotus auritus* (GCA_963455305.1), *Plectropomus leopardus* (GCF_008729295.1, GCA_026936395.1, GCA_008729295.2, GCA_011397275.1), *Plemyria rubiginata* (GCA_963576535.1), *Pleurodeles waltl* (GCA_026652325.1), *Pleuronectes platessa* (GCF_947347685.1, GCA_947347685.1), *Plodia interpunctella* (GCA_027563975.1, GCF_027563975.1), *Plotosus lineatus* (GCA_024760905.1), *Plusia festucae* (GCA_950381575.1), *Plutella xylostella* (GCA_932276165.1, GCF_932276165.1, GCA_019096205.1), *Pluvialis apricaria* (GCA_017639485.1), *Pocota personata* (GCA_963082735.1), *Podabrus alpinus* (GCA_932274525.1), *Podarcis cretensis* (GCA_951804945.1), *Podarcis lilfordi* (GCA_947686815.1), *Podarcis muralis* (GCA_004329235.1, GCF_004329235.1), *Podarcis muralis nigriventris* (GCA_014706415.1), *Podarcis raffonei* (GCA_027172205.1, GCF_027172205.1), *Podargus strigoides* (GCA_028020825.1), *Poecile atricapillus* (GCA_030490855.1, GCA_030490865.1, GCF_030490865.1), *Poecilia formosa* (GCA_013036135.2, GCA_013036085.2), *Poecilia picta* (GCA_033032245.1), *Poecilia reticulata* (GCF_000633615.1, GCA_000633615.2, GCA_904066995.1), *Poecilobothrus nobilitatus* (GCA_947095535.1), *Pogoniulus pusillus* (GCA_015220805.1), *Pogonophryne albipinna* (GCA_028583405.1), *Pogonus chalceus* (GCA_002278615.1), *Polia nebulosa* (GCA_951329385.1), *Polietes domitor* (GCA_947397865.1), *Pollachius pollachius* (GCA_949987615.1), *Pollenia amentaria* (GCA_943735925.1), *Pollenia angustigena* (GCA_930367215.1), *Pollenia labialis* (GCA_949318255.1), *Polydactylus sextarius* (GCA_016801845.1), *Polydrusus cervinus* (GCA_935413205.1), *Polydrusus tereticollis* (GCA_963576705.1), *Polymixis flavicincta* (GCA_949987655.1), *Polymixis lichenea* (GCA_949091785.1), *Polyodon spathula* (GCF_017654505.1, GCA_017654505.1), *Polyommatus icarus* (GCA_937595015.1), *Polyommatus iphigenia* (GCA_963422495.1), *Polypedilum vanderplanki* (GCA_018290095.1), *Polyploca ridens* (GCA_951394255.1), *Polypterus senegalus* (GCA_016835505.1, GCF_016835505.1), *Pomacea canaliculata* (GCA_004794335.1, GCA_003073045.1, GCF_003073045.1), *Pongo abelii* (GCF_028885655.1, GCA_028885685.1, GCA_000001545.3, GCF_002880775.1, GCA_028885655.1, GCA_002880775.3), *Pongo pygmaeus* (GCA_028885625.1, GCF_028885625.1, GCA_900086635.1, GCA_028885525.1), *Porites lutea* (GCA_958299795.1), *Porphyrio hochstetteri* (GCA_020800305.1), *Portevinia maculata* (GCA_949715645.1), *Portunus trituberculatus* (GCA_032715055.1, GCA_017591435.1), *Potorous gilbertii* (GCA_028658325.1), *Potos flavus* (GCA_028534135.1), *Prinia subflava* (GCA_021018805.1), *Prionailurus bengalensis* (GCA_016509475.2, GCF_016509475.1), *Prionailurus viverrinus* (GCF_022837055.1, GCA_022837055.1, GCA_028551425.1), *Pristionchus pacificus* (GCA_000180635.4), *Pristis pectinata* (GCF_009764475.1, GCA_009764475.2), *Procambarus clarkii* (GCA_020424385.2), *Procyon lotor* (GCA_028646535.1), *Propsilocerus akamusi* (GCA_018397935.1), *Propylea japonica* (GCA_013421045.1), *Proterorhinus semilunaris* (GCA_021464625.1), *Protocalliphora azurea* (GCA_932274085.1), *Protodeltote pygarga* (GCA_936450705.2), *Protonemura montana* (GCA_947568835.1), *Protonibea diacanthus* (GCA_028641955.1), *Protophormia terraenovae* (GCA_951394005.1), *Protopterus annectens* (GCA_019279795.1, GCF_019279795.1), *Protosalanx chinensis* (GCA_030340685.1), *Protula sp. h YS-2021* (GCA_949752745.1), *Pseudochaenichthys georgianus* (GCA_902827115.2, GCF_902827115.1, GCA_902827115.1), *Pseudocheirus occidentalis* (GCA_028646575.1), *Pseudochirops corinnae* (GCA_028646515.1), *Pseudochirops cupreus* (GCA_028627135.1), *Pseudoips prasinana* (GCA_951640165.1), *Pseudoliparis swirei* (GCF_029220125.1, GCA_029220125.1), *Pseudolycoriella hygida* (GCA_029228625.1), *Pseudophryne corroboree* (GCA_028390025.1), *Pseudorasbora parva* (GCA_024679245.1), *Psittacula echo* (GCA_963264785.1), *Psylliodes chrysocephala* (GCA_927349885.1), *Pterocles gutturalis* (GCA_009769525.1), *Pteromalus puparum* (GCA_012977825.3), *Pteropus rufus* (GCA_028533765.1), *Pterostichus madidus* (GCA_911728475.2), *Pterostichus niger* (GCA_947425015.1), *Ptilodon capucinus* (GCA_914767695.1), *Ptychobarbus kaznakovi* (GCA_027422405.1), *Ptychoptera albimana* (GCA_961205885.1), *Puma concolor* (GCA_028749985.3, GCA_028749965.3), *Pungitius pungitius* (GCA_949316345.1), *Puntigrus tetrazona* (GCA_018831695.1, GCF_018831695.1), *Pusa sibirica* (GCA_028975605.1), *Pycnopodia helianthoides* (GCA_032158295.1), *Pyemotes zhonghuajia* (GCA_025170145.1), *Pygocentrus nattereri* (GCF_015220715.1, GCA_015220715.1), *Pyralis farinalis* (GCA_947507595.1), *Pyrausta aurata* (GCA_963584085.1), *Pyrausta nigrata* (GCA_949316185.1), *Pyrgus malvae* (GCA_911387765.1), *Pyrochroa serraticornis* (GCA_905333025.2), *Pyrrhosoma nymphula* (GCA_963573305.1), *Pyxicephalus adspersus* (GCA_004786255.1, GCA_032062135.1), *Rafetus swinhoei* (GCA_019425775.1), *Raja brachyura* (GCA_963514005.1), *Rana kukunoris* (GCA_029574335.1), *Rana muscosa* (GCA_029206835.1), *Rana temporaria* (GCF_905171775.1, GCA_905171775.1), *Rangifer tarandus caribou* (GCA_019903745.2), *Rangifer tarandus platyrhynchus* (GCA_949782905.1, GCA_951394145.1), *Ranitomeya imitator* (GCA_032444005.1), *Rapana venosa* (GCA_028751875.1), *Rattus norvegicus* (GCA_000001895.4, GCA_015227675.2, GCF_015227675.2, GCF_000002265.2, GCF_000001895.5, GCA_023515785.1, GCA_000002265.1, GCA_023515805.1), *Rattus rattus* (GCA_011064425.1, GCA_011800105.1, GCF_011064425.1), *Reinhardtius hippoglossoides* (GCA_006182925.3), *Rhagium mordax* (GCA_963680705.1), *Rhagonycha fulva* (GCA_905340355.1), *Rhagonycha lutea* (GCA_958510855.1), *Rhaphigaster nebulosa* (GCA_948150685.1), *Rhea pennata* (GCA_028389875.1), *Rhegmatorhina hoffmannsi* (GCA_013398505.2), *Rhinatrema bivittatum* (GCA_901001135.1, GCF_901001135.1, GCA_901001135.2), *Rhincodon typus* (GCF_021869965.1, GCA_021869965.1), *Rhineura floridana* (GCA_030035675.1, GCF_030035675.1), *Rhingia campestris* (GCA_932526625.1), *Rhingia rostrata* (GCA_949824845.1), *Rhinoceros unicornis* (GCA_028646465.1), *Rhinolophus ferrumequinum* (GCA_004115265.3, GCF_004115265.2), *Rhinopithecus roxellana* (GCF_007565055.1, GCA_007565055.1), *Rhipicephalus microplus* (GCA_013339725.1, GCF_013339725.1), *Rhipicephalus sanguineus* (GCA_013339695.2, GCF_013339695.2), *Rhodophaea formosa* (GCA_963082605.1), *Rhogogaster chlorosoma* (GCA_944452935.1), *Rhopalosiphum maidis* (GCF_003676215.2, GCA_003676215.3), *Rhopalosiphum padi* (GCF_020882245.1, GCA_020882245.1), *Rhorus exstirpatorius* (GCA_963564615.1), *Rhynochetos jubatus* (GCA_027574665.1), *Ricordea florida* (GCA_949710005.1), *Riptortus pedestris* (GCA_019009955.1), *Rissa tridactyla* (GCA_028500815.1, GCF_028500815.1, GCA_028501385.1), *Romanogobio albipinnatus* (GCA_023566225.1), *Rousettus madagascariensis* (GCA_028533395.1), *Ruditapes philippinarum* (GCA_009026015.1), *Rutilus rutilus* (GCA_951802725.1), *Rutpela maculata* (GCA_936432065.2), *Sabethes cyaneus* (GCF_943734655.1, GCA_943734655.2), *Sacculina carcini* (GCA_916048095.2), *Saguinus midas* (GCA_021498475.1), *Saguinus oedipus* (GCA_031835075.1), *Salarias fasciatus* (GCA_902148845.1, GCF_902148845.1), *Salminus brasiliensis* (GCA_030463535.1), *Salmo salar* (GCA_000233375.4, GCA_923944775.2, GCA_905237065.2, GCA_931346935.2, GCF_000233375.1, GCF_905237065.1, GCA_021399835.1), *Salmo trutta* (GCA_901001165.1, GCA_901001165.2, GCF_901001165.1), *Salpingus planirostris* (GCA_949788335.1), *Salvelinus fontinalis* (GCF_029448725.1, GCA_029448725.1), *Salvelinus namaycush* (GCF_016432855.1, GCA_016432855.1), *Salvelinus sp. IW2-2015* (GCA_002910315.2, GCF_002910315.2), *Samia ricini* (GCA_014132275.2), *Sander lucioperca* (GCA_008315115.2, GCF_008315115.2, GCA_023718385.1), *Sander vitreus* (GCA_031162955.1), *Sarcophaga caerulescens* (GCA_927399465.1), *Sarcophaga carnaria* (GCA_958299815.1), *Sarcophaga peregrina* (GCA_014635995.1), *Sarcophaga rosellei* (GCA_930367235.1), *Sarcophaga subvicina* (GCA_936449025.2), *Sarcophaga variegata* (GCA_932273835.1), *Sarcophilus harrisii* (GCF_902635505.1, GCA_902635505.1), *Sarcoptes scabiei* (GCA_020844145.1), *Sardina pilchardus* (GCA_963854185.1), *Saturnia japonica* (GCA_033032175.1), *Saturnia pavonia* (GCA_947532125.1), *Scaeva pyrastri* (GCA_905146935.1), *Scardinius erythrophthalmus* (GCA_024453875.1), *Scatophagus argus* (GCF_020382885.2, GCA_020382885.1), *Sceloporus tristichus* (GCA_016801065.1, GCA_016801415.1), *Sceloporus undulatus* (GCA_019175285.1, GCF_019175285.1), *Schistocerca americana* (GCA_021461395.2, GCF_021461395.2), *Schistocerca cancellata* (GCA_023864275.2, GCF_023864275.1), *Schistocerca gregaria* (GCA_023897955.2, GCF_023897955.1), *Schistocerca nitens* (GCA_023898315.2, GCF_023898315.1), *Schistocerca piceifrons* (GCF_021461385.2, GCA_021461385.2), *Schistocerca serialis cubense* (GCA_023864345.3, GCF_023864345.2), *Schistosoma bovis* (GCA_944470425.2, GCA_944470445.2), *Schistosoma curassoni* (GCA_944474815.3), *Schistosoma guineensis* (GCA_944470375.2), *Schistosoma haematobium* (GCA_000699445.3, GCA_944470465.2, GCF_000699445.3, GCA_944470455.2), *Schistosoma intercalatum* (GCA_944470365.2, GCA_944470385.2), *Schistosoma japonicum* (GCA_021461655.1, GCA_025215515.1), *Schistosoma mansoni* (GCA_000237925.2, GCA_000237925.5), *Schistosoma margrebowiei* (GCA_944470205.2), *Schistosoma mattheei* (GCA_944470405.2), *Schistosoma rodhaini* (GCA_944470415.2, GCA_944470435.2), *Schistosoma spindale* (GCA_946903255.1), *Schistosoma turkestanicum* (GCA_944470395.2), *Schizaphis graminum* (GCA_020882235.1), *Schizopygopsis malacanthus* (GCA_027474115.1), *Schizopygopsis pylzovi* (GCA_027422435.1), *Schizothorax lantsangensis* (GCA_027422415.1), *Schizotus pectinicornis* (GCA_951805265.1), *Schlechtendalia chinensis* (GCA_019022885.1), *Schmidtea mediterranea* (GCA_022537955.1), *Schrankia costaestrigalis* (GCA_905475405.1), *Sciaenops ocellatus* (GCA_014183145.1, GCA_033000465.1), *Sciurus carolinensis* (GCF_902686445.1, GCA_902686445.2, GCA_028643795.1), *Sciurus vulgaris* (GCA_902686455.2), *Scleropages formosus* (GCA_023634345.2, GCF_900964775.1, GCA_023634425.1, GCA_900964775.1), *Scolanthus callimorphus* (GCA_033964015.1), *Scomber japonicus* (GCA_027409825.1, GCF_027409825.1), *Scomber scombrus* (GCA_963691925.1), *Scophthalmus maximus* (GCF_013347765.1, GCA_003186165.1, GCF_022379125.1, GCA_013347765.1, GCA_963854745.1, GCA_022379125.1), *Scortum barcoo* (GCA_023238725.1), *Scotopteryx bipunctaria* (GCA_949320045.1), *Scrobipalpa costella* (GCA_949820665.1), *Scyliorhinus canicula* (GCA_902713615.2, GCF_902713615.1, GCA_902713615.1), *Sebastes schlegelii* (GCA_014673565.1), *Sebastes umbrosus* (GCF_015220745.1, GCA_015220745.1), *Seladonia tumulorum* (GCA_913789895.3), *Selenia dentaria* (GCA_917880725.1), *Sepiola atlantica* (GCA_963556195.1), *Sepioteuthis lessoniana* (GCA_963585895.1), *Serinus canaria* (GCA_022539315.2, GCF_022539315.1), *Seriola aureovittata* (GCA_021018895.1, GCF_021018895.1, GCA_023856345.1), *Sesia apiformis* (GCA_914767545.1), *Sesia bembeciformis* (GCA_943735995.1), *Setipinna tenuifilis* (GCA_030347295.1), *Setophaga coronata coronata* (GCA_001746935.2), *Shargacucullia verbasci* (GCA_947562105.1), *Shinisaurus crocodilurus* (GCA_021292165.1), *Sialis fuliginosa* (GCA_961205875.1), *Sialis lutaria* (GCA_949319165.1), *Sicus ferrugineus* (GCA_922984085.1), *Silurus aristotelis* (GCA_946808225.1), *Silurus meridionalis* (GCF_014805685.1, GCA_023972475.1, GCA_014805685.1), *Sinella curviseta* (GCA_004115045.4), *Siniperca chuatsi* (GCA_011952085.1, GCF_020085105.1, GCA_027580155.1, GCA_020085105.1), *Siniperca knerii* (GCA_011952075.1), *Siniperca scherzeri* (GCA_011952095.1, GCA_027580175.1), *Sinohyriopsis cumingii* (GCA_028554795.1), *Sinonovacula constricta* (GCA_009762815.1, GCA_007844125.1), *Sinotaia purificata* (GCA_028829895.1), *Siphamia tubifer* (GCA_020466265.1), *Siphlonurus alternatus* (GCA_949825025.1), *Siphonodentalium dalli* (GCA_032622095.1), *Sipunculus nudus* (GCA_026874595.1), *Sisyra nigra* (GCA_958496155.1), *Sisyra terminalis* (GCA_958496175.1), *Sitobion miscanthi* (GCA_008086715.1), *Sitodiplosis mosellana* (GCA_021018905.1), *Sitta carolinensis* (GCA_032173965.1), *Slavum lentiscoides* (GCA_032441835.1), *Smittia aterrima* (GCA_033063855.1), *Smittia sp. YW1114-2* (GCA_033064975.1), *Sogatella furcifera* (GCA_017141385.1, GCA_014356515.1), *Solea senegalensis* (GCA_919967415.2, GCA_019176455.1, GCF_019176455.1), *Solea solea* (GCF_958295425.1, GCA_958295425.1), *Solen grandis* (GCA_021229015.1), *Solenopsis invicta* (GCA_018691235.1, GCF_016802725.1, GCA_009650705.1, GCA_009299965.1, GCA_009299975.1, GCA_010367695.1, GCA_016802725.1), *Somateria mollissima* (GCA_030142145.1, GCA_951416345.1, GCA_951411735.1), *Sorex araneus* (GCA_027595985.1, GCF_027595985.1), *Sparus aurata* (GCA_003309015.1, GCA_900880675.1, GCF_900880675.1, GCA_900880675.2), *Spea bombifrons* (GCA_027358695.2, GCF_027358695.1), *Sphaeramia orbicularis* (GCF_902148855.1, GCA_902148855.1), *Sphaerodactylus townsendi* (GCA_021028975.2, GCF_021028975.2), *Sphaerophoria taeniata* (GCA_943590905.1), *Sphecodes monilicornis* (GCA_913789915.3), *Sphenella marginata* (GCA_951509765.1), *Spheniscus humboldti* (GCA_027474245.1), *Sphinx pinastri* (GCA_947568825.1), *Spilarctia lutea* (GCA_916048165.1), *Spilosoma lubricipeda* (GCA_905220595.1), *Spisula solida* (GCA_947247005.1), *Spisula subtruncata* (GCA_963678985.1), *Spodoptera exigua* (GCA_011316535.1, GCA_022315195.1, GCA_902829305.4), *Spodoptera frugiperda* (GCA_012979215.2, GCA_011064685.2, GCF_023101765.2, GCA_023101765.3, GCF_011064685.2), *Spodoptera littoralis* (GCA_902850265.1), *Spodoptera litura* (GCF_002706865.2, GCA_002706865.3), *Spongilla lacustris* (GCA_949361645.1), *Sprattus sprattus* (GCA_963457725.1), *Squalius cephalus* (GCA_022829025.1, GCA_949319135.1), *Squalus acanthias* (GCA_030390025.1), *Stegostoma tigrinum* (GCA_022316705.1, GCA_030684315.1, GCF_022316705.1, GCF_030684315.1), *Steinernema carpocapsae* (GCA_000757645.3), *Steinernema hermaphroditum* (GCA_030435675.1), *Stelis phaeoptera* (GCA_943735885.1), *Stenchaetothrips biformis* (GCA_030522485.1), *Stenella coeruleoalba* (GCA_951394435.1), *Steno bredanensis* (GCA_028646385.1), *Stenoptilia bipunctidactyla* (GCA_944452665.1), *Stenurella melanura* (GCA_963583905.1), *Sterna hirundo* (GCA_009819605.1), *Steromphala cineraria* (GCA_916613615.1), *Sthenelais limicola* (GCA_942159475.1), *Stichopus chloronotus* (GCA_021234535.1), *Stictonetta naevosa* (GCA_011074415.1), *Stomorhina lunata* (GCA_933228675.1), *Stomoxys calcitrans* (GCA_963082655.1, GCF_963082655.1), *Stramonita haemastoma* (GCA_030674155.1), *Stratiomys singularior* (GCA_954870665.1), *Streblospio benedicti* (GCA_019095985.1), *Streptopelia turtur* (GCA_901699155.2), *Strigamia acuminata* (GCA_949358305.1), *Strigops habroptila* (GCA_004027225.2, GCF_004027225.2), *Strix aluco* (GCA_031877795.1, GCA_031877785.1), *Strongyloides ratti* (GCF_001040885.1, GCA_001040885.1), *Strongyloides stercoralis* (GCA_029582065.1), *Sturmia bella* (GCA_963662145.1), *Sturnus vulgaris* (GCA_023376015.1), *Subacronicta megacephala* (GCA_958496365.1), *Suillia variegata* (GCA_949127995.1), *Suncus etruscus* (GCA_024139225.1, GCF_024139225.1), *Suricata suricatta* (GCA_006229205.1, GCF_006229205.1), *Sus scrofa* (GCA_015776825.1, GCA_900119615.2, GCA_000003025.6, GCA_023065335.1, GCA_007644095.1, GCA_030704935.1, GCF_000003025.6, GCA_023065355.1, GCA_002844635.1, GCA_031225015.1, GCA_031306245.1, GCA_024718415.1), *Sus scrofa domesticus* (GCA_020567905.1, GCA_017957985.1), *Sus scrofa scrofa* (GCA_006511355.2), *Sylvia atricapilla* (GCA_009819655.1), *Sylvia borin* (GCA_014839755.1), *Sylvicola cinctus* (GCA_963854165.1), *Sylvilagus bachmani* (GCA_015711505.1), *Sylvilagus floridanus* (GCA_949820135.1), *Sympetrum striolatum* (GCA_947579665.1), *Symphalangus syndactylus* (GCA_028878055.1, GCA_028878085.1, GCA_028642525.1, GCF_028878055.1), *Symphodus melops* (GCA_947650265.1), *Symsagittifera roscoffensis* (GCA_963678635.1), *Synanceia verrucosa* (GCA_029721515.1), *Synanthedon andrenaeformis* (GCA_936446665.2), *Synanthedon formicaeformis* (GCA_945859745.1), *Synanthedon myopaeformis* (GCA_944738685.1), *Synanthedon tipuliformis* (GCA_947623395.1), *Synanthedon vespiformis* (GCA_918317495.1), *Synaphobranchus kaupii* (GCA_029718625.1), *Synchiropus splendidus* (GCA_027744825.1, GCF_027744825.2), *Syngnathoides biaculeatus* (GCA_019802595.1), *Syngnathus acus* (GCA_948146105.1, GCF_901709675.1, GCA_901709675.2), *Syngnathus scovelli* (GCA_024217435.4), *Syngnathus typhle* (GCA_033458585.1, GCF_033458585.1), *Syritta pipiens* (GCA_905187475.1), *Syrphus vitripennis* (GCA_958431115.1), *Tachina fera* (GCA_905220375.1), *Tachina grossa* (GCA_949987645.1), *Tachina lurida* (GCA_944452675.1), *Tachyglossus aculeatus* (GCA_015852505.1, GCA_015598185.1, GCF_015852505.1), *Tachypleus gigas* (GCA_014155125.1), *Tachypleus tridentatus* (GCA_004210375.1), *Tachystola acroxantha* (GCA_963506565.1), *Tachysurus fulvidraco* (GCA_022655615.1, GCF_022655615.1, GCA_023638525.1), *Tachysurus vachellii* (GCF_030014155.1, GCA_030014155.1, GCA_033026395.1), *Tadarida brasiliensis* (GCA_030848825.1), *Taenia multiceps* (GCA_001923025.3), *Taenia pisiformis* (GCA_023968675.1), *Taenioides sp. WSHXM2023* (GCA_029959045.1), *Taeniopygia guttata* (GCA_008822105.2, GCA_008822125.1, GCA_008822115.3, GCA_000151805.2, GCA_003957565.4, GCF_008822105.2, GCF_003957565.2, GCA_009859065.2, GCF_000151805.1), *Takifugu bimaculatus* (GCA_004026145.2), *Takifugu flavidus* (GCA_003711565.2, GCF_003711565.1), *Takifugu rubripes* (GCF_901000725.2, GCA_000180615.2, GCA_901000725.3, GCF_000180615.1, GCA_901000725.2), *Tamandua tetradactyla* (GCA_023851605.1), *Tamias sibiricus* (GCA_025594165.1), *Tanypteryx hageni* (GCA_028673005.1), *Taphrorychus bicolor* (GCA_951812265.1), *Tapirus indicus* (GCA_031878705.1, GCA_031878655.1), *Tapirus terrestris* (GCA_028533255.1), *Tauraco erythrolophus* (GCA_009769465.1), *Taurulus bubalis* (GCA_910589615.1), *Tautogolabrus adspersus* (GCA_020745685.1), *Tegillarca granosa* (GCA_029721355.1), *Teleiodes luculella* (GCA_948473455.1), *Telenomus remus* (GCA_020615435.1), *Teleopsis dalmanni* (GCA_002237135.2, GCA_002237135.5, GCF_002237135.1), *Telmatherina bonti* (GCA_933228915.1), *Tenodera sinensis* (GCA_030324105.1, GCA_030765045.1), *Tenthredo distinguenda* (GCA_947538915.1), *Tenthredo mesomela* (GCA_943736025.1), *Tenthredo notha* (GCA_914767705.2), *Terebella lapidaria* (GCA_949152475.1), *Tetanocera ferruginea* (GCA_958299015.1), *Tethea ocularis* (GCA_963555595.1), *Tetheella fluctuosa* (GCA_951216915.1), *Tetragnatha montana* (GCA_963680715.1), *Tetramorium bicarinatum* (GCA_928718305.1), *Tetrao urogallus* (GCA_951394365.1), *Thalassophryne amazonica* (GCA_902500255.1, GCF_902500255.1), *Thaleichthys pacificus* (GCA_023658055.1), *Thalpophila matura* (GCA_948465475.1), *Thamnaconus septentrionalis* (GCA_009823395.1), *Thamnophis elegans* (GCA_009769535.1, GCF_009769535.1), *Thecocarcelia acutangulata* (GCA_914767995.1), *Thecophora atra* (GCA_937620795.1), *Thelaira solivaga* (GCA_947397855.1), *Theocolax elegans* (GCA_026168455.1), *Thera britannica* (GCA_939531255.2), *Thera obeliscata* (GCA_947578465.1), *Theretra japonica* (GCA_033459515.1), *Thereva nobilitata* (GCA_963855945.1), *Thereva unica* (GCA_949987705.1), *Therioaphis trifolii* (GCA_027580255.1), *Theristicus caerulescens* (GCA_020745775.1), *Theropithecus gelada* (GCA_028533075.1, GCF_003255815.1, GCA_003255815.1), *Tholera decimalis* (GCA_943138885.2), *Thomomys bottae* (GCA_031878675.1, GCA_031878665.1), *Thumatha senex* (GCA_948477245.1), *Thunnus albacares* (GCA_914725855.1, GCA_914725855.2, GCF_914725855.1), *Thunnus maccoyii* (GCA_910596095.1, GCF_910596095.1), *Thyatira batis* (GCA_905147785.2), *Thylacinus cynocephalus* (GCA_007646695.3), *Thymallus thymallus* (GCA_023634145.1, GCA_004348285.1), *Thymelicus acteon* (GCA_951805285.1), *Thymelicus sylvestris* (GCA_911387775.1), *Thysanoteuthis rhombus* (GCA_963457665.1), *Tiaroga cobitis* (GCA_030578255.1), *Tigriopus californicus* (GCA_007210705.1, GCF_007210705.1), *Tigriopus japonicus* (GCA_010645155.1), *Tiliacea aurago* (GCA_948098905.1), *Timandra comae* (GCA_958496195.1), *Timema cristinae* (GCA_002009905.3, GCA_002928295.1), *Tinea pellionella* (GCA_948150575.1), *Tinea semifulvella* (GCA_910589645.1), *Tinea trinotella* (GCA_905220615.1), *Tiphia femorata* (GCA_944319695.1), *Tipula confusa* (GCA_963556175.1), *Tipula helvola* (GCA_963556165.1), *Tipula unca* (GCA_951394425.1), *Tipula vernalis* (GCA_958295665.1), *Tolmerus cingulatus* (GCA_959613345.1), *Tomocerus qinae* (GCA_020055645.1), *Topomyia yanbarensis* (GCF_030247195.1, GCA_030247195.1), *Tortricodes alternella* (GCA_947859335.1), *Tortrix viridana* (GCA_963241965.1), *Toxonevra muliebris* (GCA_963691655.1), *Toxorhynchites rutilus septentrionalis* (GCA_029784135.1, GCF_029784135.1), *Toxotes chatareus* (GCA_016801885.1), *Toxotes jaculatrix* (GCF_017976425.1, GCA_017976425.1), *Trachemys scripta elegans* (GCA_013100865.1, GCF_013100865.1), *Trachops cirrhosus* (GCA_028533065.1), *Trachurus trachurus* (GCA_905171665.2), *Tremarctos ornatus* (GCA_028551375.1), *Tribolium castaneum* (GCA_031307605.1, GCA_000002335.3, GCF_000002335.3), *Tribolium confusum* (GCA_019155225.1), *Tribolium freemani* (GCA_939628115.1, GCA_022388455.1), *Trichinella murrelli* (GCA_002221485.1), *Trichinella spiralis* (GCA_008807755.1, GCA_008807815.1, GCA_000181795.3, GCA_008807795.1, GCA_008807775.1), *Trichobilharzia regenti* (GCA_944472135.2), *Trichobilharzia szidati* (GCA_944472155.2), *Tricholauxania praeusta* (GCA_949775025.1), *Trichomycterus rosablanca* (GCA_030014385.1), *Trichoplusia ni* (GCA_003590095.1, GCF_003590095.1), *Trichopria drosophilae* (GCA_030407085.1), *Trichosurus vulpecula* (GCF_011100635.1, GCA_011100635.1), *Tridacna crocea* (GCA_032873355.1, GCA_943736015.1), *Tridacna derasa* (GCA_963210305.1), *Tridacna gigas* (GCA_945859785.2), *Trididemnum clinides* (GCA_963675345.1), *Trilocha varians* (GCA_030269945.2), *Triplophysa bombifrons* (GCA_029783895.1), *Triplophysa dalaica* (GCA_015846415.1, GCF_015846415.1), *Triplophysa rosa* (GCF_024868665.1, GCA_024868665.2), *Triplophysa tibetana* (GCA_008369825.1), *Triplophysa yarkandensis* (GCA_033220385.1), *Trisateles emortualis* (GCA_947095525.1), *Trogon surrucura* (GCA_020746105.1), *Troides aeacus* (GCA_033220335.2), *Tromatobia lineatoria* (GCA_949699805.1), *Trypoxylus dichotomus* (GCA_023509865.1), *Tuberolachnus salignus* (GCA_956483605.1), *Tupaia chinensis* (GCA_033439345.1), *Tursiops truncatus* (GCF_011762595.1, GCA_011762595.1), *Tuta absoluta* (GCA_029230345.1), *Tyria jacobaeae* (GCA_947561695.1), *Udea ferrugalis* (GCA_950022985.1), *Udea olivalis* (GCA_947369235.1), *Uloborus diversus* (GCA_026930045.1, GCF_026930045.1), *Uranotaenia lowii* (GCF_029784155.1, GCA_029784155.1), *Urbanus simplicius* (GCA_949699795.1), *Urocyon cinereoargenteus* (GCA_032313775.1), *Uromys caudimaculatus* (GCA_028551405.1), *Urophora cardui* (GCA_960531455.1), *Ursus arctos* (GCA_023065955.2, GCF_023065955.2), *Valencia hispanica* (GCA_963556495.2), *Vandiemenella viatica* (GCA_031002005.1), *Vanessa atalanta* (GCF_905147765.1, GCA_905147765.2), *Vanessa cardui* (GCA_905220365.1, GCA_905220365.2, GCF_905220365.1), *Varecia variegata* (GCA_028533085.1), *Venturia canescens* (GCF_019457755.1, GCA_019457755.1), *Venusia cambrica* (GCA_951394065.1), *Verasper variegatus* (GCA_013332515.1, GCA_012393435.1, GCA_026259375.1), *Vespa crabro* (GCF_910589235.1, GCA_910589235.2), *Vespa velutina* (GCF_912470025.1, GCA_912470025.1), *Vespula germanica* (GCA_905340365.1, GCA_014466195.1), *Vespula pensylvanica* (GCA_014466175.1, GCF_014466175.1), *Vespula vulgaris* (GCA_905475345.1, GCA_014466185.1, GCF_905475345.1), *Vidua chalybeata* (GCF_026979565.1, GCA_026979565.1), *Vidua macroura* (GCF_024509145.1, GCA_024509145.1), *Villa cingulata* (GCA_951394055.1), *Vimba vimba* (GCA_022828995.1), *Vipera latastei* (GCA_024294585.1), *Vipera ursinii* (GCA_947247035.1), *Volucella bombylans* (GCA_949129095.1), *Volucella inanis* (GCA_907269105.1), *Volucella inflata* (GCA_928272305.1), *Vombatus ursinus* (GCA_028626985.1), *Vulpes corsac* (GCA_030463245.1), *Vulpes ferrilata* (GCA_024500485.1), *Vulpes lagopus* (GCA_018345385.1, GCF_018345385.1), *Watersipora subatra* (GCA_963576615.1), *Watsonalla binaria* (GCA_929442735.1), *Wyeomyia smithii* (GCA_029784165.1, GCF_029784165.1), *Xanthia icteritia* (GCA_949128155.2), *Xanthia togata* (GCA_963853775.1), *Xanthogramma pedissequum* (GCA_910595825.1), *Xanthorhoe designata* (GCA_963582015.1), *Xanthorhoe spadicearia* (GCA_947086425.1), *Xanthostigma xanthostigma* (GCA_963575645.1), *Xenentodon cancila* (GCA_014839995.1), *Xenopus borealis* (GCA_024363595.1), *Xenopus laevis* (GCF_017654675.1, GCA_017654675.1, GCF_001663975.1, GCA_001663975.1), *Xenopus tropicalis* (GCA_013368275.1, GCF_000004195.4, GCA_000004195.4), *Xerus rutilus* (GCA_028644305.1), *Xestia ashworthii* (GCA_950022955.1), *Xestia c-nigrum* (GCA_916618015.1), *Xestia rhomboidea* (GCA_963853795.1), *Xestia sexstrigata* (GCA_941918905.2), *Xestia xanthographa* (GCA_905147715.2), *Xestospongia muta* (GCA_963693285.1), *Xiphias gladius* (GCF_016859285.1, GCA_016859285.1), *Xiphophorus couchianus* (GCF_001444195.1, GCA_001444195.3), *Xiphophorus hellerii* (GCA_003331165.2, GCF_003331165.1), *Xiphophorus maculatus* (GCA_002775205.2, GCF_002775205.1), *Xiphydria camelus* (GCA_963678675.1), *Xylocampa areola* (GCA_935421205.1), *Xylophagus ater* (GCA_963422695.1), *Xylota segnis* (GCA_963583995.1), *Xylota sylvarum* (GCA_905220385.1), *Xyrauchen texanus* (GCA_025860055.1, GCF_025860055.1), *Xyrichtys novacula* (GCA_962446985.1), *Yoshiicerus persimilis* (GCA_020352615.2), *Yponomeuta cagnagella* (GCA_947311075.1, GCA_947310995.1), *Yponomeuta malinellus* (GCA_947308005.1), *Yponomeuta plumbellus* (GCA_947310845.1), *Yponomeuta sedellus* (GCA_934045075.1), *Ypsolopha scabrella* (GCA_910592155.1), *Ypsolopha sequella* (GCA_934047225.1), *Zalophus californianus* (GCA_009762305.2, GCF_009762305.2), *Zele albiditarsus* (GCA_958496275.1), *Zelleria hepariella* (GCA_949319315.1), *Zerene cesonia* (GCA_012273895.2, GCF_012273895.1), *Zeugodacus cucurbitae* (GCF_028554725.1, GCA_028554725.2), *Zeugodacus tau* (GCA_031772095.1), *Zeus faber* (GCA_960531495.1), *Zeuzera pyrina* (GCA_907165235.1), *Zingel zingel* (GCA_023566125.1), *Zonotrichia leucophrys* (GCA_028769735.1), *Zootoca vivipara* (GCF_963506605.1, GCF_011800845.1, GCA_011800845.1, GCA_963506605.1), *Zophobas atratus* (GCA_022388445.1), *Zoroaster sp. YZ-2022* (GCA_029582265.1), *Zygaena filipendulae* (GCA_907165275.2), *allo-octoploid hybrid of Carassius auratus x Cyprinus carpio* (GCA_024542945.1), and *unclassified Cladorhizidae* (GCA_028752895.1, GCA_028752905.1). Genomes downloaded from other sources were: *Amphioctopus fangsiao* (https://figshare.com/s/fa09f5dadcd966f020f3), *Ephydatia muelleri*(https://spaces.facsci.ualberta.ca/ephybase/), *Rhopilema esculentum* (http://gigadb.org/dataset/view/id/100720/Sample_page/2/File_page/2).

## Supplementary Information Guide

### Supplementary Information *1*

#### Glossary

In an effort to clarify the intended meanings of our text, we include below a glossary of terms defined in this manuscript, or existing terms and the meaning of these terms based on previous publications.

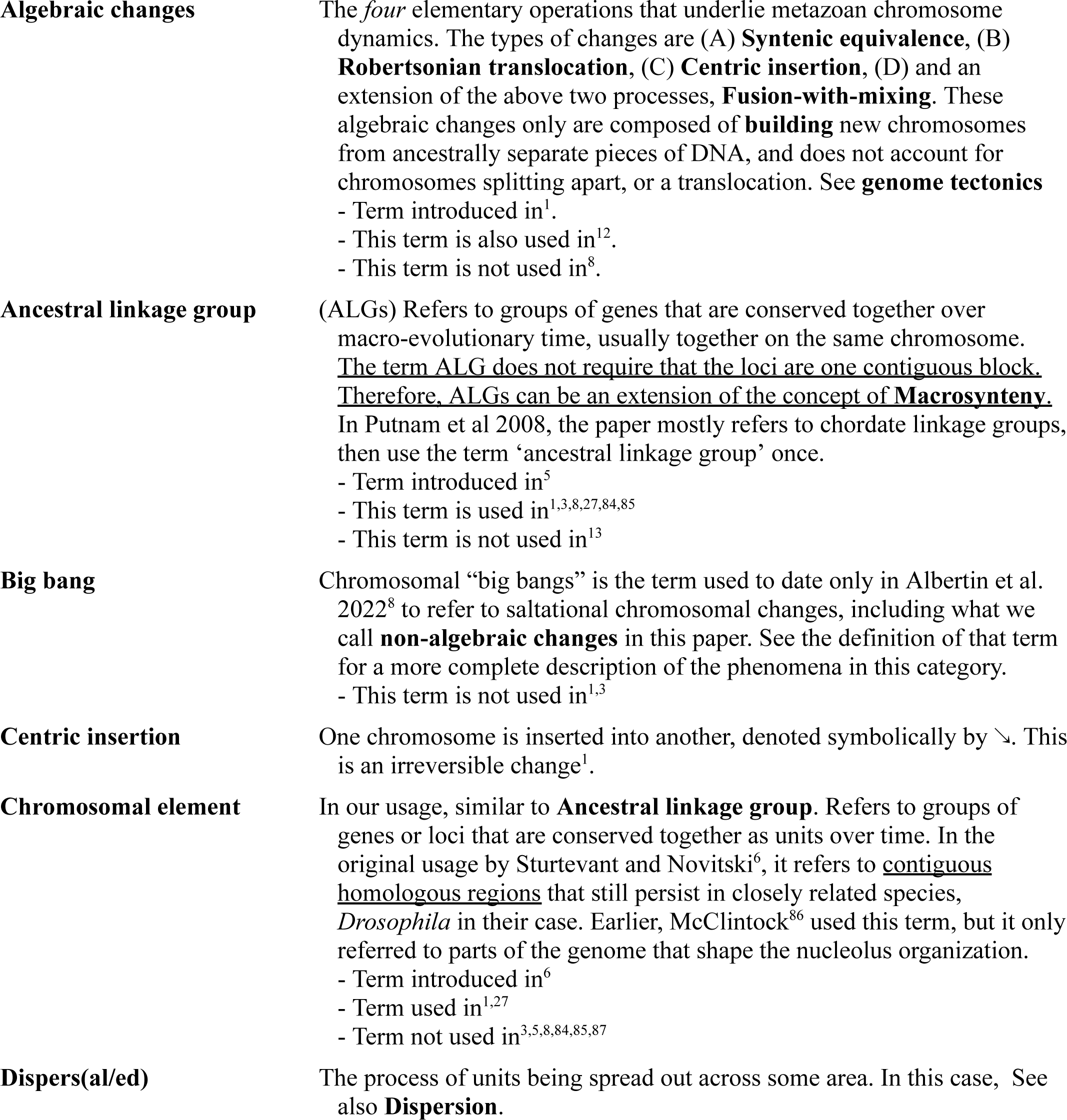
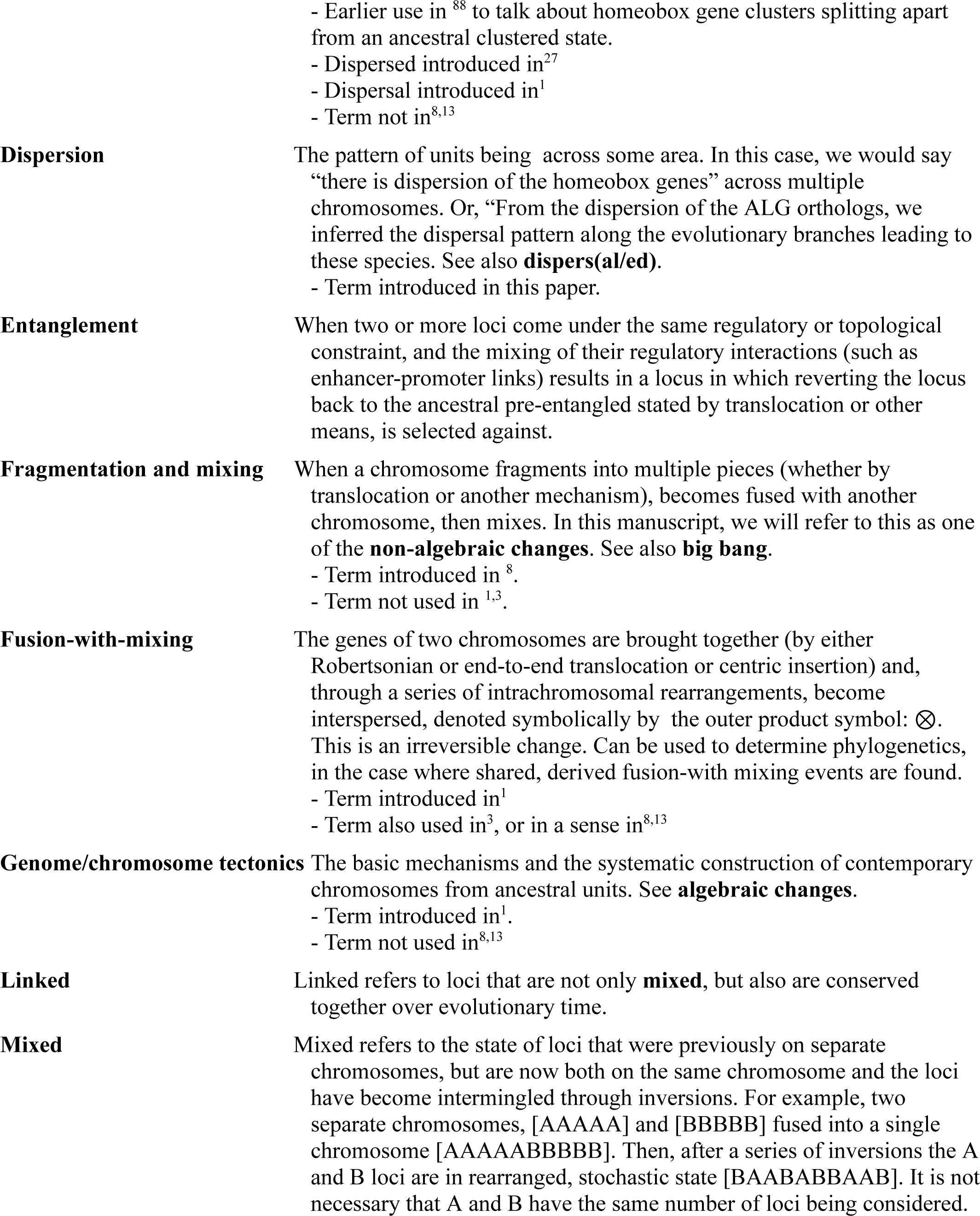
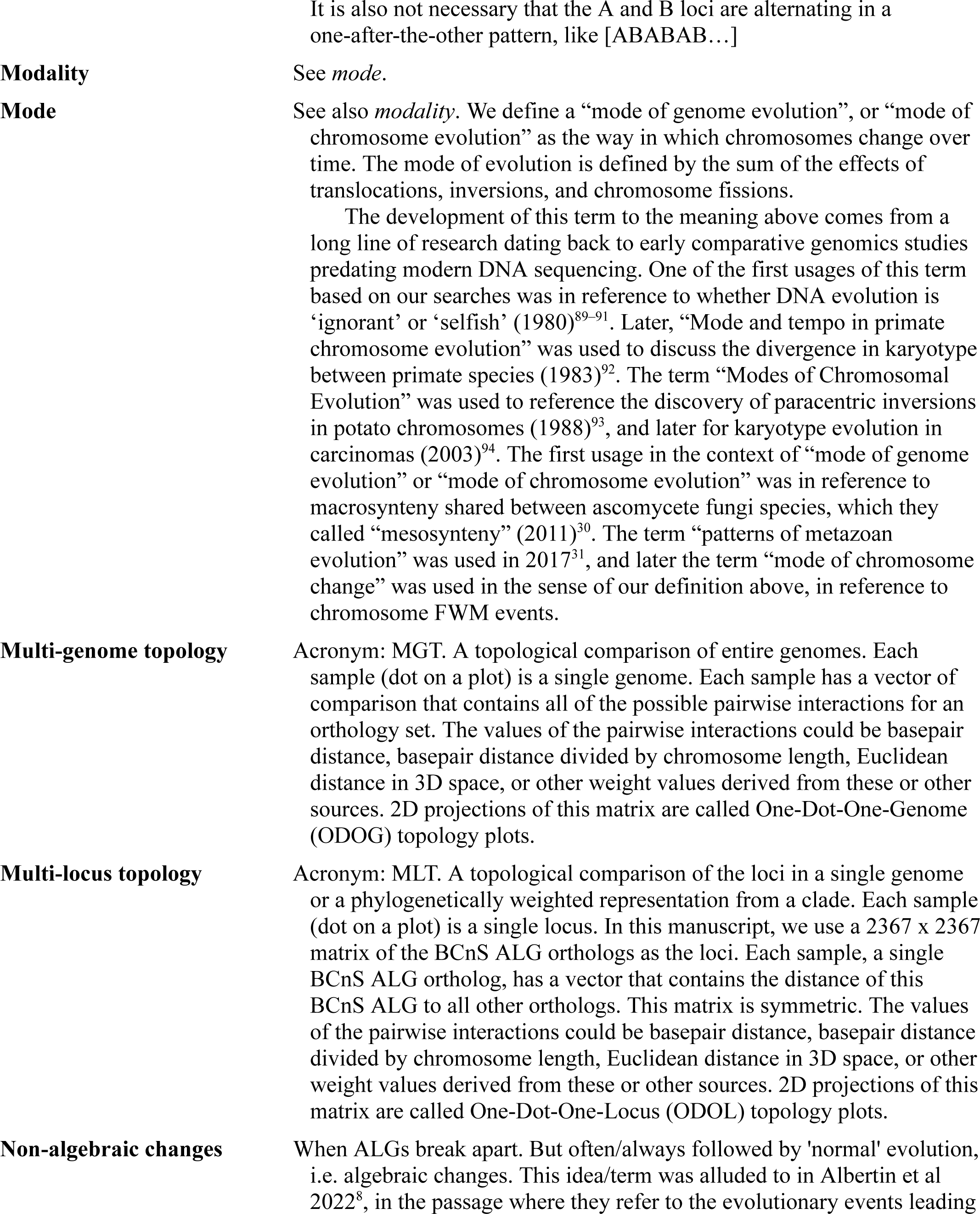
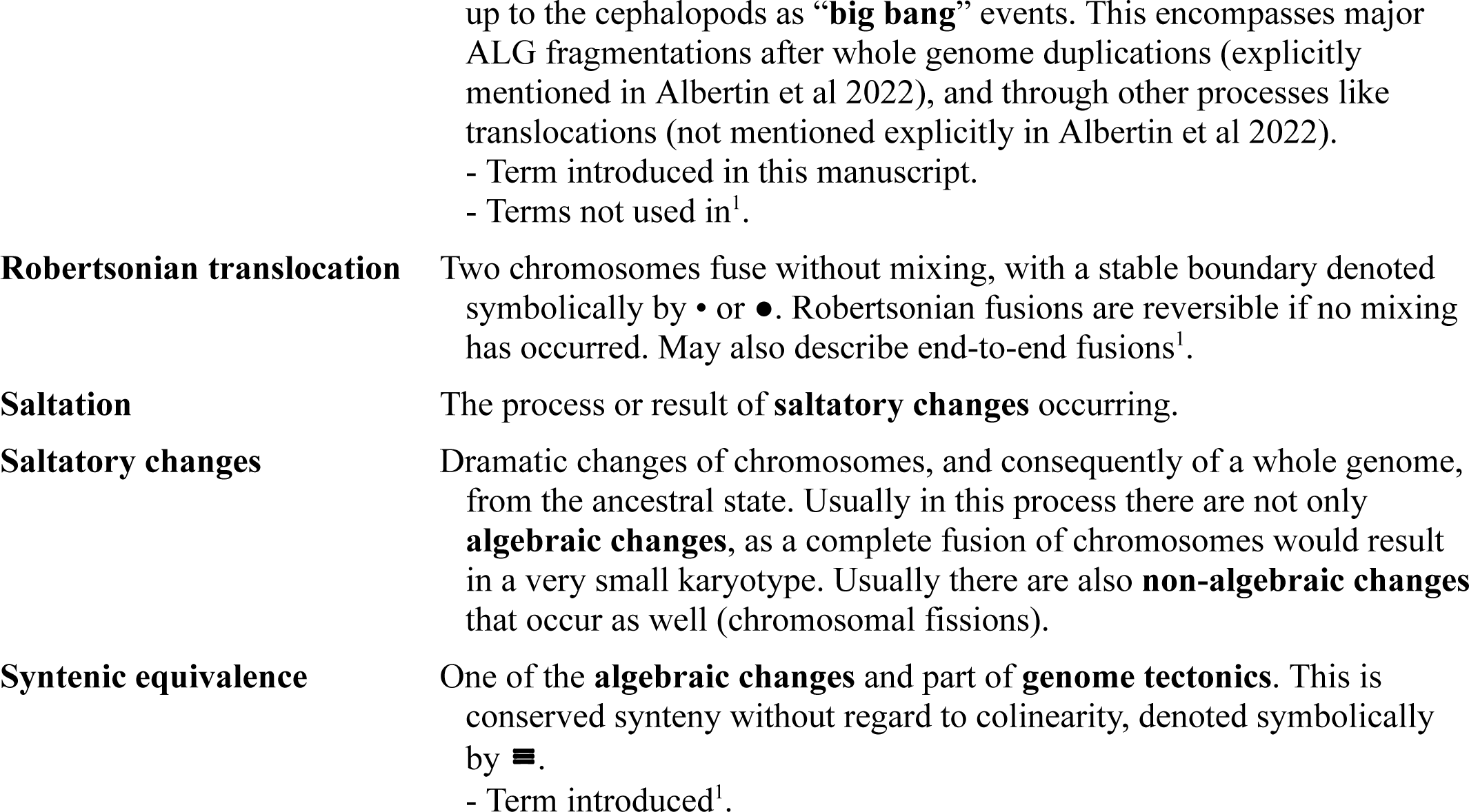

### Supplementary Information *2*

#### Periodicity in genomic changes

One persistent theme in paleobiology is the search for factors associated with, and the patterns of, speciation and extinction. One major pattern evident from the fossil record is that speciation and extinction events happen in bursts^25,95^ in spite of an apparent constant probability of extinction^96^. Moreover, there appears to be a periodicity of these bursts^25,26^, most significantly in cycles of 62^25,26^ and 140 million years^26^. The potential biologic and abiotic factors related to this periodicity are discussed in the literature cited above.

## Methods

With our dataset of chromosomal evolutionary rates across the tree of life, we sought to determine whether the changes in rates had significant underlying periods. To achieve this, we performed Fourier analyses and Monte Carlo simulations to estimate false discovery rates on the rates of chromosomal changes in vertebrates and protostomes, following the method described by Rohde and Muller (2005)^26^. In this manuscript, the authors analyzed the periodicity of the species origination and extinction rates through geological time by analyzing Sapkoski’s compendium of fossil genera^97^.

Our input data were the rates of ALG fusions or ALG losses across the vertebrates or protostomes, generated earlier in the manuscript. To process the input data, we first limited the oldest time for each clade to between 500 and 525 million years ago, given that there are very few branches at the base of each clade, and therefore a higher chance to over- or under-estimate the rates of change. We also removed the bin representing the most recent million years, as this bin often contained outliers of evolutionary rates compared to the rest of the dataset. From the truncated datasets, we then fit cubic or quartic polynomials to the rates of change, and created a detrended dataset by subtracting this function from the data. We then performed discrete Fourier transforms on the raw data, and on zero-padded data. The results measuring periods taken from both padded and non-padded data are reported in this manuscript. We identified the strongest two or three peaks in each spectral power distribution, and selected these frequencies to analyze further with Monte Carlo false discovery rate simulations.

In Rohde and Muller (2005)^26^, two types of Monte Carlo simulations were performed to infer the false discovery rates (α) for the strengths of magnitude for the 62 and 140 million year peaks in the Fourier power spectrum. The first type, ‘R’, was a random walk through the observed transitions between timepoints in the species extinction or origination measurements. The second type, ‘W’, was a random walk through clusters of transitions, wherein the original dataset of transitions was partitioned into 20 equally-sized vectors, then shuffled. In both of these methods the starting rate was kept the same as the observed data, and therefore the rate of the final bin was also the same. The benefit of the ‘W’ type is that short-term correlations are preserved, whereas they are completely lost in the ‘R’ type. We propose that the ‘R’ type simulation is an extension of the ‘W’ type, in which the block size is the same size as the input vector of observations.

We performed both types, ‘R’ and ‘W’, of Monte Carlo simulations on the chromosome change rate data. From these simulations we noticed that contrary to the reported observation that changing the number of blocks for type ‘W’ having little effect on the background power spectra, that changing the block size for the chromosome change rate data caused variations of 20-30% in the percent of spectra with magnitudes matching those in the real data. Given that “the choice of 20 blocks is somewhat arbitrary”^26^ and the high variance of the background rate, we opted to perform the ‘W’ type simulations for a rank of block sizes ranging from 10 to 50 blocks. Each simulation for each peak contained 5000 random trials, and the results were summarized reporting the percent of trials whose target peaks were above a given magnitude (See Supplemental Figure S1 in Rohde and Muller (2005)^26^). For each simulation, we also calculated the function for the assumption of exponential form.

## Results

As we mentioned, Monte Carlo simulations were performed for block quantities between [10-50) for each condition of clade, chromosomal change type, padding treatment, and period. For this reason there is a range of support values for each simulation — one for each block quantity. We will report the median support value in text.

The two conditions with the highest support were BCnS ALG losses in protosomes with a period of 102.8-107.53 million years (support of 97.96% for unpadded, 98.04% for zero-padded), BCnS ALG losses in vertebrates with a period of 49.8 million years (support of 94.01% for unpadded, 93.45% for zero-padded).

Notably, we find that for our dataset of chromosome-scale animal genomes the most highly supported condition for Metazoans is fusions with a period of 62-64.38 million years (support of 93.8% for unpadded, 93.00% for zero-padded). This is the same period that was first noticed from a fossil diversity dataset^25^, and was later found to be the strongest period underlying the species extinction and origination patterns of more than 30 thousand metazoan marine genera^26^.

## Discussion

Our findings echo previous observations of periodicity in biological evolution from the fossil record, albeit our findings are derived independently from genomic changes. This suggests that similar underlying mechanisms may drive periodicity in both fossil records and genomic evolution. The consistency of the 62-million-year cycle across different data types and methods emphasizes its potential significance in understanding the temporal dynamics of evolution. We note, and others have noted^26^, that the number of 62 million years is a consequence of the sampling rate and number of observations. Future work in this area might focus on a refinement of node ages across the tree of life, and finer modeling of the periodicity given the uncertainties of node ages, to better ascertain the variances in the periodicity of genomic changes over time.

### Supplementary Information *3*

#### State formulation of the (sub)chromosomal mixing process

##### Probability vector and matrix multiplication of genomic states

This study suggests the presence of distinct irreversible evolutionary trajectories in animal genomes, with different degrees of irreversibility, depending on the number of components that are being mixed and the strength of their coupling (for example, chromosomal meiotic constraints or regulatory linkages at the sub-chromosomal level). The irreversibility is a product of mixing of genes that are located on distinct evolutionary units. In our previous studies^1,27^, these units were simply the chromosomal elements (or chromosomes) themselves. In this study, we generalize this mixing to include mixing of previously separated sub-chromosomal elements.

This degree of irreversibility depends on the size of the mixing units (chromosomal vs sub-chromosomal) and mechanistic reasons (regulatory linkage, meiotic constraints) behind the fusion process. An important property of the mixing is the loss of the information that the constituent elements can be separated back into the more ancestral and distinct units. For this reason, we can only identify the existence of these ancestral states through evolutionary outgroup comparisons. To begin to study these processes within a more rigorous and quantitative framework, we describe a mathematical approach to quantify the degree of irreversibility and its application in the manifold approximation context presented in this paper.

##### (Sub-)chromosomal fusion-with-mixing representation

We treat each chromosome as a coordinate system assigning a probability of observation for a given genomic feature. For example, the probability of finding a gene within a chromosome can be defined as a probability density vector of a size of total chromosomal positions. To find two genes to be colocalized at the same coordinate is simply the inner product of the two corresponding gene vectors (**a * b, or < a | b >**). Genomic features move according to their propagation mechanism, such as translocations. An ‘evolution operator’ U multiplied by a given state vector (**U | a >**) gives the new probability of observing feature **a** at the next evolutionary step. The system of two elements a and b can be described as a tensor product of these two, **a** ⊗ **b**, as that (two-dimensional) matrix will peak at a point where both densities are highest in relation to the coordinate systems of a and b. For systems of many genes, this tensor product is simply generalized in a longer string of products (**a**⊗**b**⊗**c**⊗**d** …).

In our previous studies, we represented the fusion-with-mixing process as a tensor product of two chromosomes. However, this is not a completely accurate representation, as the tensor (outer) product simply defines a combination of two coordinate systems and correspondences of each element between the two coordinate systems. If a gene ‘**a**’ on chromosome **A** has probability density of **a=(1,0,0)** and gene ‘**b**’ on chromosome **B** has probability of **b=(0,0,1,0)**, then **a** ⊗ **b = (0,0,1,0; 0,0,0,0; 0;0;0;0)** (or an expanded vector). This describes the system state according to the corresponding coordinate systems (e.g., chromosomal position) of **a** and **b**, but does not provide a framework to qualitatively or quantitatively describe how and how well the chromosomes **a** and **b** have become mixed since their initial fusion event

To represent the mixing property one has to measure the ‘entanglement’ of the two vector spaces, where, as an initial proxy, correlative measures (such as Pearson correlation) can be used. An initial, unmixed, state consisting of two fused chromosomes (via a Robertsonian fusion), each of size two, and each containing a single gene that can be found with equal probability at either of the two positions, can be represented as **a = (0.5,0.5,0,0)** and **b = (0,0,0.5,0.5)** and **a** ⊗ **b = (0, 0, 0.25, 0.25; 0, 0, 0.25, 0.25; 0, 0, 0, 0; 0, 0, 0, 0)**. Similar to the eigenvector decomposition, one can then use singular value decomposition of **a** ⊗ **b** to find that this matrix’ 1st singular value is explaining 100% of the matrix, suggesting that **a** and **b** can be decomposed and are therefore not yet mixed. However, if the two genes translocate within the newly formed fused chromosome, one possible resulting matrix is **(0, 0, 0, 0; 0.25, 0, 0.25, 0; 0.25, 0, 0, 0; 0.25, 0, 0, 0)**, and the 1st singular value will explain less than 100% of variance in the matrix indicating that a and b coordinates cannot be decomposed clearly any longer, reflecting mixing of the two ancestral states. Together, this suggests that fusion (without yet mixing) is represented by the tensor (or outer) product (⊗ or, **|a><b|**), whereas the mixing itself is moving the elements within this state matrix. Therefore, braiding (⤬, as defined in category theory) is mathematically a more appropriate representation of the mixing process. In our current implementation in this manuscript, we are using distance between orthologous genes as input for dimensionality reduction approaches and manifold projection. This allows us to compare evolutionarily very distant genomes, and focus on the variation in the genomic distance for the profiled set of orthogroups (n=2,361). In this context, the positional information of the example above is encoded by the orthogroup pair identifier. In the future, instead of profiling orthogroups, this approach will also allow to generalize to other genomic features, yet always requiring a high confidence underlying homology analysis.

In a multi-dimensional system represented by the tensor product of its components’ state vectors, a single mixing event (analogous to a chromosomal inversion or translocation) of the components along their coordinate systems is achieved by swapping values within the matrix. Each swap alone, without subsequent changes, is reversible (single braid). Irreversibility comes in when the topology-driven mixing results in a state where a and b identities cannot be distinguished anymore, without the knowledge of their separate identities from the outgroups. Thus, correlative approaches such as singular value decomposition, principal component analysis, and manifold approximation/projection are useful for the empirical identification of mixed states in genomic spaces. Additionally, singular value decomposition (or any other topological entropy measurements) can provide important insight into the ‘strength’ of mixing, with stronger mixing of the individual states being less likely to reverse.

